# Mammalian spermatogenesis at single-cell resolution downstream of genetic perturbations in meiotic double-strand break repair

**DOI:** 10.64898/2026.05.14.724910

**Authors:** I. Agarwal, B. Myers, M. Houlard, A. Hinch, E. Bitoun, S. R. Myers

## Abstract

Fertility in mammals relies on successful pairing of homologous chromosomes mediated by DNA double-strand breaks (DSBs). Here, we develop a system of mouse hybrids in which (a)symmetry of binding by the break-positioning protein PRDM9 to homologs varies over broad scales in the genome. Profiling transcription and chromatin in single nuclei, we trace how resulting delays in repair of variable subsets of meiotic DSBs propagate through spermatogenesis and drive large fertility differences in animals. We demonstrate that only asymmetry-generating mutations in PRDM9-binding motifs, not high average (∼1%) divergence, disrupt chromosomal pairing. We observe substantial variation in animal-level sensitivity to asymmetry, and identify an interacting locus containing *Dmc1* and *Mei1* controlling (R^2^=0.64) this variation. Silencing of unpaired autosomes downstream of asynapsis and failure of normal sex chromosome silencing independently explain cell death in pachytene. Surprisingly, many cells with synaptic defects evade cell cycle arrest, and even those where physical division arrests still exhibit *transcriptional* progression to post-division states. Attrition of abnormal cells via arrest continues beyond the first division; nonetheless, cells that complete both meiotic divisions exhibit aneuploidy, especially of the sex chromosomes. This partly reflects *de novo* segregation errors explained by silencing of only chromosomes 16 and 19. Even “normal” euploid spermatids show crossovers redistributed at multi-megabase scales, indicating novel and potentially post-zygotic impacts of delays in meiotic DSB-repair. We thus elucidate cell-level and chromosome-specific impacts of regulatory variation in ∼0.03% of the genome cascading through germline development, advancing our understanding of fertility and reproductive isolation.

## Introduction

The defining feature of sexual reproduction is a process of recombination between pairs of homologous chromosomes inherited from two parents, so that gametes carry unique combinations of parental genetic material. In mammals, at least one crossover event is thought to be required for proper segregation of chromosome pairs (*1*, *2*). The key steps in meiosis include a round of replication (so the 2n genome becomes 4n), followed by the formation of ∼300 double-strand breaks (DSBs) by SPO11 (*3–6*) at recombination hotspots positioned (with rare exceptions) by binding of the zinc-finger protein PRDM9 (*5*, *7–11*). DSBs are then involved in a process of homology search allowing proper chromosome synapsis during zygotene, and are directed during pachytene to be repaired as either non-crossover (∼90%) or crossover (∼10%) products (*12*). There follow two cell divisions, with the first (MI) separating homologous chromosomes at their centromeres, and the second (MII) then separating the remaining (sister) chromosomal copies to produce haploid gametes (*2*, *13*). A number of surveillance mechanisms (“checkpoints”) ensure the proper progression of these processes, with failure resulting in cell cycle arrest and eventual apoptosis (*14–16*).

Because of its role in positioning meiotic DSBs (*8–10*), perturbations of PRDM9 binding in the genome have provided a useful window into the early steps of recombination. PRDM9’s zinc fingers evolve rapidly, altering their binding affinity and transforming (fine-scale) recombination hotspot positioning genome-wide (*17–21*). Changes to the recombination landscape also arise from the evolutionary erosion of existing DNA motifs bound by a specific PRDM9 allele (*9*, *22*). Random mutations within binding motifs can disrupt binding leading to preferential (“asymmetric”) binding of only one of the two homologous chromosomes in heterozygous mutation carriers. Because PRDM9 recruits DSB formation in *cis*, breaks preferentially occur on the unmutated motif copy in heterozygotes, and the mutated motif is used as a template for repair: the resulting over-transmission of the mutated motif to offspring over many generations, leads ultimately to all individuals carrying the mutated motif (*23*) and turnover in preferred binding locations in the genome for the corresponding allele (*9*, *24–26*).

Alongside the recruitment of DSBs to recombination hotspots, binding of PRDM9 at the same location on the other, *uncut*, homolog has been shown to facilitate homology search and aid synapsis (*27–29*). At individual asymmetric sites (i.e., where one homologous copy remains unbound), DSBs show delayed repair (*29*, *30*) and generate lower levels of both crossover and non-crossover events, with evidence suggesting DSBs sometimes repair non-homologously at such sites (*27*, *28*). Hybrid mice with mainly asymmetric hotspots, generated by crossing the laboratory mouse strains PWD, B6, and CAST (representing inbred lines from the *musculus*, *domesticus,* and *castaneus* subspecies respectively, all possessing distinct *Prdm9* alleles) show incomplete synapsis and pachytene arrest in cytological spreads, accompanying infertility or reduced fertility at least in males: in fact, *Prdm9* is the only known mammalian speciation gene to date (*17*, *31–34*). In sterile PWD x B6 hybrids, for example, the PWD *Prdm9* allele has driven motif erosion on the PWD background and therefore preferentially binds to the B6 background (which has a distinct *Prdm9* allele and has therefore not suffered this erosion). The opposite occurs for the B6 allele, so that almost all recombination locations in PWD x B6 mice are asymmetric. Synapsis (and fertility) are rescued in these and other animals by mutating PRDM9 to symmetrically bind new locations in the genome (*29*, *30*). Further, introgression of B6 segments (of length as small as 20 million bases) into individual PWD chromosomes in PWD x B6 hybrids substantially rescues synapsis specifically of the introgression-containing chromosomes (*35*).

Despite this progress, prior studies have been unable to rule out a contribution of divergence between homologous chromosomes outside hotspots to asynapsis, and to elucidate the full spectrum of effects downstream of asynapsis. In particular, it remains unknown if there are any potential impacts on viable offspring in addition to those on fertility, via aneuploidy in surviving cells or shifts in broad-scale crossover landscapes, and how these impacts may vary by chromosome and/or variable amounts of genome-wide PRDM9-binding asymmetry. This is an important gap as many mouse crosses possess only partial PRDM9 asymmetry and remain able to produce offspring, and further this situation might be expected to occur frequently in nature given the evolutionary properties of PRDM9. Additionally, although it seems *a priori* likely that co-factors might interact with PRDM9 and mediate its impacts, the only known such case to date is the X-linked *Hstx* locus (*36–39*). A number of questions also remain about the relationship between meiotic silencing on the X chromosome and unsynapsed autosomes and how they each contribute to the pachytene checkpoint.

Here, we employ a mammalian system that randomly perturbs PRDM9 binding symmetry at broad scales along the genome, without requiring disruption of any particular gene and leaving overall heterozygosity constant, to address these questions. Specifically, with the aim that these mice might show a spectrum of fertility, we study male PWD x (B6 x CAST) F1/2 hybrids with the “infertile” PWD/B6 *Prdm9* allelic combination, where the maternally inherited homologous chromosomes are PWD and paternal homologs carry a random mix of B6 and CAST DNA. Divergence between any pair of the three mouse subspecies we examine is almost identical at ∼0.8% (*25*), only slightly lower than that between humans and chimpanzees (∼1%; (*40*)), so overall heterozygosity in these mice is high and fairly constant along the genome. However, *erosion* at PRDM9 binding motifs is expected to vary strongly by background. The PWD and B6 *Prdm9* alleles are likely to be eroded on their respective backgrounds, but not on the other two backgrounds, so we expect PRDM9 to bind more symmetrically where the paternal homolog is CAST rather than B6. Individual mice in our system therefore have varying patterns of PRDM9 binding asymmetry both animal-wide and on individual chromosomes, genetically perturbing (only) the earliest steps of DSB repair, namely homology search and synapsis.

For each mouse, we quantify fertility measures as well as gene expression and chromatin accessibility from single cells throughout spermatogenesis, leveraging hybrid ancestry to track recombination and cell division independently of expression-based cell type assignments. Analyzing these data using both existing and novel statistical approaches, we show that this system provides a powerful model to study mammalian spermatogenesis, allowing simultaneous assessment of transcriptional silencing by chromosome, the activity of meiotic checkpoints, the recombination landscape, rates of aneuploidy, and fertility outcomes. By carefully examining the downstream impacts of stochastic synapsis failure on one or more chromosomes in single cells, we show that PRDM9-driven asynapsis results in far-reaching (and sometimes chromosome-dependent) disruptions to all stages of meiosis, driving fertility reductions, accumulation of cell populations with decoupled cell division and expression programs, increased rates of potentially transmittable aneuploidy, and, despite previous evidence suggesting this is highly robust (*19*, *21*, *41*), strong broad-scale disruption of the crossover landscape.

## Results

### Data and fertility measurements

We generated and processed male germline tissue from 32 mice (see Methods, Supplementary Data). We sampled two parental strains, B6 and CAST (n=4, and n=1, respectively), and 27 hybrids, including B6 x CAST mice (with the human *Prdm9* allele; n=2), PWD x B6 mice (with B6 and PWD *Prdm9* alleles; n=2), and 23 PWD x (B6 x CAST) mice (with B6 and PWD *Prdm9* alleles) (Fig. 1A). We observed substantial variation in testes weight and sperm count among these animals and in particular the PWD x (B6 x CAST) mice, ranging from normal fertility (as observed in the parental strains and B6 x CAST hybrids with human PRDM9) to sterility (as observed in PWD x B6 mice) (Fig. 1B). The PWD x (B6 x CAST) mice differ from PWD x B6 mice in the presence of varying stretches of CAST sequence along the genome (on average 45% of the paternal genome in our sample). Therefore, introducing this CAST sequence has widely varying impacts, from full to no rescue of fertility.

**Fig 1.**
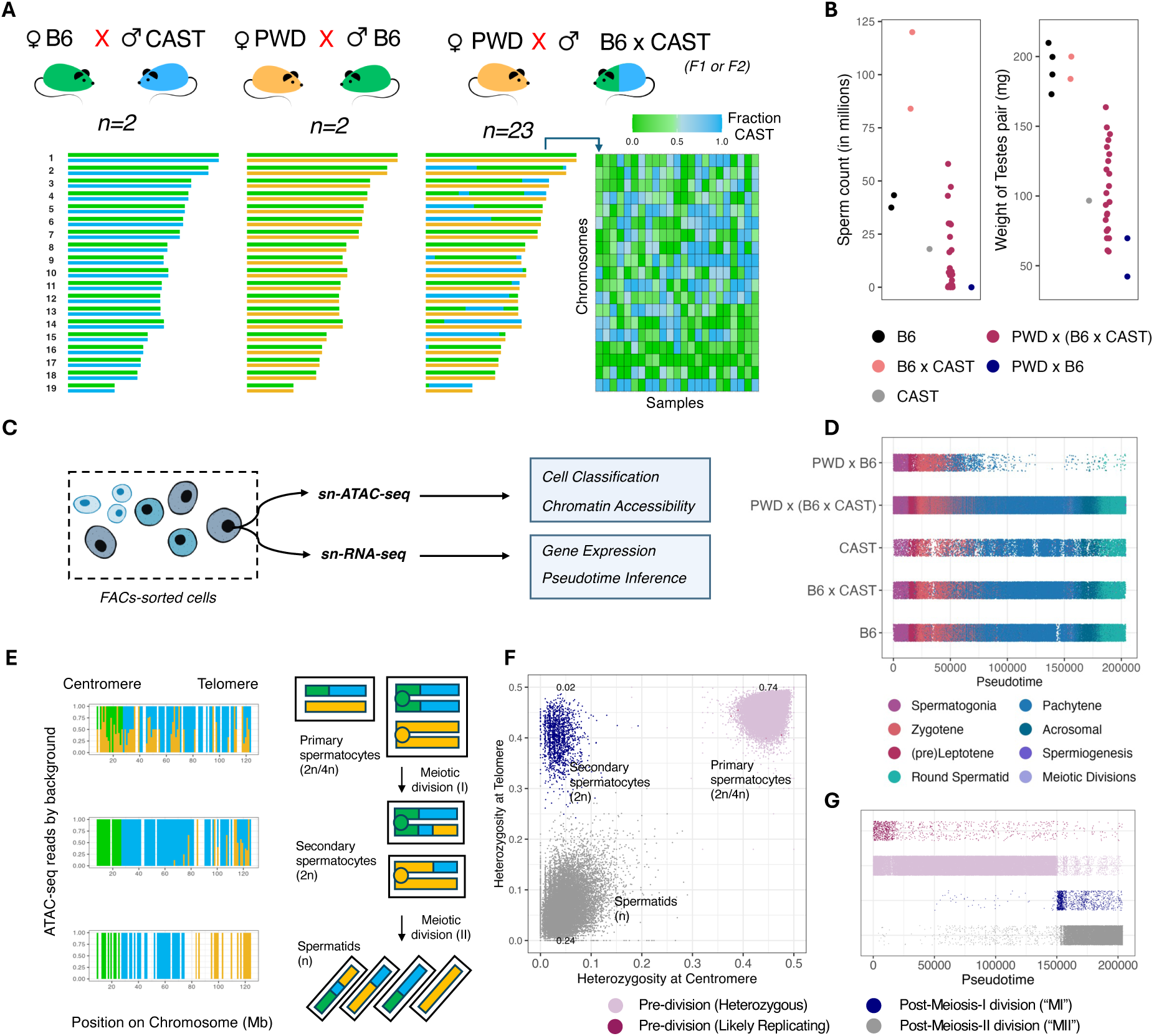
Overview of study. (A) Genome-wide ancestry patterns in the three types of mouse hybrids studied. PWD x (B6 x CAST) mice have a variable amount of B6 and CAST ancestry on the paternal homolog for each chromosome. PWD x (B6 x CAST) mice have the PWD and B6 *Prdm9* alleles, similar to PWD x B6 hybrids (see Methods). The B6 x CAST hybrids in this study carry the human *Prdm9* allele (*29*). (B) Fertility measurements (Testes weight and Sperm count) by animal and genotype (Methods). (C) A schematic representation of the single-cell study design, with ATAC-seq and RNA-seq data obtained from individual sorted cell nuclei from mouse testes using 10x Chromium Single Cell Multiome ATAC + Gene Expression protocols (see Methods). (D) Cell types represented in the study, ordered by the inferred Pseudotime ranks for individual cells (see Methods), shown separately for each genotype. Points (representing individual cells) are vertically jittered to aid visualization. Cell type identification is done by comparing gene expression in our study (see Methods) with a reference set of cell type informative gene expression patterns in a previous single-cell study of mouse spermatogenesis (*42*). The procedure for Pseudotime inference is described in Methods. (E) The distribution of ATAC-seq reads, colored by the strain of origin (PWD, B6, or CAST) along one diploid chromosome-pair in three exemplar meiotic cells from a PWD x (B6 x CAST) mouse. The schematic depicts the genetic background on the homologous chromosome pair as meiosis progresses, where chromosomes are heterozygous along their entire length before they undergo recombination and are partially (at the centromere) or fully homozygous after the first and second meiotic divisions, respectively. (F) Observed centromeric and telomeric heterozygosity by cell (averaged across chromosomes within a cell), in B6 x CAST, PWD x B6, and PWD x (B6 x CAST) hybrids. Cells are colored by our ATAC-seq based classification into the main meiotic cell types; only cells with normal chromosome types are included (see Methods). (G) Meiotic cell type designations (as in panel F) through Pseudotime. Points (representing individual cells) are vertically jittered to aid visualization.

To investigate the mechanisms that underlie this variation in fertility, we obtained data on gene expression and chromatin openness in single cells (using the 10x Genomics multiome assay, with RNA-seq and ATAC-seq reads from the same nucleus; Fig. 1C) throughout spermatogenesis (Methods). Our experimental approach involved FACS sorting of nuclei to enrich for short-lived germ cell populations in early/mid meiosis (Methods). We note that this pipeline obtains RNA-seq reads from the nucleus rather than the entire cell. This enriches for nascent transcripts and minimizes the potential presence and sequencing of cytoplasmic RNA copies originating in other cells (*42*, *43*) due to large numbers of inter-cytoplasmic connections arising from incomplete prior cellular divisions. We performed standard quality control procedures to filter poor quality cells and reads (Methods).

### Expression programs and ancestry-stratified ATAC-seq data independently identify meiotic cell types

We obtained 203,869 cells for further study, with a median 10,951 ATAC fragments and 1,394 UMIs (from 1,019 genes) per cell.

To examine the cell type composition of our sample, we used our gene expression data (at 15,000 genes) in individual cells to calculate weights for a published set of expression components capturing major spermatogenic programs (*42*), and assign an initial cell type based on the dominant expression component (Methods). Because meiosis represents a developmental process with fairly continual progression within and between cell stages (e.g., from leptotene, through zygotene, and into pachytene), we developed an approach to infer a set of time-varying gene expression components and pseudotime from our data (Methods). Starting with an initial ordering of cells based on the expression components in (*42*), we used a Gibbs sampling algorithm with Dirichlet priors to obtain a set of 75 gene expression components with distinct profiles through time (Methods, Supplementary Fig. 1), and obtained loadings on these time profiles for each cell given the genes it expresses. We used these cell weights to then define a smooth time trajectory and positioned each cell in this trajectory using linear interpolation to obtain a set of final cell ranks that we used for all subsequent analyses (Methods). We observe that the ranks are concordant with inferred cell types (Fig. 1D), with selected marker genes expressed at the expected times across genotypes and in the correct time order (Supplementary Figs. 2-3). These initial analyses suggest that our sampled cells capture the major germ cell populations present through spermatogenesis for each genetic background studied (Fig. 1D; note that sterile PWD x B6 hybrids possess few cells progressing beyond early pachytene).

ATAC-seq is a genome-wide assay of chromatin accessibility, identifying sites where the Tn5 transposase cuts DNA (*44*, *45*). We leveraged the high genetic diversity in our mice to first assign the strain of origin (PWD, B6, or CAST) to ATAC-seq reads from each cell: 50% of reads overlap one of ∼30 million ancestry-informative SNPs in the genome and therefore carry some information on background (Methods). Based on the likelihood of each read coming from each background and combining all data for an animal (Methods), we were able to quantify the relative proportions of the different ancestries by animal and chromosome, and identify ancestry segments and their boundaries (i.e., where paternal ancestry switches between B6 and CAST) in PWD x (B6 x CAST) hybrids (Supplementary Fig. 4).

We then analyzed ATAC-seq data in individual cells to infer their genomic state using a Hidden Markov Model (see Methods; Fig. 1E-G, Supplementary Fig. 5). Because they span meiosis, cells vary in chromosomal copy number (n, 2n, or 4n) with characteristic patterns of ancestry on homologous chromosomes (Fig. 1E). Combining information across chromosomes, we were able to separate cells that have not undergone a meiotic division (including a subset likely undergoing active replication), those that have undergone one division, and those that have undergone both meiotic divisions and are haploid (Figs. 1F-G). Reassuringly, these cell state inferences, which were independent of our pseudotime inference (based on gene expression patterns), yielded results strongly consistent with the order of events in spermatogenesis (Fig. 1G). These pseudotime and cell stage inferences are used extensively in our subsequent analyses.

### PRDM9 binding at recombination hotspots in early meiosis lags gene expression, and has variable levels of asymmetry by background

Following cellular commitment to meiosis, the expression of *Prdm9* is a key early event in the initiation of recombination. Consistent with previous findings (*42*, *46*)*, Prdm9* shows a strong peak of (nascent) expression in our data in a pseudotime interval likely corresponding to early leptotene (Fig. 2A, Supplementary Fig. 2).

**Fig. 2.**
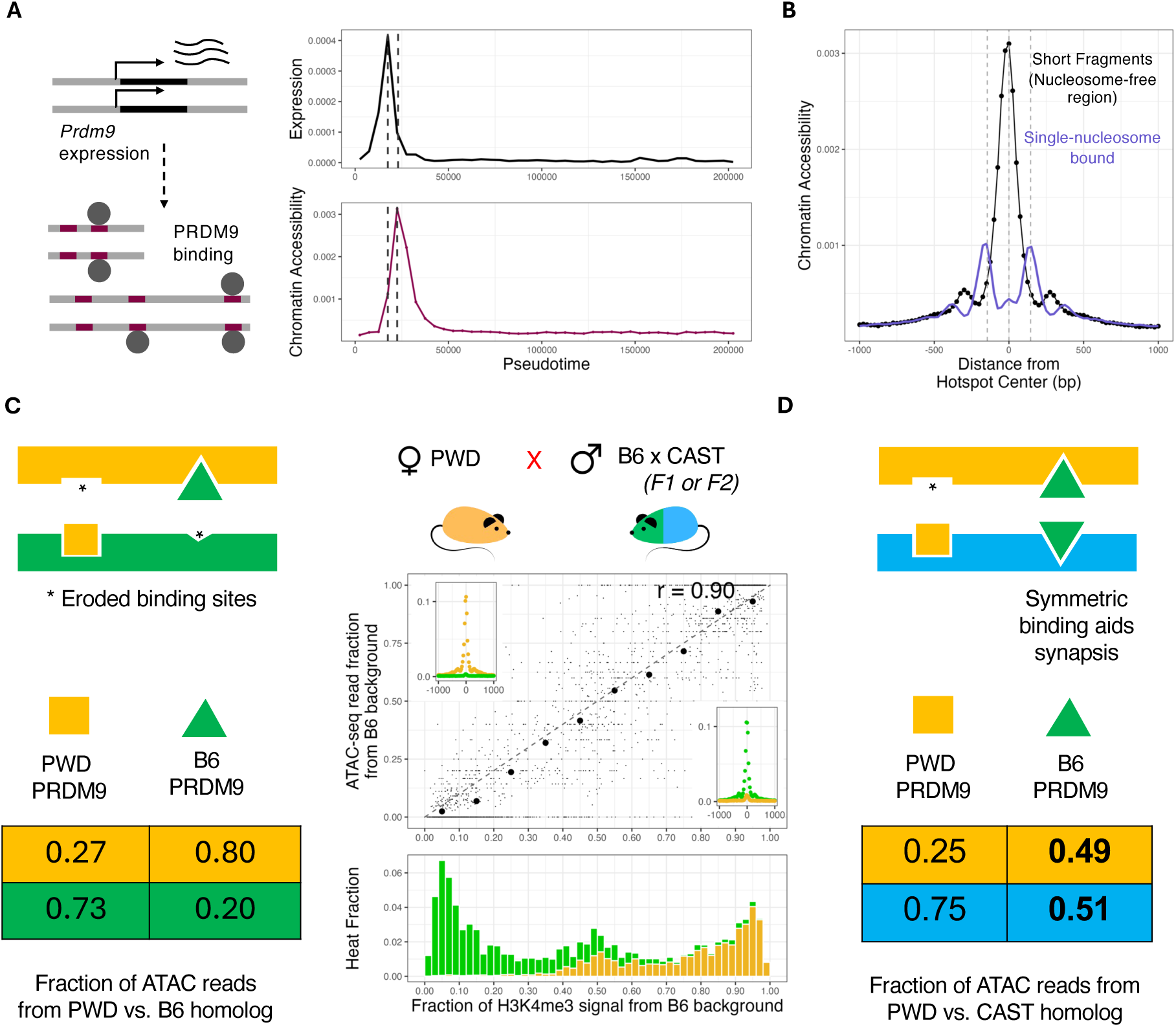
*Prdm9* expression and binding to recombination hotspots in early meiosis. (A) *Prdm9* expression relative to total gene expression in the same cells (top), and chromatin accessibility at recombination hotspots, defined as the proportion of sub-nucleosomal (“short”) ATAC fragments (<120 bp) genome-wide in 25 bp hotspot centers (bottom), through Pseudotime (see Methods). The two vertical dotted lines indicate the timing of the peaks of expression and binding, respectively. (B) Local chromatin organization around the average hotspot (center ± 1 Kb) at the peak of PRDM9 binding (in panel A). The y-axis is the proportion of sub-nucleosomal and nucleosomal fragments (of genome-wide counts by fragment type) in each 25 bp window in the selected Pseudotime bin, with each Pseudotime bin corresponding to 5000 cells (see Methods). (C) Asymmetric PRDM9-binding on the B6 and PWD homologous chromosomes in the PWD/B6 background. The table shows the average proportion (weighted by hotspot heats) of transposase cutting on the PWD vs. B6 homologous chromosomes at hotspot centers (25 bp), in hotspots that bind the PWD or B6 PRDM9 proteins. The scatterplot shows for individual hotspots the relationship of the ATAC-seq read fraction from the B6 homolog to the strength of H3K4me3-binding on that homolog (from Davies et al. 2016). Insets show the relative proportion of binding on each homolog at hotspot centers in hotspots with H3K4me3 signal from the B6 background < 10% (left) and > 90% (right). The stacked histogram shows the distribution of hotspot heats (calculated as the proportion of subnucleosomal ATAC fragments in each hotspot), stratified by the PRDM9 allele bound by the hotspot. (D) PRDM9-binding on the CAST and PWD homologous chromosomes. The table shows the average proportion (weighted by hotspot heats) of transposase cutting on the PWD vs. CAST homologous chromosomes at hotspot centers (25 bp), in hotspots that bind the PWD or B6 PRDM9 proteins.

We analyzed the binding of PRDM9 at 12,175 previously identified recombination hotspots controlled by PWD or B6 *Prdm9* alleles in PWD x B6 males, with identified (>95% posterior probability) motifs (*29*). Examining the genome-wide proportion of short ATAC-seq fragments (<120bp) at PRDM9 binding motifs, in cells binned by Pseudotime (see Methods), we observe a highly temporally localised opening of chromatin at these sites (Fig. 2A). The distribution of short and nucleosome-bound (150-250 bp) fragments by distance from the motif center (averaged across hotspots) reveals the expected signature of 2-3 nucleosomes flanking the open binding site (Fig. 2B) (*47*, *48*). The temporal peak of the DNA-binding signal occurs after the peak of transcription of *Prdm9* (Fig. 2A), reflecting time taken by transcript processing, translation, and reimport of PRDM9 into the nucleus.

To investigate allele-specific PRDM9-binding, we assigned (where possible) ATAC-seq fragments in PWD x (B6 x CAST) mice to a given homologous chromosome at each site, either “PWD” or “non-PWD”, with the latter further separated into B6 or CAST backgrounds. Using background-annotated short ATAC-seq fragments, we next calculated homolog-specific chromatin accessibility for each hotspot in the two possible local backgrounds in PWD x (B6 x CAST) mice: PWD/B6, and PWD/CAST (Methods). Focusing first on PWD/B6 regions, we observe much higher accessibility on the PWD homolog (80%) than the B6 homolog (20%) at hotspots controlled by the B6-derived PRDM9 allele, at individual hotspots and on average (Fig. 2C). Conversely, our data imply the PWD-derived PRDM9 allele binds mainly (∼73%) the B6 genome in these regions (Fig. 2C, Supplementary Fig. 6). We see a strong average correlation (90%, despite some noise for individual hotspots; Fig. 2C) with published estimates of PRDM9 binding asymmetry at the same sites (measuring PRDM9-associated H3K4me3 deposition to the PWD vs. B6 background for each hotspot in PWD x B6 animals (*29*)). We interpret this as implying that both short ATAC-seq fragments, and H3K4me3 levels, indeed reflect homolog-specific binding by PRDM9 at each hotspot, and that this occurs very similarly in PWD x (B6 x CAST) and PWD x B6 animals at individual hotspots for the same local background.

PRDM9-binding in PWD/CAST regions reveals an almost identical asymmetry for the PWD-derived allele (∼75% binding to the CAST genome), but a strikingly different pattern for the B6-derived allele, which now binds equally (51%/49%) to each background (Fig. 2D, Supplementary Fig. 6), with modestly higher overall binding of PRDM9 within PWD/CAST segments vs PWD/B6 segments, of ∼1.26-fold. These results are explained by the absence of evolutionary erosion of B6 motifs on either homolog in PWD/CAST regions, which then bind PRDM9 symmetrically, as observed. We then predict that the stronger symmetric binding by PRDM9 within PWD/CAST segments might aid synapsis of individual CAST-containing chromosomes and improve fertility–to an as-yet unclear extent. Because there is no systematic difference in *overall* genetic diversity in PWD/CAST vs PWD/B6 segments (only in hotspots that cover just ∼1% of the genome, and mainly concentrated in mutations within PRDM9 binding motifs constituting just 0.03% of the genome; (*29*)), PRDM9 binding asymmetry represents the *sole* consistent difference between these backgrounds that could plausibly drive asynapsis in *cis*.

### Asynapsis on sex chromosomes is accompanied by strong transcriptional silencing and two temporally-varying chromatin signatures

Our data do not contain direct information about DSB formation and synapsis. Because unsynapsed chromatin undergoes transcriptional silencing (meiotic silencing of unsynapsed chromatin, or “MSUC”) in pachytene (*13*, *49*), however, we investigated whether we might detect chromosomal asynapsis using its impact on gene expression.

We first examined the proportion of gene expression from the X chromosome in individual cells. The sex chromosomes, which are only able to form crossovers in the small pseudo-autosomal region, remain unsynapsed across most of their length (*50*, *51*), and are transcriptionally silenced at the onset of pachytene in a process called Meiotic Sex Chromosome Inactivation (MSCI) (*49*, *50*, *52*, *53*), thought to be a special case of MSUC. We observe strong and transient X-chromosome silencing in all animals (Figure 3A, Supplementary Fig. 7), though with imperfect silencing in some cells in some animals (see below). Because the restoration of X chromosome expression coincides with the first meiotic division, for simplicity we henceforth refer to the window of X-silencing as “pachytene” with respect to MSUC/MSCI, although genes associated with diplotene are also expressed prior to the division (Supplementary Fig. 2).

**Fig 3.**
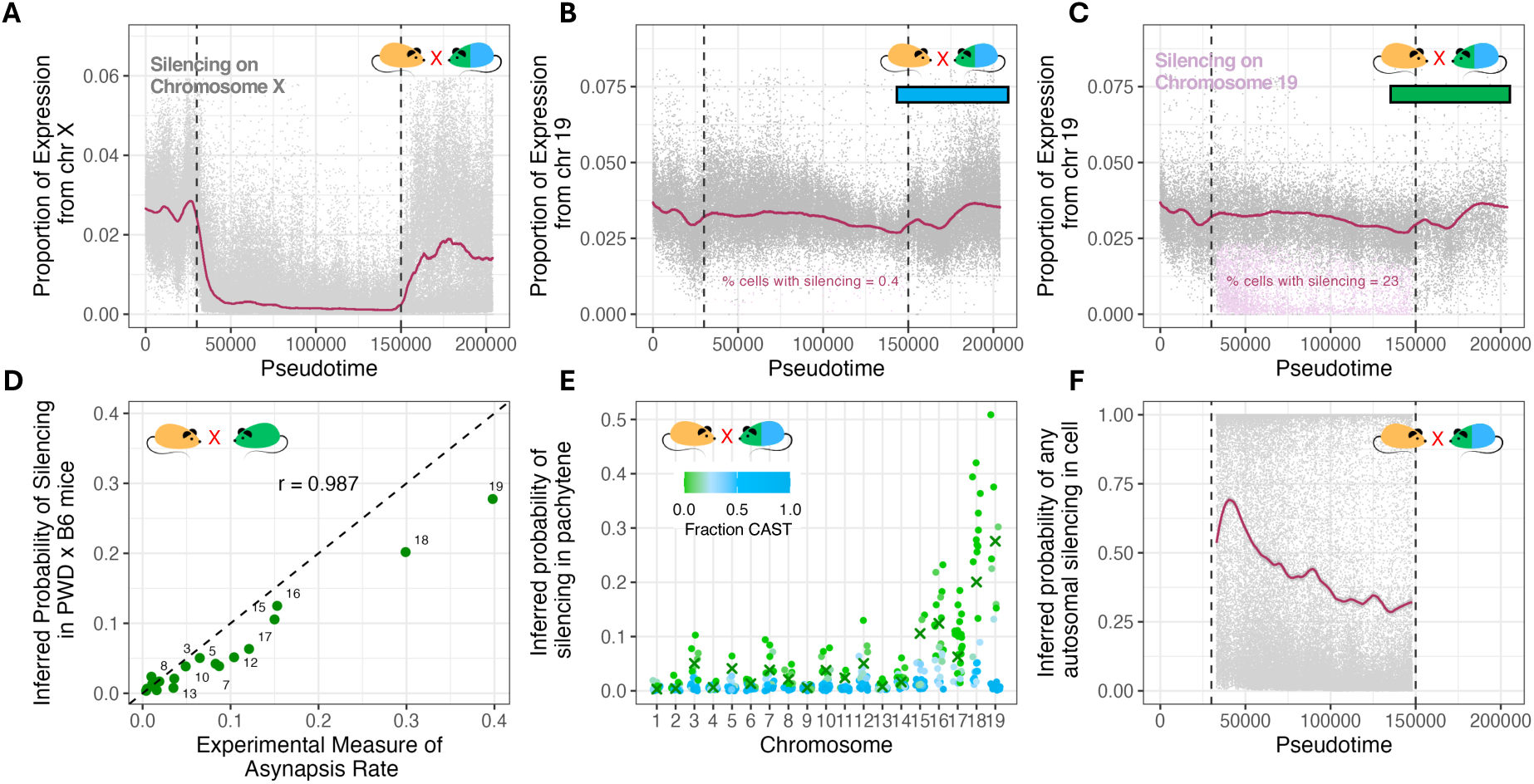
Transcriptional Silencing of the X and Autosomes. (A) The proportion of gene expression in single cells (in grey) from the X chromosome through Pseudotime in PWD x (B6 x CAST) mice. The maroon line indicates the median cell at a given Pseudotime. Vertical dashed lines indicate the Pseudotime interval corresponding to X chromosome silencing (MSCI). (B) The proportion of gene expression in single cells (in grey) from chromosome 19 through Pseudotime in mice with PWD/CAST background on chromosome 19. The maroon line indicates the median cell at a given Pseudotime. Vertical dashed lines indicate the Pseudotime interval corresponding to X chromosome silencing in panel (A). Cells with >50% silencing probability on chromosome 19 (see Methods) are highlighted in lilac. (C) The proportion of gene expression in single cells (in grey) from chromosome 19 through Pseudotime in mice with PWD/B6 background on chromosome 19. The maroon line indicates the median cell at a given Pseudotime. Vertical dashed lines indicate the Pseudotime interval corresponding to X chromosome silencing in panel (A). Cells with >50% silencing probability on chromosome 19 (see Methods) are highlighted in lilac. (D) Comparison of the average inferred silencing probability in PWD x B6 mice in our sample with experimental measurement of chromosome-specific asynapsis using FISH (*35*). (E) Inferred silencing probability by chromosome in PWD x (B6 x CAST) mice. Individual chromosomes are colored by the fraction of CAST (vs. B6) sequence. The average values for the two PWD x B6 mice are indicated as crosses. (F) The inferred probability of at least one autosome being silenced (see Methods) in single cells through Pseudotime, for all individual cells (denoted by grey points) in PWD x (B6 x CAST) mice. The maroon line indicates the LOESS fit through Pseudotime. Vertical dashed lines indicate the Pseudotime interval corresponding to X chromosome silencing in panel (A).

Comparing ATAC-seq fragment size distributions on the X and autosomes in B6 animals with properly synapsed autosomes (Supplementary Fig. 8A) through pseudotime, we observe a clear and temporally varying difference in this distribution between the X and autosomes that is specific to pachytene (Supplementary Fig. 8A), with two distinguishable components (Supplementary Figs. 8-9). The first is consistent with an X-specific increase in the proportion of normal nucleosome-bound fragments accompanying reduced chromatin accessibility, which co-occurs with the onset of transcriptional silencing, and the second with a pachytene-specific remodeling that presents as increased nucleosome size and/or spacing on the X, with a small increase in accessibility (Supplementary Fig. 8A-B). The latter “altered nucleosome” signature not only shows a strong mid-pachytene peak on the X, but also a noticeable smaller and earlier peak at the time of DSB formation on both the X and autosomes (Supplementary Fig. 8B-C). This pattern is consistent with activity of the DNA damage response (DDR) machinery, initially recruited to DSB sites genome-wide, but also required for the formation of the sex body and progressively accumulating on the sex chromosomes in mid-pachytene (*54–58*). Our data are too sparse to identify individual such sites, but the subtle altering of nucleosomes/nucleosome spacing observed could potentially reflect γH2AX accumulation around unrepaired DSBs, which then spreads over the sex chromosomes or recruits further histone modifications and protein-binding in mid-late pachytene (*59–61*).

### Quantifying chromosome-specific transcriptional silencing in individual cells identifies asynapsis rescue on chromosomes with symmetric PRDM9 binding

Given the above properties of MSCI, a similar reduction in expression (due to MSUC) is expected on unsynapsed autosomes. PWD x B6 animals indeed possess a distinct population of cells where chromosome 19 shows reduced (though rarely completely absent) expression, as do some PWD x (B6 x CAST) animals, notably those with a B6 chromosome 19 (Fig 3B, 3C, Supplementary Fig. 10).

Silencing of both homologs in individual cells is apparently symmetric and simultaneous, with no evidence of particular regions with stronger/weaker silencing (Supplementary Fig. 11).

We developed a statistical approach to quantify the signal of transcriptional silencing on autosomes, generating a model-based estimate of silencing using RNA-seq data (Methods). Taking as input the number of RNA-seq reads for each chromosome in each cell we estimate the fraction of pachytene cells with reduced expression (<75% of expected value) for each animal and chromosome. Our approach allows for statistical noise (i.e., varying numbers of reads per cell), variation through time in the fraction of reads from each chromosome (due to temporal changes in gene expression), and variation among cells and among animals (overdispersion, perhaps due to stochastic variability in gene expression in part due to different genetic backgrounds). We assume that the variation through time and variation across cells is shared among animals, and fit this using data from B6 animals, which properly synapse all autosomes. This approach does not use prior knowledge regarding individual or chromosomal genetic background, allowing us to compare the resulting estimates across animals. Consistent with normal synapsis, the average inferred silencing rate in the parental subspecies is <1% on all chromosomes and similarly low (≤1%) for B6 x CAST F1 animals (humanized at PRDM9) (Supplementary Fig. 10). We further validated the inferred silencing probabilities for each chromosome in the two PWD x B6 mice using previous (and independent) direct estimates of chromosome-specific asynapsis rates in PWD x B6 animals, obtained by manual counting of individual cells marked by chromosome-specific FISH (Figure 3D; (*35*)). There is a nearly perfect (98.7%) correspondence, implying that indeed transcriptional silencing is an excellent marker for autosomal asynapsis.

PWD x (B6 x CAST) hybrids show a wide range of inferred MSUC values, varying by chromosome and animal (Figure 3E, Supplementary Fig. 10B). Strikingly, for every chromosome we observe negligible silencing in animals possessing mainly CAST ancestry, but on shorter chromosomes in particular with mainly or completely B6 ancestry silencing is observed at rates comparable to those seen in PWD x B6 mice (Figure 3E). Given that the rate of transcriptional silencing mirrors that of asynapsis, this implies that in PWD x (B6 x CAST) animals, asynapsis impacts the same chromosomes as in PWD x B6 animals, but only where the background is PWD/B6, with synapsis fully rescued by the presence of CAST DNA segments. As we showed above, these segments possess more symmetric hotspots (Fig 2D), but not lower sequence divergence. Our results therefore demonstrate a powerful rescue of pairing in *cis* specifically by symmetric PRDM9 binding sites, as had been hypothesized to explain previous findings (*35*).

### The silencing response to autosomal asynapsis is incomplete relative to MSCI and does not disrupt it in the same cells

Animals with large amounts of synapsis failure on autosomes almost invariably also show failure of MSCI to some degree, though the precise mechanism by which this occurs and its timing remain unclear (*33*, *49*). Examination of X-chromosome expression in our data identified a subset of pachytene cells with aberrant X chromosome expression (at or above 75% of pre-pachytene levels), with the highest rate of failure in PWD x B6 animals (Supplementary Fig. 7). Notably, we do not find a meaningful correlation between silencing on autosomes and aberrant expression of the X within single cells in pachytene (ρ = -0.06, after a linear correction for the time trend), suggesting that unsynapsed autosomes do *not* directly compete with the X for silencing effector proteins in the same cells. Accordingly, autosomal silencing also does *not* meaningfully disrupt broad scale chromatin patterns on the X in the same cells (Supplementary Fig. 8D).

The severity of MSCI failure appears to decrease through early pachytene in our hybrids (Supplementary Fig. 7). The fraction of cells showing autosomal silencing also declines as pachytene progresses (Figure 3F), with chromosomes varying in the speed of decline of the silencing signal, not obviously related to their overall asynapsis rates (Supplementary Fig. 12). Given that failure to silence the sex chromosomes has been shown to drive apoptosis through the untimely expression of apoptosis-triggering genes including *Zfy1* and *Zfy2* (*62*), and since it does not strongly co-occur with silencing on autosomes, chromosome-specific declines in average silencing levels raise the possibility that misexpression of key autosomal genes, variably represented among chromosomes, also drives apoptosis in pachytene.

We note that despite both initiating simultaneously (Fig 3A-C), silencing even on the most asynapsis-prone autosomes is never as pervasive as (analogously defined, at < 75% of pre-pachytene levels) silencing on the unsynapsed X chromosome. We investigated whether transcriptional silencing on autosomes might be accompanied by the same histone modifications as observed on the X chromosome in the same cells (Supplementary Fig. 8D). Consistent with the strong molecular connection between MSCI and MSUC, silenced chromosomes show an accumulation of altered histones, and conversely this is only observed in cells and animals with expression silencing (Supplementary Fig. 8D, Supplementary Fig. 13). Supporting our analysis of consistently incomplete silencing, however, only partial (appearing variable among cells; Supplementary Fig. 13) incorporation of altered nucleosomes occurs on autosomes, implying that an incomplete analog of MSCI occurs for unsynapsed autosomes.

### A 20 Mb region containing *Dmc1 and Mei1* interacts with PRDM9 binding symmetry to control silencing/asynapsis levels across animals

Perhaps the simplest possible model of synapsis rescue by symmetric hotspots is that each DSB in a CAST region of the genome provides a constant probability of allowing synapsis of the whole chromosome, so that the probability of asynapsis declines exponentially from the value observed in PWD x B6 animals with the length of CAST DNA on a chromosome. We fit the exponential decay rate from the data using all chromosomes and animals jointly (Methods). Our model implies a ∼50% reduction in asynapsis for each 25 Mb of CAST DNA, on chromosomes with otherwise B6 DNA. Though our mice have much higher levels of divergence between homologs (*25*), this parameter is fairly similar to observations of the impact of introgressed B6 segments on the same PWD x B6 background (*35*), as expected if divergence outside hotspots does not contribute to asynapsis.

We compare the fit of this model for individual chromosomes with observations of transcriptional silencing, again using the latter as a surrogate for asynapsis. The model provides a good fit, explaining ∼75% of the variance in asynapsis levels across chromosomes (Fig 4A; r = 0.86, p < 2×10^-15^). However, the fit is imperfect: asynapsis-prone chromosomes lacking (< 2Mb total) CAST DNA show variation in silencing between animals around the PWD x B6 value (Fig. 3E, Fig. 4A). We estimated an animal-specific parameter that reflects overall levels of silencing relative to our model fit (Methods). Although estimated from the subset of chromosomes within each animal that lack CAST DNA, this measure captures a shared effect across chromosomes within an animal.

**Figure 4.**
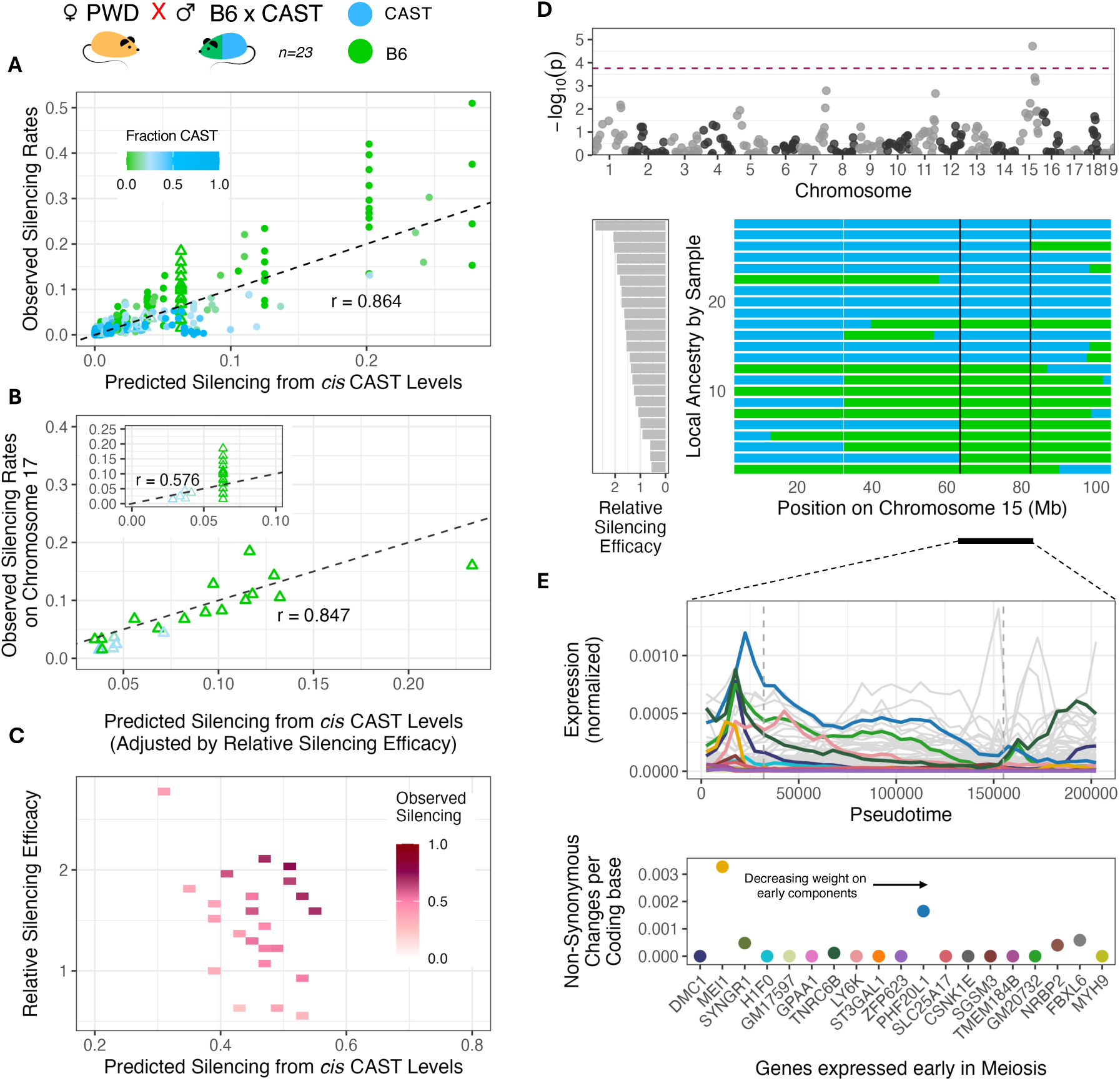
The genetic contributors to autosomal asynapsis in PWD x (B6 x CAST) mice. (A) The “observed” probability of silencing by chromosome and animal (inferred from gene expression data in single cells; see Methods) vs. silencing predicted from *cis* CAST levels by chromosome and animal, under a model of exponential decay with CAST sequence, from the value observed in PWD x B6 animals (see Methods). Points reflect individual chromosomes across animals and are colored by the proportion of CAST (vs. B6) sequence on a chromosome. Values corresponding to chromosome 17 are denoted by open triangles. (B) The observed probability of silencing for chromosome 17 by animal vs. silencing on chromosome 17 predicted from CAST levels on that chromosome and an animal-specific measure of silencing efficacy (see Methods). The inset shows the relationship between observed silencing for chromosome 17 and the predicted silencing levels from *cis* CAST levels from panel (A) for comparison. (C) The relationship of observed silencing with the underlying *cis* (on the x-axis) and *trans* predictors (on the y-axis). (D) Genome-wide association tests between the Relative Silencing Efficacy phenotype and B6 vs CAST background at 286 loci. The top panel shows all results genome-wide, with the horizontal dotted line indicating the genome-wide Bonferroni-corrected significance threshold (=0.05/286). The bottom panel shows the distribution of the phenotype (left) and of local ancestry on chromosome 15 by animal (right), with the most strongly associated segment (63.5 - 82.4 Mb) indicated with vertical black lines. (E) Potential candidates with an impact on animal-specific silencing efficacy within the 18.9 Mb highlighted region on chromosome 15. The top panel shows gene expression profiles through Pseudotime for the 181 genes in the segment with at least 500 reads across all 23 animals combined (in grey). Vertical dashed lines (in grey) indicate the Pseudotime interval corresponding to X chromosome silencing (MSCI). Genes with more than 10% of their weight in early expression components (see Methods) are highlighted as likely candidates. For highlighted genes, the bottom panel indicates the number of non-synonymous changes observed per coding base in the CAST (vs. B6) background.

We demonstrate this property using chromosome 17, which is 100% B6 in many animals in our sample (because animals were chosen to possess the B6 *Prdm9* allele on chromosome 17), but still shows a substantial amount of variation in silencing between animals when B6 (Figure 4B inset). The animal-specific silencing parameter, when calculated excluding chromosome 17, accurately predicts chromosome 17 silencing even for those animals with 100% B6 (Fig 4B, r = 0.79, p <3×10^-4^), despite their identical genetic backgrounds in *cis* for this chromosome, and likely due to residual variability in asynapsis among animals lacking CAST DNA on this chromosome. The silencing differences captured by this measure (which we refer to as “relative silencing efficacy”) are then *trans*-acting across chromosomes and cells within an animal.

At the level of animals (vs. individual chromosomes above), both our model of the *cis* impacts of CAST DNA, and the *trans*-acting relative silencing factor explain similar fractions of the variance in the probability of at least one silenced chromosome being observed (R^2^ = 0.8; p < 1×10^-7^ for the joint model fit; Fig 4C, Supplementary Table 1, Methods). The total proportion of B6 DNA in the animal, which reflects the overall genome-wide level of PRDM9-binding asymmetry by animal but is distinct from our autosomal model by not accounting for chromosomal differences in asynapsis, in contrast is not predictive (ρ = 0.26, p = 0.23; Supplementary Table 1).

Given it is animal-specific, we examined whether the relative silencing factor might be mappable. Treating it as a phenotype, we tested for association with the variable CAST/B6 presence along the genome. This revealed (Fig. 4D) a very strong association (r = 0.8; p = 1.9×10^-5^) with an 18.9 Mb locus on chromosome 15 (63.5 - 82.4 Mb), almost perfectly separating higher from lower relative silencing and remaining genome-wide significant after permutation testing (p=0.002; Methods). In this region (unlike the overall pattern), CAST DNA is associated with stronger silencing (and so likely higher asynapsis). Consistent with this, the two PWD x B6 sterile hybrids, though not used in our formal testing, fall within the group with lower relative silencing (their sterility is still expected, because they have a maximal level of asymmetric PRDM9 binding). The most strongly associated segment segregates identically across mice (Fig. 4D) and contains 309 genes (including 261 protein coding genes, and small numbers of miRNA, snRNA, and lincRNA genes), of which 181 have at least 500 reads across the 23 mice considered. Though we cannot identify the causal mutation, several genes expressed in early meiosis (Fig. 4E) represent excellent candidates: in particular the genes *Dmc1* (essential for homology search; (*63*)) and *Mei1* (required for DSB formation; (*64*)) both fall within the region. We do not observe obvious allele-specific expression differences for these genes (Supplementary Fig. 14), but *Mei1* possesses 13 non-synonymous coding changes between the B6 and CAST alleles, making it outlying within the region, and an especially strong candidate (Fig. 4E).

### PRDM9-binding asymmetry is a *cis* driver of MSUC but a *trans* driver of MSCI failure, both independently contributing to cell death in pachytene

Prior data have been unable to address the full set of downstream impacts of asynapsis, and how naturally occurring genetic variation beyond PRDM9 might impact these processes. Starting with genetic factors and leveraging the fact that earlier events in meiosis impact later ones, we predicted intermediate meiotic phenotypes and fertility, systematically performing conditional analysis to construct an objectively fit model of the impacts of various meiotic events on one another (Methods; Fig. 5A; Supplementary Tables 1-5).

**Fig 5.**
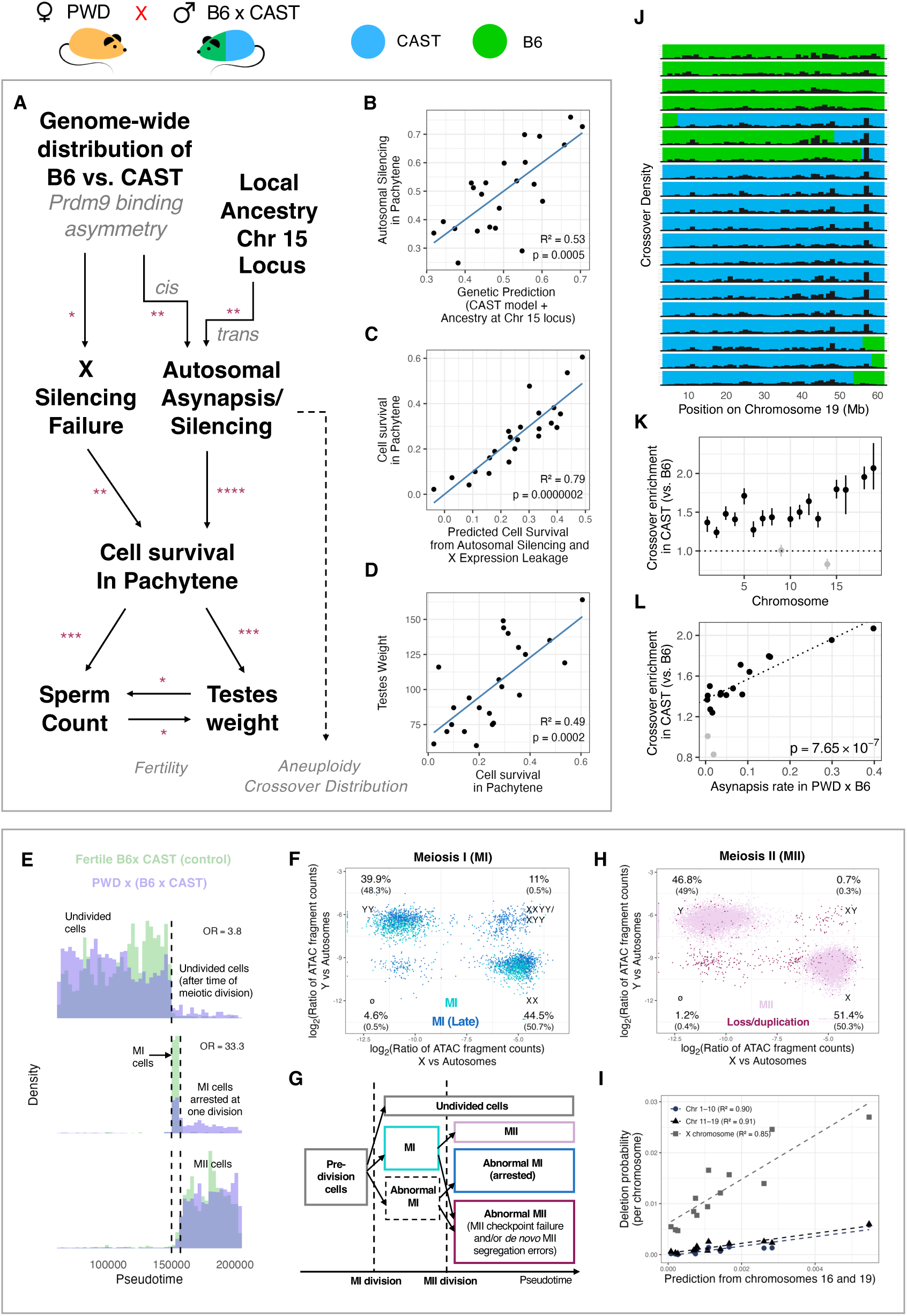
Downstream effects of PRDM9-binding asymmetry on fertility, aneuploidy rates, and crossover positioning in PWD x (B6 x CAST) male mice. (A) Predicting intermediate meiotic phenotypes and fertility from genetic factors (Supplementary Tables 1-5). Significant relationships are indicated by * (* p < 0.05, ** p < 0.005, *** p < 0.0005, **** p < 0.00005). (B) Predicting animal wide transcriptional silencing from *cis* and *trans*-acting genetic factors. The genetic prediction is from a joint model of local ancestry (B6 or CAST) on the Chromosome 15 locus identified earlier (see Fig. 4) and the animal wide silencing based on the CAST content of individual chromosomes. (C) Predicting cell survival in pachytene (calculated as the proportion of pre-division cells in late pachytene; see Methods) from the failure of X chromosome silencing and estimated autosomal silencing levels per animal. (D) Predicting the testes weight phenotype from the cell survival in pachytene measure calculated as in (C). (E) The distribution of meiotic cell types (Undivided, MI, and MII) through Pseudotime in PWD x (B6 x CAST) compared to fertile B6 x CAST animals (controls). Vertical dashed lines indicate the timing of the first and second meiotic divisions, respectively (see Methods). The odds ratios are calculated, for undivided cells as the ratio of PWD x (B6 x CAST) vs. B6 x CAST cell counts post-division relative to the ratio in late pachytene; for MI cells, cell counts are compared in the two genotypes in the time window between the first and second divisions and after both divisions (p-values are obtained using FET) (see Methods). (F) The X and Y chromosome content (determined from ATAC-seq data; see Methods and Supplementary Fig. 15) relative to autosomes of individual MI cells in PWD x (B6 x CAST) hybrids. The estimated proportions of the four types of cells (XX, YY, sex chromosome duplication, and sex chromosome deletion) are indicated, with the corresponding proportions in fertile B6 x CAST mice in parentheses. Cells are colored to indicate whether they belong to the early or late MI populations identified in panel (E). (G) A schematic representation of possible cell populations generated through the two meiotic divisions and associated checkpoint mechanisms, and their expected gene-expression based Pseudotime designation. (H) The X and Y chromosome content (determined from ATAC-seq data; see Methods) relative to autosomes of individual MII cells in PWD x (B6 x CAST) hybrids. The estimated proportions of the four types of haploid cells (X, Y, sex chromosome duplication, and sex chromosome deletion) are indicated, with the corresponding proportions in fertile B6 x CAST mice in parentheses. Cells are colored to indicate whether they carry detectable deletions or duplications of the sex chromosomes and/or autosomes or are otherwise normal (MII) in their chromosomal content. (I) The estimated per chromosome rate of homozygous deletions in MII cells (separately for chromosomes 1-10, 11-19, and the X chromosome) vs. the predicted values from a model of autosomal transcriptional silencing on chromosomes 16 and 19 (see Methods). Points indicate average values across MII cells within individual mice. (J) The distribution of crossover rates by genomic position on Chromosome 19 in PWD x (B6 x CAST) mice. The genetic background on this chromosome for each mouse is indicated in green (PWD/B6) and blue (PWD/CAST). (K) Enrichment of crossovers in the CAST (vs. B6) background (see Methods), averaged across cells in all animals, by chromosome. (L) The relationship between crossover enrichment in the CAST (vs. B6) background (see Methods) and a previously reported (*35*) measure of the asynapsis rate of chromosomes in PWD x B6 mice.

We began by predicting autosomal asynapsis/silencing: now in the more challenging setting of predicting properties (i.e., some chromosome failing) of animals, rather than synapsis of individual chromosomes. We use only genetic factors, namely, our model of the *cis* impact of CAST sequence by chromosome, and two *trans*-acting genetic factors: the novel chromosome 15 locus we identified which modulates the efficacy of silencing genome-wide, and the genome-wide proportion of B6 (vs CAST) DNA, which captures overall PRDM9-binding asymmetry by animal. Autosomal asynapsis and expression silencing remain well predicted (53%) by the first two measures, with no residual impact of genome-wide B6 level (Fig. 5B, Supplementary Table 1), confirming a dominant role for PRDM9 binding symmetry in *cis* in driving synapsis within this system. The chromosome 15 locus, containing key meiotic genes, rather than acting independently, appears to amplify the negative impacts of PRDM9 binding asymmetry when it is CAST (Supplementary Table 1).

In contrast to autosomal asynapsis, failure of sex-chromosome silencing shows only the genome-wide proportion of B6 ancestry as a significant predictor: we observe a significant correlation of the degree of X silencing failure with the overall proportion of B6 sequence in PWD x (B6 x CAST) animals (ρ = 0.61; p = 0.002; Supplementary Table 2). Not only do we not observe a direct relationship between autosomal silencing and X chromosome expression in the same cells, but at the animal-level, our autosomal silencing model also does not predict X-chromosome expression in pachytene (Supplementary Table 2), suggesting it is unlikely that autosomal silencing (even in nearby cells sharing transcripts, or with a time lag) directly mediates aberrant X-chromosome expression.

Rather than direct competition between silencing on X and autosomes, these observations favour a mechanism in which delayed DSB repair or synapsis at many asymmetric hotspots simultaneously, prior to the onset of silencing, leads to inappropriate autosomal retention of signaling and other proteins essential for early MSCI (*54–56*, *65*), reducing their availability to the sex chromosomes and causing MSCI to fail in some cells. Failure to silence X chromosomes, in turn, has been shown to cause cell loss in pachytene because of expression of apoptosis-triggering genes (*62*); whether autosomal factors independently contribute remains unknown.

Consistent with cell death in pachytene occurring as a consequence of misexpression of genes on both the X *and* autosomes (or alternatively an MSCI/MSUC triggered pachytene checkpoint mechanism), we find that cell survival in late pachytene (i.e., the proportion of pre-division cells in late pachytene; see Methods) is significantly reduced both by increased failure of X-chromosome silencing, and increasing autosomal asynapsis (and silencing), and extremely well predicted (79%) by the joint model (Fig. 5C; Supplementary Table 3). In our conditional analyses, the genetic factors show significant associations but mediate cell survival only through these intermediate silencing phenotypes (Supplementary Table 3).

For fertility measures further downstream (sperm count and testes weight), importantly, only cell survival in pachytene was a significant predictor, with all upstream factors conditionally non-significant (Fig. 5A, 5D, Supplementary Tables 4-5). This implies that the major driver of reduced fertility in our system is failure of cells to survive and reach the meiotic divisions. As we show, however, incomplete attrition of cells with synapsis defects likely further impacts gamete viability and/or quality.

### Persistent cell populations with arrested division show *transcriptional* progression through meiosis

Despite reduced fertility in our system, almost all animals have at least some cells that show gene expression patterns consistent with secondary spermatocytes and spermatids (Fig. 1D). We examined these cells for evidence of cell division failure, having previously leveraged genome-wide ATAC-seq data to independently identify cells that have undergone the first meiotic division (hereafter “MI” cells) and those that have undergone both divisions (“MII” cells) in PWD x (B6 x CAST) and B6 x CAST hybrids (Methods, Fig. 1E-G), and using stringent criteria to filter cells that may potentially be doublets.

Surprisingly, we observe (Fig. 5E) many cells in PWD x (B6 x CAST) animals that have gene expression programs aligned with late (post-division) meiotic cell types, yet have failed to divide (OR = 3.8 vs. B6 x CAST animals; p<<10^-16^; Methods). A number of cells in these animals also possess specifically MII-like gene expression but have only completed the MI division (Fig. 5E; OR=33 vs. B6 x CAST, p<<10^-16^). This implies that a number of cells fail to progress through cell division but nonetheless progress *transcriptionally* through meiosis: the average transcriptional state of a cell can thus successfully decouple from the cell division program. Together with a previous observation of decoupling of cell differentiation from “arrest” in pachytene (*66*), this raises the possibility that a further impact on fertility may arise from the accumulation of abnormal cells with arrested division that nonetheless may resemble spermatids, although such cells may still undergo apoptosis during spermiogenesis.

Another implication of these findings is that post-pachytene checkpoints (*67*) are active in these mice and flag substantial numbers of cells for division arrest. The pattern we observe is consistent both with a large proportion of cells with severe synapsis defects undergoing apoptosis during pachytene, and with the first division being relatively permissive to such defects in surviving cells in all animals: regardless, a non-negligible number of cells in PWD x (B6 x CAST) animals survive and divide once but fail to undergo the second division at much higher rates than in fertile mice without synapsis defects.

### Aneuploidy of sex chromosomes and asynapsis-prone autosomes arises from checkpoint failure and from *de novo* segregation errors at the second division

To assess whether synapsis failure leads to segregation defects in cells where at least one division occurred and to investigate drivers of aneuploidy, we examined cells for evidence of aneuploidy (0 or >2 copies in MI and 0 or >1 copy in MII) of the autosomes (Methods). We also quantified numbers of abnormal MI and MII cells possessing neither or both sex chromosomes (these should normally have either X or Y: in two copies for MI and one copy for MII cells).

Late-pseudotime cells which show evidence of arrest after the first division (Fig. 5E) show strong enrichment for homozygous deleted autosomes and sex chromosome abnormalities (OR = 5.9, p << 10^-5^, Supplementary Table 6), and are also enriched for potentially high-copy autosomes (Methods; Supplementary Table 6), implying that MII division failure often results from failure of MI to properly segregate chromosomes. Supporting this, of the over 15% of MI cells classified as having sex chromosome abnormalities (vs. 1% in fertile B6 x CAST hybrids), 89% are inferred to have been arrested at or before the second division (Fig. 5F-G). Accordingly, cells which *do* undergo the second division show much lower rates (2%) of X/Y abnormalities (though still ∼2x the estimated rate in fertile B6 x CAST mice) (Fig. 5H, Supplementary Fig. 15).

To identify the drivers of these chromosomal abnormalities, we tested by permutation (Methods) for correlation of the rates of inferred autosomal abnormalities with maximally asymmetric PRDM9 binding (i.e., an entirely PWD/B6 background) on the same chromosomes. Despite our modest MI cell counts (2,944 cells), probably due to the short time between divisions, and despite unknown attrition rates of arrested cells, asymmetric PRDM9-binding significantly increased rates of MI chromosomal abnormalities: autosomal deletions (complete deletions, OR = 2.08; p < 0.001) and duplications (Supplementary Table 7). Thus some MI segregation defects are readily explained by asynapsed chromosomes escaping pachytene arrest, potentially due to failure to undergo crossing over resulting in missegregation into daughter cells (*1*, *2*). Missegregation of X/Y occurs at very high rates but is not enriched in cells with autosomal abnormalities (OR = 0.83, p=0.46), and does not correlate with our asynapsis measures (beyond being specific to PWD x (B6 x CAST) animals), so we are uncertain as to the cause and whether it is influenced by *cis* (e.g., due to failure of pseudoautosomal recombination) or as yet unknown *trans* factors.

In turn, MII abnormalities could arise from escape of abnormal cells from MI arrest, or as *de novo* segregation errors at the second division (Fig. 5G). MII defects may lead to pre-zygotic sperm attrition, but they also have the potential to impact post-zygotic cells. Although both autosomal deletions and duplications were significantly (p < 10^-4^ for both comparisons) rarer in MII (2.45% and 0.82% of all autosomes, respectively) than in MI (3.26% vs 3.94% of autosomes), the decrease in the proportion of duplicated chromosomes is consistent with an estimated 80% rate of arrest in MI, while the proportion of deletions in MII is much higher than expected given a 90% rate of MI arrest.

Moreover, we observe a higher rate of duplications for chromosomes with asymmetric PRDM9 binding, but do not observe such a signal for deletions on the same chromosomes (Supplementary Table 7). These findings can be explained if both types of abnormalities cause MI arrest, some escape from arrest occurs at least for autosomal duplications, and there is an additional source of deletions for MII cells. Our proxy for arrest may be imperfect, but our observations suggest that there are likely substantial *de novo* segregation errors at the second division.

Intriguingly, PRDM9-binding asymmetry also appears to drive this signal, now via a different mechanism: in contrast to MI cells, we see a pattern of deletions in MII apparently driven in *trans* with shared drivers for both the autosomes and X/Y-chromosomes. Specifically, we observe very strong correlation in the observed rates of chromosomal deletions of the sex chromosomes and of autosomes (r = 0.83, p = 3.6 x 10^-4^), and also between longer (chromosomes 1-10) vs. shorter autosomes (11-19) (r = 0.96, p = 1.6 x 10^-7^), by animal, and a strong enrichment of co-occurring X/Y and autosomal deletions in the same cells (OR = 20, p < 2×10^-16^). X/Y-chromosome deletions are though more frequent (Fig. 5I), occurring at almost 9-fold higher rates than average autosomal rates. Although we observed a significant correlation (R^2^ = 0.62, p = 0.0014) between per-animal (pachytene) asynapsis rates and MII autosomal deletion rates, this correlation was significantly improved (R^2^ = 0.92 ; p = 2.16 x 10^-7^, Supplementary Table 8), and fully explained, by a model built from the asynapsis rates of just two asynapsis-prone chromosomes: 16 and 19, suggesting these as *trans* drivers. The same model predicted X/Y-chromosome deletions similarly well (R^2^ = 0.85, p = 7.3 x 10^-6^), despite not being fitted to those data. *Trans*-driven MII deletions therefore likely possess an MII-specific cause, e.g. failure to properly separate sister chromatids into distinct daughter cells. A possible mechanism is that silencing of key genes on chromosomes 16 and/or 19 in cells where these chromosomes show asynapsis earlier in meiosis compromises the ability of such cells, or nearby cells they share transcripts with, to properly undergo MII.

### (A)symmetric binding by PRDM9 reprograms crossover rates at multi-megabase scales

Beyond severe gametic defects, disruptions to early meiosis may alter the crossover landscape in otherwise normal gametes, ultimately impacting the parental contributions inherited by offspring. Previous work suggests that DSBs at individual asymmetric hotspots are less likely to homologously repair via the crossover or non-crossover pathways (*27*, *28*). However, PRDM9 binding has *not* been previously found to play a meaningful role in *broad*-scale recombination landscapes (*68*), which remain similar among different *Prdm9* alleles (*19*, *27*, *28*) and even different species (*21*), despite profoundly different fine-scale recombination hotspot positions. This robustness to finer scale perturbations has suggested that different factors regulate broad-scale and fine-scale rates, so we investigated if broad-scale recombination rates showed similar robustness in a setting with multi-megabase regions of PRDM9-binding asymmetry.

We mapped recombination crossover positions in individual MI and MII cells with normally segregated chromosomes in fertile or semi-fertile PWD x (B6 x CAST) and fertile B6 x CAST hybrids, analyzing our ATAC-seq data using a Hidden Markov Model (Methods, Supplementary Figures 16-17), yielding 245,368 crossover events in 24 mice (207,475 in 22 PWD x (B6 x CAST) mice). Overall patterns of crossing over in our animals appear broadly normal: 97% of chromosomes in MI cells have at least one detectable crossover in our sample, reflecting the expected requirement for at least one crossover, and 86% possess exactly 1 crossover. In MII cells across all mice, 46% of chromosomes were non-recombinants, and 48% showed exactly 1 crossover, suggesting strong crossover interference, again as expected. We observed on average 11.4 crossover events per *haploid* cell, consistent with reported estimates (*69*), with some variation across animals (Supplementary Fig. 16). The two B6 x CAST mice showed similar crossover landscapes (R=0.95; Supplementary Fig. 17), in contrast to individual PWD x (B6 x CAST) mice (Supplementary Fig. 17, Fig. 5J).

We next examined where crossovers occur *within* chromosomes in PWD x (B6 x CAST) mice (Fig. 5J): for each chromosome we constructed separate maps at the 2Mb scale for the two possible backgrounds: PWD/CAST and PWD/B6, to explore whether multi-megabase B6 segments, enriched for asymmetric hotspots, show altered crossover rates. By estimating “enrichment” of crossovers within PWD/CAST segments relative to PWD/B6 segments in mice with both backgrounds, separately for each chromosome (Methods), we obtained enrichment values > 1 for (symmetric-hotspot-enriched) CAST segments for all chromosomes except chromosomes 9 and 14 (Fig. 5K).

Excluding these, ratios >1 are seen for 97% of individual animal/chromosome combinations possessing 5-95% CAST ancestry (p < 0.05 in 70% of cases). The enrichment varies by chromosome, correlating nearly perfectly with asynapsis rates in PWD x B6 mice (r = 0.91; p = 7.7×10^-7^) and reaching ∼2-fold for chromosomes 18 and 19 (Figures 5K-L). Moreover, the two exceptions, chromosomes 9 and 14, are precisely those autosomes containing large introgressed segments of *domesticus* ancestry on the PWD background (*70*) restoring PRDM9 binding symmetry in PWD/B6 segments: so the observed *lack* of crossover enrichment within CAST segments for these chromosomes again matches predictions based on PRDM9 binding asymmetry. Thus, the binding properties (in particular, binding symmetry) of PRDM9 can alter the broad-scale recombination landscape at scales of many megabases.

What explains the stronger enrichment of crossovers within CAST segments on asynapsis-prone chromosomes, given that only normally synapsed chromosomes are expected to be able to perform crossing over, and synapsis is a chromosome-wide property in late pachytene? Because *Prdm9* expression is restricted to early meiosis, the observed relationship between crossover distributions and PRDM9 binding asymmetry indicates that the crossover/non-crossover decision is made early: after the onset of, but (at least in our PWD x (B6 x CAST) animals) prior to the completion of synapsis. If synapsis probabilistically initiates within regions containing symmetric hotspots, partial synapsis will most often occur first within CAST regions, only later spreading chromosome-wide. If crossover designation occurs in this key time period, when synapsis is normally completed in fully fertile animals but often incomplete for chromosomes that take longer to synapse, crossover sites would be strongly biased towards CAST regions, specifically on those chromosomes prone to asynapsis, as observed. If this model is correct, our results constrain the possible processes via which a subset of DSBs are directed into the crossover pathway, supporting both that synapsis initiates at symmetric hotspots as previously suggested (*29*), and that (only) local synapsis is required for crossover designation, with this decision made quite early. Experimental evidence also suggests this is plausible: the protein HEIP1 for instance, thought to be involved in crossover site designation, shows early foci on chromosomal axes prior to synapsis which relocate during synapsis to regions where homologs are in close proximity and to avoid remaining DMC1 foci (*71*), which will likely include slower-repairing asymmetric hotspots in our system.

## Discussion

Mutations that only disrupt symmetric binding of PRDM9 on homologous chromosomes and drive less efficient homology search and slower repair of meiotic DSBs at these sites (without requiring disruption of a key gene), produce severe and varied downstream impacts in the precisely regulated setting of spermatogenesis. Beyond reduced fertility, we see particularly surprising effects late in meiosis which potentially impact gamete and/or offspring fitness, including profound changes in broad-scale crossover rates, and persistent aneuploidy and segregation errors at the second meiotic division. Notably, despite a number of distinct proximal causes, the majority of these effects trace back specifically to asymmetry-generating genetic variation in PRDM9-binding motifs comprising ∼0.03% of the genome.

By exploiting broad-scale variation in PRDM9-binding asymmetry in our system, we powerfully demonstrate that synapsis is highly sensitive to (a)symmetry *in cis*, yet almost completely robust to sequence divergence outside hotspots. This supports a model where synapsis depends almost exclusively on homology recognition initiating at symmetrically bound DSB sites and then spreads chromosome-wide. An observed ∼50% reduction in asynapsis for each 25 Mb of CAST DNA on chromosomes with otherwise B6 DNA suggests, based on 300 DSBs genome-wide, that only 1-2 DSBs at symmetrically bound sites are likely sufficient for an entire chromosome to synapse. Also supporting our model, we observe striking multi-megabase-scale crossover perturbations, with crossovers redistributed into CAST DNA segments on asynapsis-prone chromosomes. PRDM9 binding is thus able to direct broad-scale recombination rates alongside (as previously known) fine-scale rates. This does not result from asynapsis *per se* (which would prevent crossing over) but instead is readily explained by crossover repair pathway choice occurring early in meiosis, designating DSBs in regions of the genome containing early-synapsing symmetric-hotspots as crossovers.

While PRDM9-binding asymmetry on autosomes drives asynapsis and meiotic silencing in *cis*, we find a novel *trans* impact on the X chromosome: higher overall asymmetric binding of PRDM9 and slower DSB repair genome-wide, rather than on particular chromosomes, appears to drive failure of proper silencing of the X chromosome in some cells, including where autosomes have synapsed normally. One implication is that even transient asymmetric binding of PRDM9 (as hotspots are lost by evolutionary drive) insufficient to actually cause asynapsis may nevertheless still have non-zero fitness costs due to this genome-wide effect, potentially helping to drive the observed rapid evolution of this gene not just across but also within all examined species possessing it.

PRDM9-binding asymmetry drives cell death and other downstream impacts through both its *cis* and *trans* effects. While failure of sex chromosome silencing has previously been shown to cause cell apoptosis in pachytene (*62*), we find that autosomal asynapsis independently predicts pachytene cell survival, and moreover, that the relationship with cell loss appears to vary by autosome. Together with evidence from oocytes suggesting silencing and not asynapsis itself as the main source of cell death in pachytene (*72*), our data supports a more general gene-centric view where consequences of silencing depend on specific genes affected, manifesting differently at different times depending on when they are essential (or lethal), and potentially affecting nearby cells sharing transcripts through cytoplasmic bridges.

We show that despite observable attrition of defective cells through pachytene and even beyond the first meiotic division, the effects of PRDM9-binding asymmetry nonetheless reach surviving gametes: crossover redistribution as well as aneuploidy, including of the sex chromosomes, persist past the second meiotic division in fertile or semi-fertile individuals. These defects arise again both from direct, *cis*-acting consequences of asynapsis on the affected chromosome, as in the case of crossover redistribution and synapsis failure causing missegregation of the same chromosome, and *trans*-acting errors driven by silencing-induced misexpression. Supporting the latter pathway, we find that silencing on just two chromosomes (16 and 19) strongly predicts de novo segregation errors on all autosomes and at especially high rates on the X chromosome at the second meiotic division far downstream of asynapsis: this is also consistent with the chromosome or gene-specific impact of silencing highlighted above. These previously unappreciated consequences of asynapsis are of particular interest because of their potential postzygotic impacts: while changes to the crossover landscape might only have subtle fitness effects, elevated aneuploidy (and in particular on sex chromosomes, because XO and XXY mice are viable), likely constitutes a real additional fitness cost of PRDM9-binding asymmetry on gametes and/or offspring.

A further surprising phenomenon evident from our analysis of single cells is that cell cycle arrest at both pachytene and post-pachytene checkpoints produces populations of cells in testes that still exhibit *transcriptional* (and potentially morphological) progression to post-divisional states despite failure to undergo one or both meiotic divisions. Although these cells may undergo apoptosis during spermiogenesis, given that they span the full range of downstream pseudotimes another possibility is that they persist as unviable gametes in testes and impose an additional (cryptic) cost on fertility.

Yet another unexpected finding is that transcriptional silencing on unsynapsed autosomes (MSUC) only partially resembles normal MSCI. While asynapsis therefore is one factor promoting MSCI in normal cells, our results imply that there are additional *cis* factors that reinforce this silencing by recruiting more or additional silencing machinery, for instance, which autosomes lack even if they fail to synapse. Such factors have also been suggested to underlie stronger *Xist*-mediated inactivation of the X chromosome relative to autosomes (*73*). These may have evolved on the X chromosome to ensure full silencing, and would be consistent for example with the sex chromosomes carrying genes that are especially harmful when expressed in pachytene in males. Thus, MSCI is not just a special case of MSUC at least in mammals, but has distinct properties and its own regulatory apparatus.

Beyond effects of perturbations to PRDM9 binding, we also observed a *trans* impact of a locus on chromosome 15, modulating silencing and/or asynapsis (as well as fertility measures). Alongside the previously known *Hstx* locus (*36–39*), this represents another region strongly impacting hybrid sterility and interacting with *Prdm9*, and fine-mapping of this locus should be a future research aim. Depending on the causal gene, the chromosome 15 locus might associate with numbers of DSBs (if *Mei1*): distinct from *Hstx*, we do *not* detect a clear impact of this locus on genome-wide crossover rates (p=0.078) despite variability in genome-wide crossover counts (measured in MII cells) across animals, and other modes of action (for instance, efficient recruitment or sequestering of factors involved in repairing DSBs as crossovers, facilitating synapsis, or altering the strength of the MSUC response to asynapsis) remain possible. Given there are at least two segregating large effect loci modulating sensitivity to PRDM9 binding asymmetry (and likely others exist) in male mice, and sensitivity to the negative impacts of PRDM9 binding asymmetry is also reduced in females (*33*, *72*, *74*, *75*), it may be possible for systems to evolve to be very insensitive to this property through a variety of developmental switches, and ultimately lose *Prdm9* function, as has been seen repeatedly across evolution (*11*, *76*).

Although we focus here on the fitness and phenotypic consequences of genetic perturbations to PRDM9 binding, our approach might be applied more widely to diagnose and understand other pathologies of germline development, including those involving faulty DSB-repair. It may also provide insights into germline mutation: that inefficiencies in repair at even a subset of ∼300 meiotic DSBs can cause high rates of cell apoptosis suggests that other types of DNA damage may sequester key DNA damage response proteins and temporarily impact fertility, providing a possible explanation for why the germline appears to be relatively protected from damage (*77*, *78*). Moreover, overlap with proteins involved in meiotic DSB repair might substantially increase selection against variants that impact the mutation rate, if these also impact synapsis or MSCI. Beyond germline-specific traits, this work elucidates one possible mechanism by which non-coding (as well as coding) variation, through subtle alterations individually of early processes and in particular their timing under developmental constraints, ultimately may have large cascading effects on complex phenotypes, and suggests that examining other such cases may aid our understanding of regulatory variation in human complex traits.

## Acknowledgements

We thank members of the Hinch and Myers groups for helpful discussions, and Molly Przeworski for comments on an earlier version of the manuscript. We thank Ruddy Montandon and the team at the Flow Cytometry Facility, Center for Human Genetics, University of Oxford, Andrew Worth at the Flow Cytometry Facility, The Jenner Institute, University of Oxford and Robert Hedley and Vasiliki Tsioligka at the Flow Cytometry Facility, Sir William Dunn School of Pathology, University of Oxford for providing technical assistance. Computation relied on the Oxford Biomedical Research Computing (BMRC) facility. This work was supported in part by a Wellcome Trust award 221761/Z/20/Z to AH and a Wellcome Trust Investigator Award 212284/Z/18/Z to SRM.

## Data availability

All raw and processed data generated as part of this study will be available on publication. Code used for analysis and to generate figures will be made available on GitHub at *agarwal-i/spermatogenesis_single_cell*.

## Supplementary Figures and Tables

**Supplementary Fig. 1.**
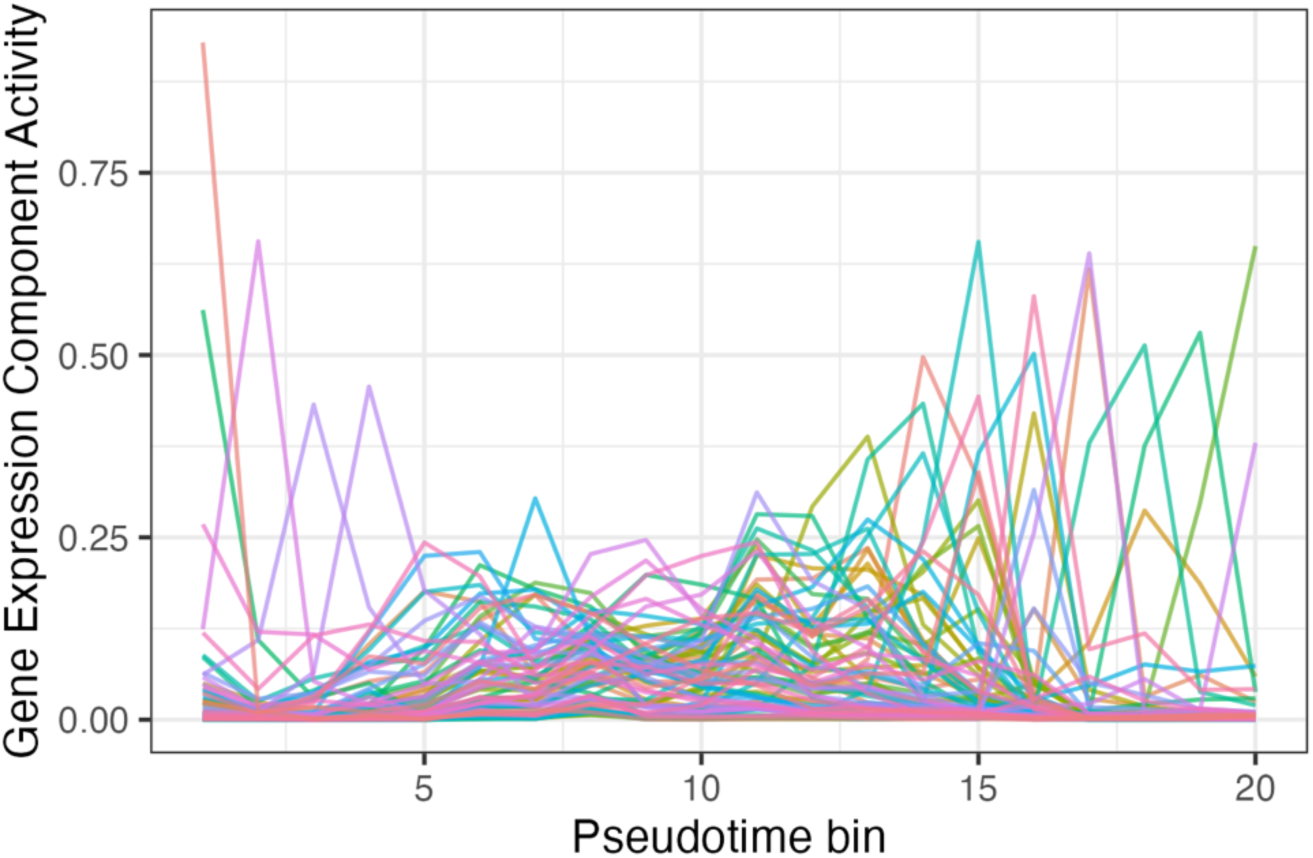
Pseudotime profiles of 75 Gene Expression components. Components were inferred from expression levels of 25,963 genes in 203,869 cells ranked by Pseudotime, in bins of 10,000 cells (see Methods).

**Supplementary Fig. 2.**
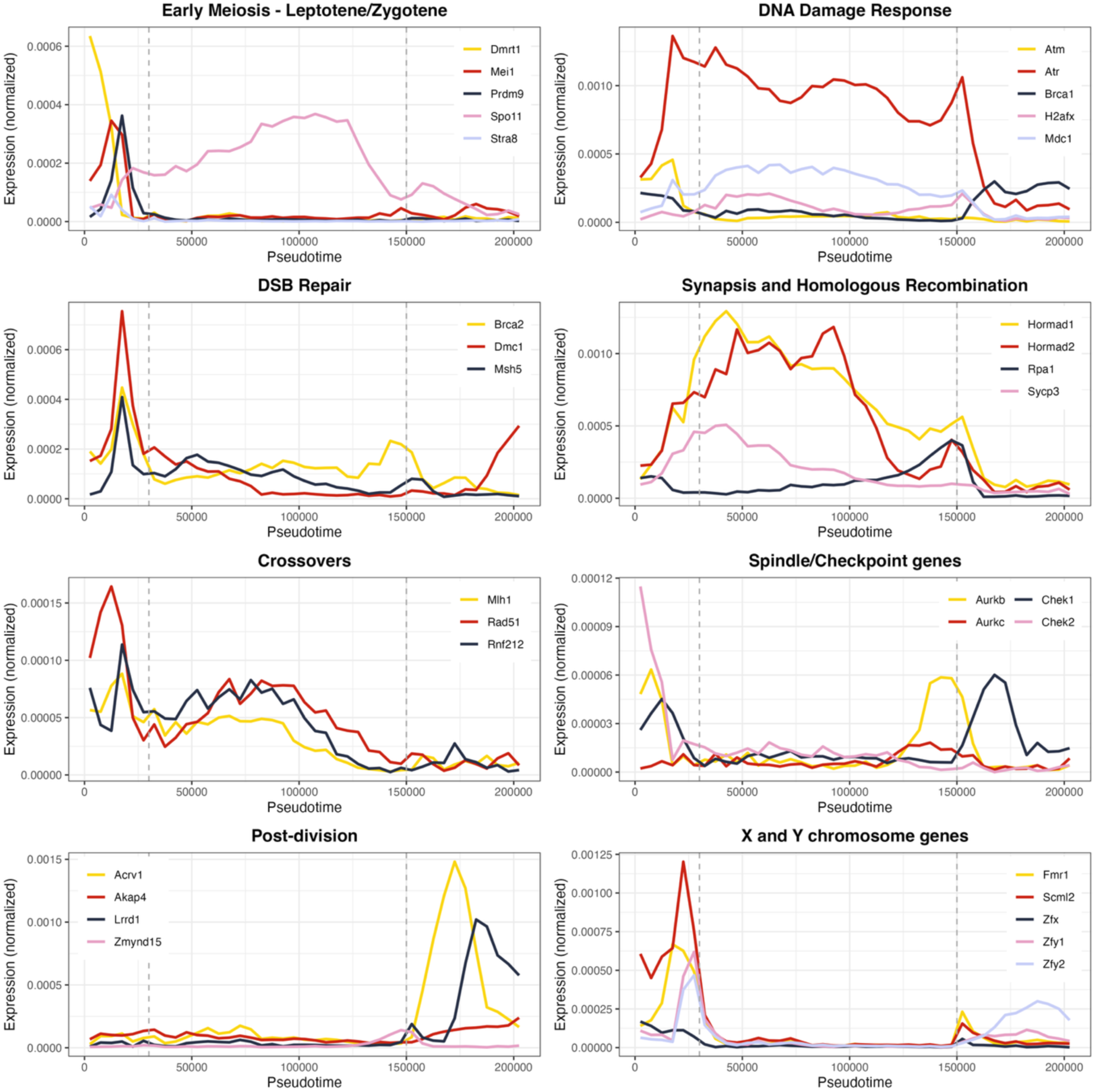
Gene expression through Pseudotime for key meiosis genes and other markers. (across all mice in our sample). Normalized gene expression for a given gene is calculated as the total number of reads from that gene divided by total gene expression, combining data from all cells in a given Pseudotime bin (each with 5000 cells). Vertical dashed lines indicate the Pseudotime interval corresponding to X chromosome silencing, which begins at the onset of Pachytene and recovers just prior to the first meiotic division.

**Supplementary Fig. 3.**
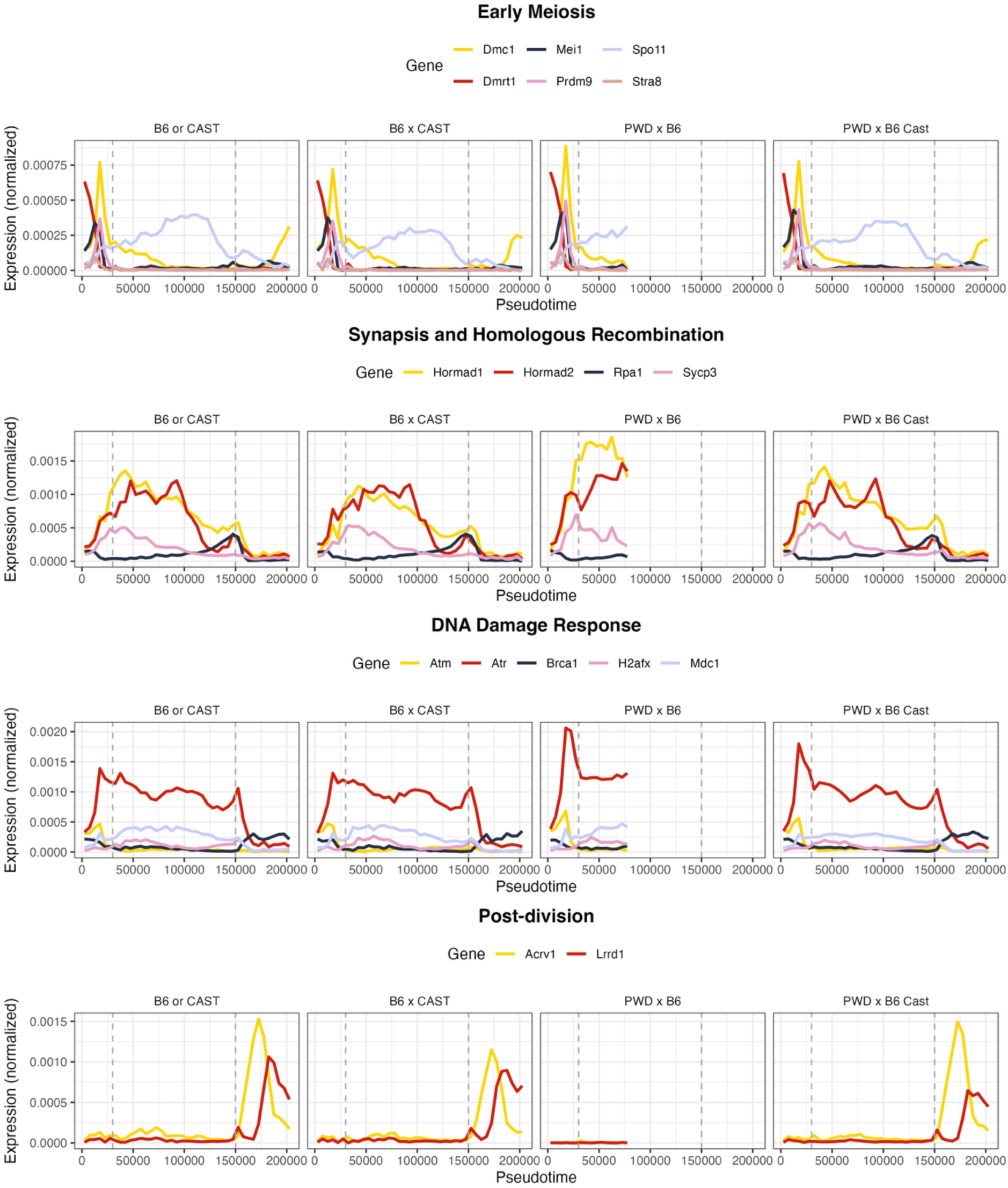
Gene expression through Pseudotime for key meiosis genes and other markers, by genetic background. Pseudotime was inferred jointly across samples. Normalized gene expression for a given gene and background is calculated as the total number of reads from that gene divided by total gene expression, combining data from all cells corresponding to samples with that genotype in each fixed Pseudotime bin of 5000 cells. Bins with fewer than 50 cells were removed. Vertical dashed lines indicate the Pseudotime interval corresponding to X chromosome silencing, which begins at the onset of Pachytene and recovers just prior to the first meiotic division.

**Supplementary Fig. 4.**
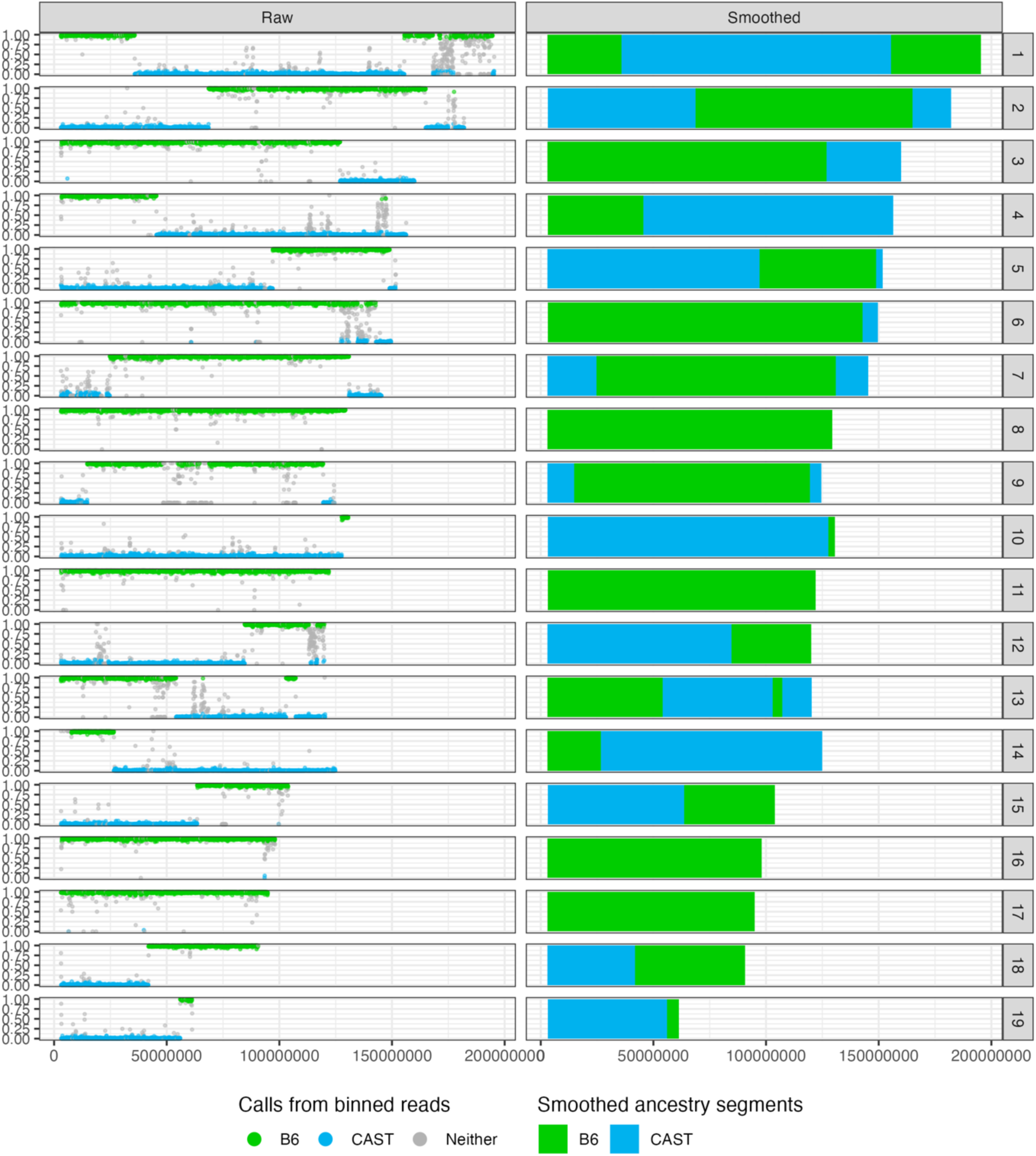
Local ancestry inference using ATAC-seq reads in PWD x (B6 x CAST F1/F2) mice. Colors indicate the background (B6 or CAST) on each paternal chromosome in one animal; maternal chromosomes have only the PWD background. The left panel shows binned reads (annotated with their allelic origin) in 50 kb bins with each facet indicating one chromosome, and the y-axis showing the proportion of reads in a bin with a B6 (vs. CAST) allele. Bins with >10 reads that were confidently assigned to one of the two backgrounds, with >90% reads from one of the backgrounds, were called as B6 or CAST; bins uncalled by this approach are shown in grey. Smoothed ancestry segments in the same animal obtained using a Hidden Markov model (see Methods) are shown in the right panel.

**Supplementary Fig. 5.**
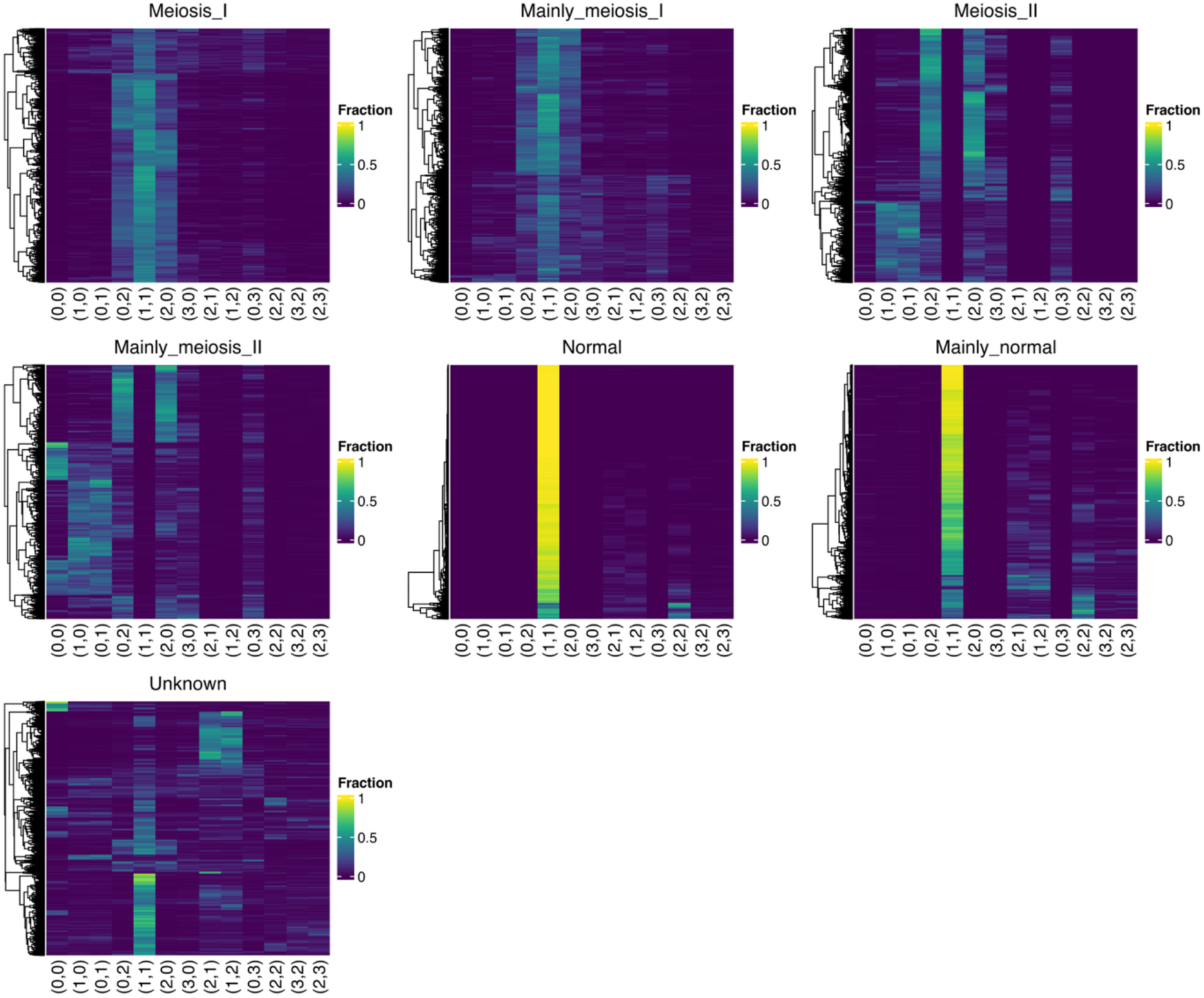
Genome-wide state composition by inferred meiotic cell type. Each row in a heatmap represents an individual cell of the corresponding cell type. The proportion of the genome in each of 13 possible states in each cell was inferred from ATAC-seq data using a Hidden Markov model (see Methods). States (shown on the x-axis) represent possible copy numbers for the two homologous chromosomes during spermatogenesis, with (0,0) representing a homozygous deletion, and (1,1) representing a normal diploid cell.

**Supplementary Fig. 6.**
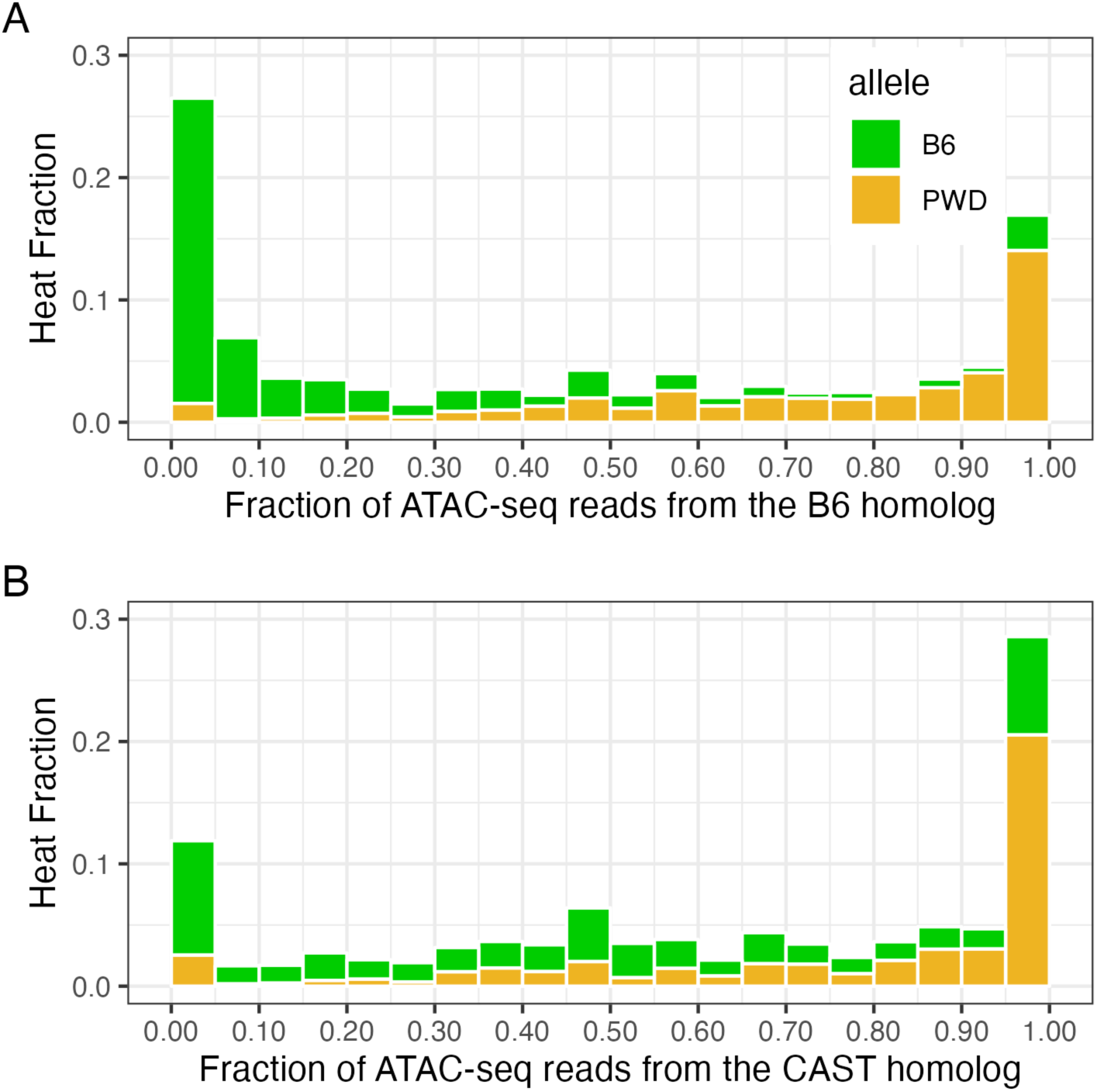
Hotspot heat distributions by allele and background on the non-PWD homolog in PWD x (B6 x CAST) mice. (A) The x-axis shows an ATAC-seq based measure of PRDM9 binding asymmetry, namely, the fraction of ATAC fragments in hotspot centers in PWD/B6 regions that are assigned to the B6 homolog, for two types of hotspots within that background, controlled by the PWD and B6 PRDM9 alleles, respectively. The y-axis is the (PRDM9-binding) heat of hotspots, calculated as the fraction of ATAC fragments in a given hotspot’s center (vs. all hotspots in the same background). (B) The x-axis shows the fraction of ATAC fragments in hotspot centers in the PWD/CAST regions that are assigned to the CAST homolog, for two types of hotspots within that background, controlled by the PWD and B6 PRDM9 alleles, respectively. The y-axis is the (PRDM9-binding) heat of hotspots, calculated as the fraction of ATAC fragments in a given hotspot’s center (vs. all hotspots in the same background).

**Supplementary Fig. 7.**
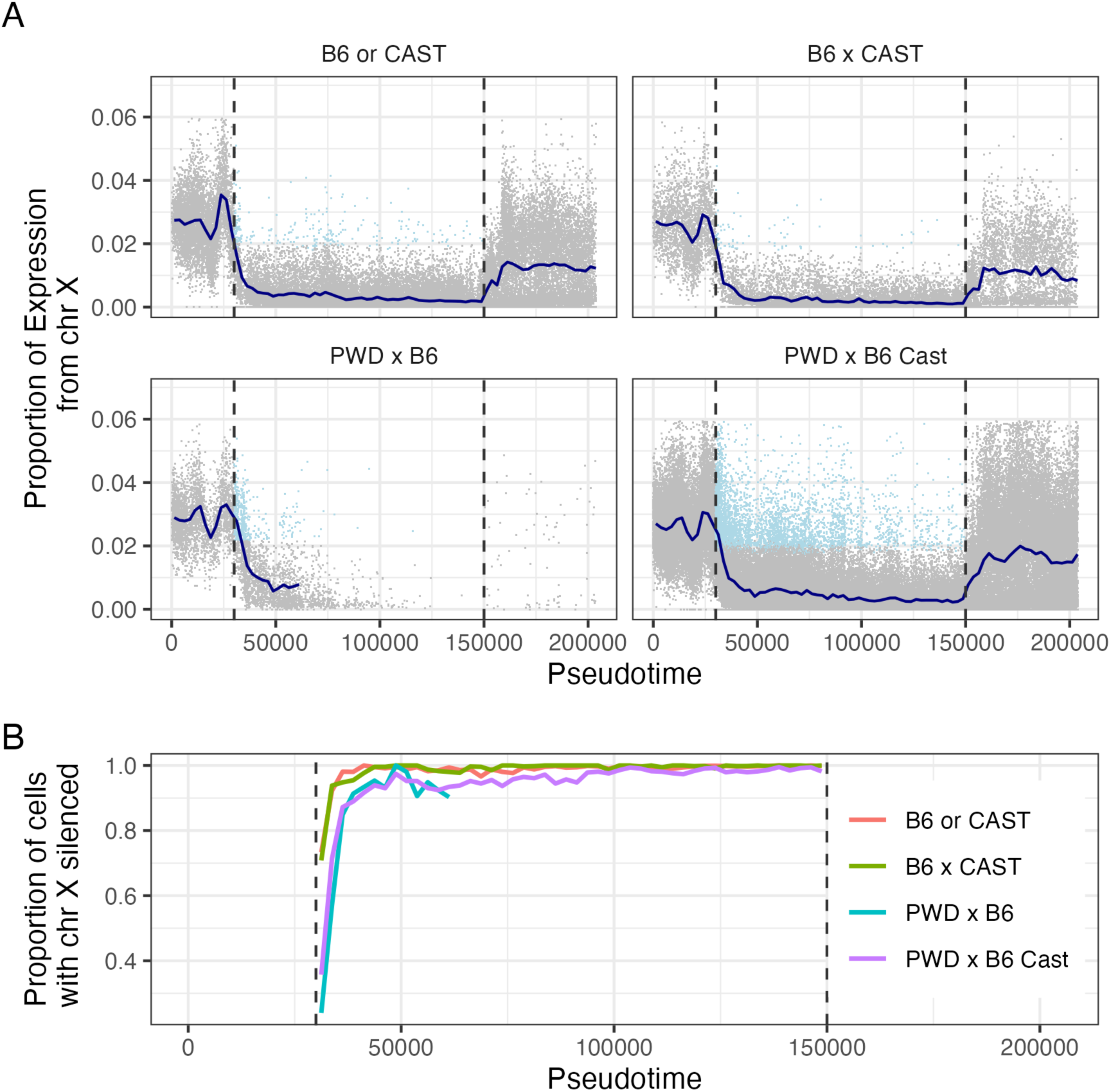
Transcriptional Silencing on the X chromosome by genetic background. (A) The proportion of gene expression reads per cell from the X chromosome. The navy line indicates the average value in a given Pseudotime bin (2500 cells). Cells with likely aberrant silencing (corresponding to >75% of pre-pachytene expression) are highlighted in blue. Vertical dashed lines indicate the Pseudotime range corresponding to the period of X silencing. (B) The proportion of cells in pachytene with X chromosome silencing (≤75% of pre-pachytene expression) through Pachytene, by genetic background.

**Supplementary Fig. 8.**
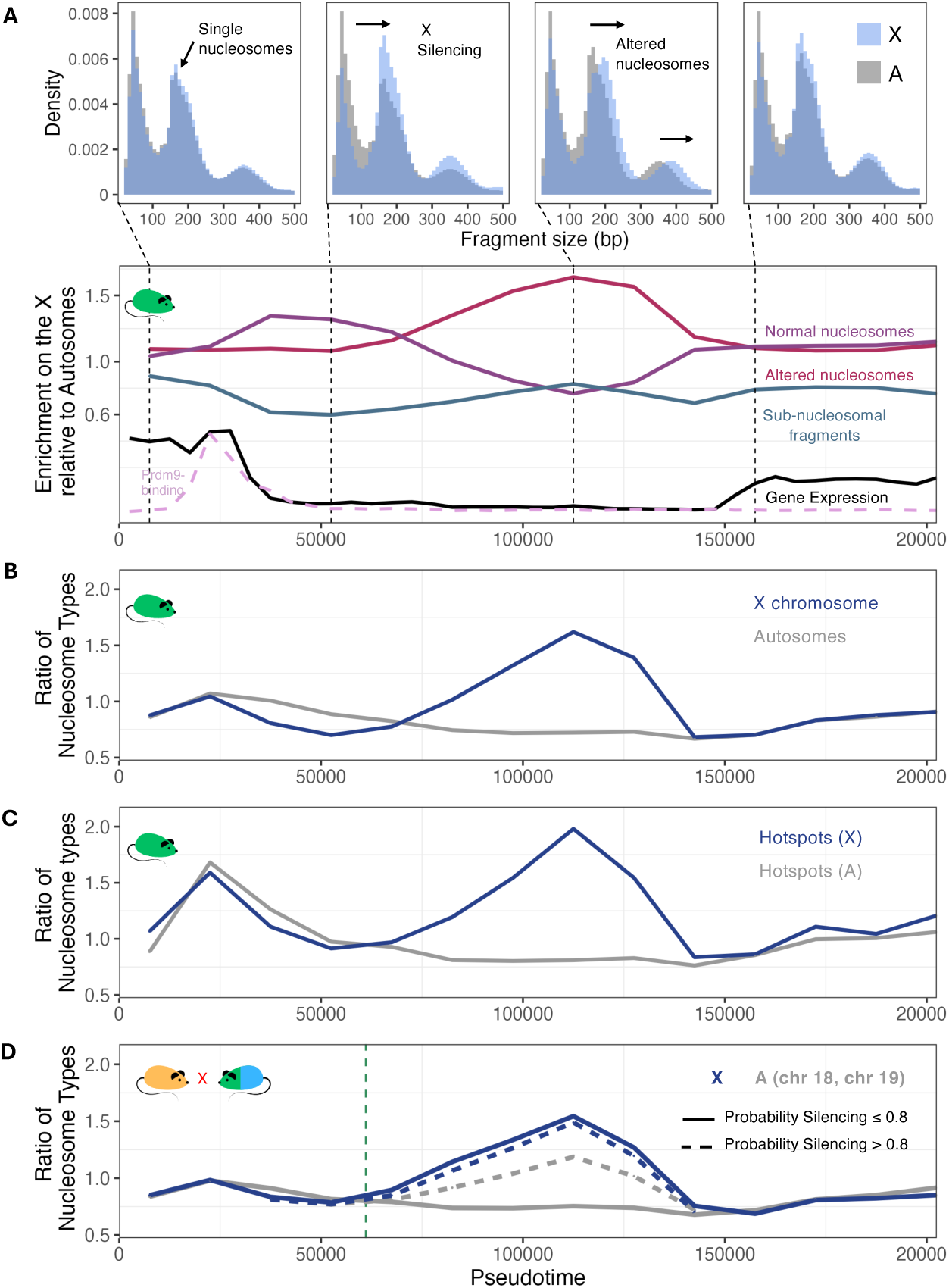
Chromatin alterations on the X chromosome and autosomes through pseudotime. (A) Comparison of ATAC fragment sizes on the X chromosome and autosomes through pseudotime in normal B6 mice. The top panel shows fragment size distributions in four pseudotime windows (pseudotime indicated by dashed lines). The bottom panel shows the ratio through pseudotime (the median value in cells in bins of 15,000 cells) of the number of ATAC fragments in a given size category on the X chromosome vs. autosomes, where size categories are: Sub-nucleosomal fragments (30-120 bp), Normal nucleosomes (150-190) bp, and Altered nucleosomes (210-300 bp) (see also Supplementary Fig. 9). The timing of transcriptional silencing on the X chromosome and PRDM9 binding at recombination hotspots in the same cells is shown for reference. The solid black line indicates (scaled by a factor of 20 to aid visualization) total gene expression on the X vs. autosomes and the dashed pink line indicates chromatin accessibility at PRDM9 motifs (scaled by a factor of 200). (B) The ratio of fragment counts corresponding to the Normal and Altered nucleosome size categories, separately on the X chromosome and Autosomes in B6 mice. Lines indicate the average in bins of 15,000 cells ordered by pseudotime. (C) The ratio of fragment counts corresponding to the Normal and Altered nucleosome size categories in PRDM9 hotspots (± 300 bp around hotspot centers) on the X chromosome and Autosomes (in B6 mice). Lines indicate the average in bins of 15,000 cells ordered by pseudotime. (D) The ratio of fragment counts corresponding to the Normal and Altered nucleosome size categories on chromosomes X, 18, and 19 in PWD x (B6 x CAST) hybrids. Lines indicate the average in bins of 15,000 cells ordered by pseudotime. Blue lines indicate the X chromosome: the solid line indicates the ratio of nucleosome types on the X for cells with weaker (or absent) silencing (≤80% inferred probability of silencing) on both chromosomes 18 *and* 19; the dashed line shows the same value for cells with strong silencing (>80% inferred probability of silencing) on chromosomes 18 *or* 19. Grey lines indicate chromosomes 18 and 19: the solid line indicates the ratio of nucleosome types on chromosome 18 or 19 in cells with weaker (or absent) silencing on both chromosomes (≤80% inferred probability of silencing for each chromosome); the dashed line indicates this value on chromosomes 18 or 19 when they are strongly silenced (>80% inferred probability of silencing). The dashed vertical line marks the end of the pseudotime interval containing 90% of all cells in sterile PWD x B6 mice (cell rank 61,090 of 203,869).

**Supplementary Fig. 9.**
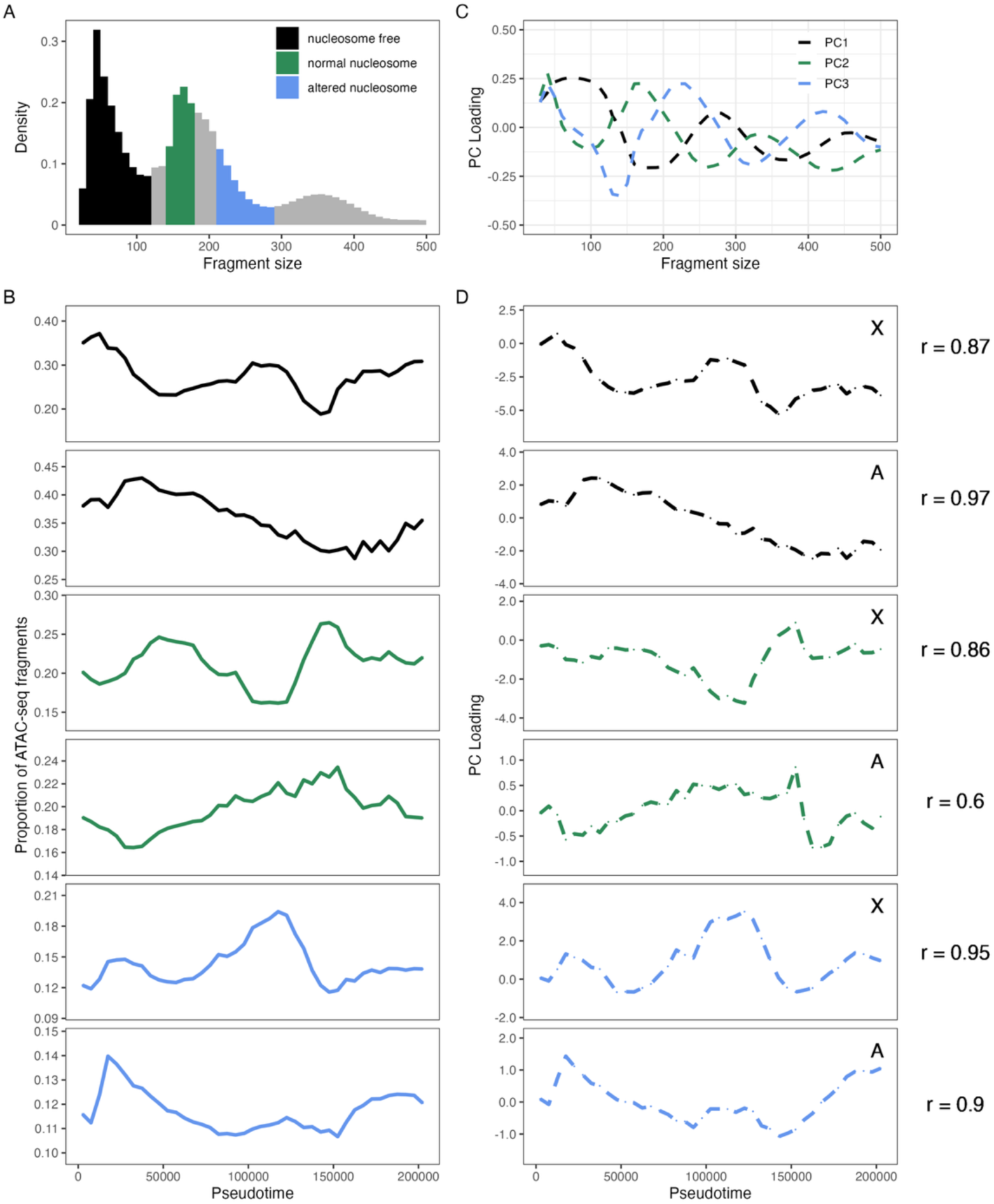
Variation in ATAC-seq fragment-size distributions on the X chromosome and autosomes, through Pseudotime. (A) The autosomal ATAC fragment size distribution, with shaded regions indicating fragment size categories used to define the nucleosome types. (B) The proportion of ATAC-seq fragments by size category on the X and Autosomes, through Pseudotime. Lines represent median proportions in Pseudotime bins of 5000 cells each. (C) Fragment size loadings for the first 3 PCs from a principal component analysis of fragment size distributions by cell and chromosome (see Methods). (D) Median loadings on the X and autosomes of the first 3 PCs (explaining 16%, 4.5%, and 4% of the variance, respectively) through Pseudotime (in bins of 5000 cells). The correlations between the median time profiles of the three PCs on the X and autosomes in panel D and the corresponding fragment size categories in panel B are indicated on the right.

**Supplementary Fig. 10.**
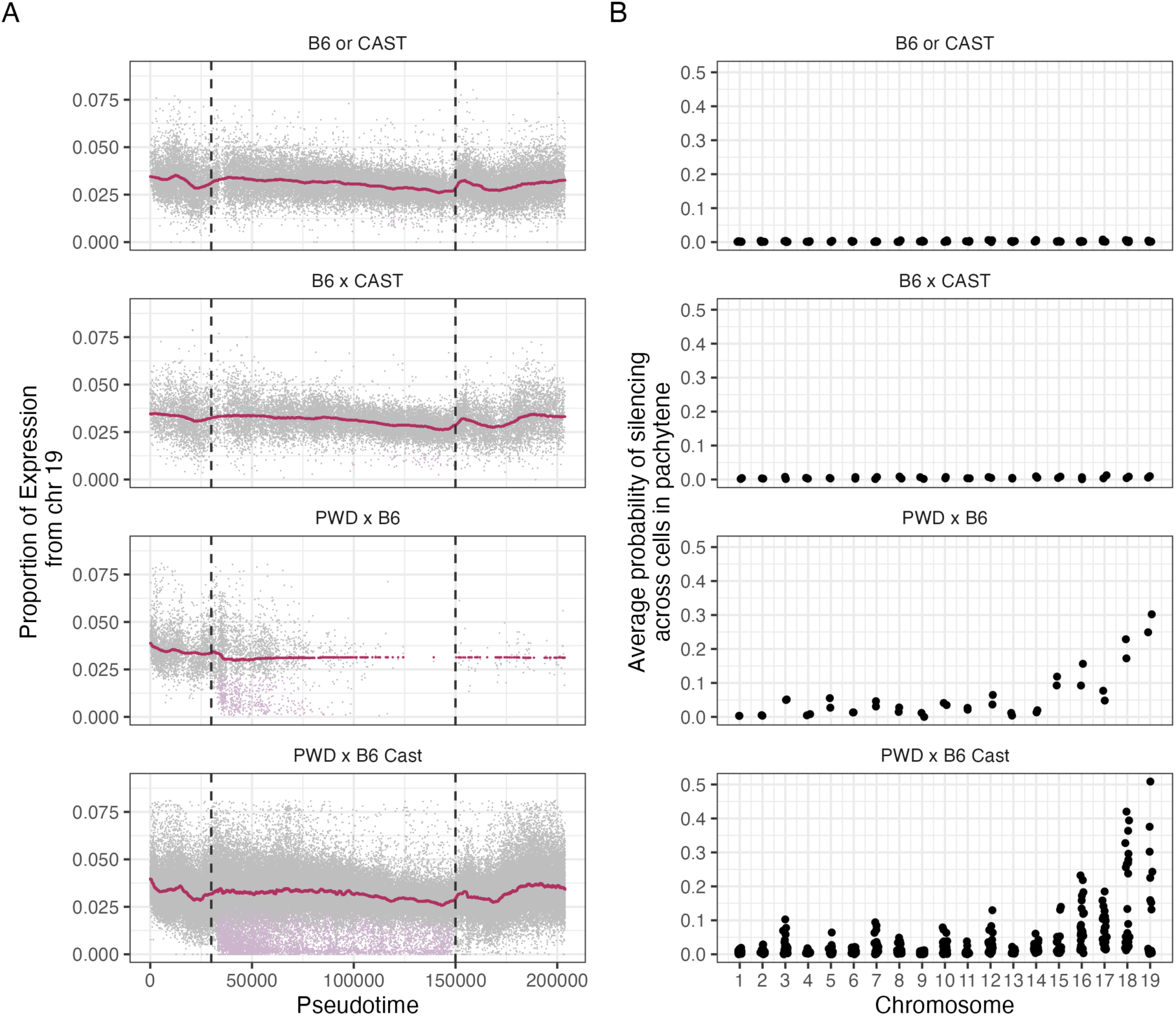
Transcriptional Silencing on Autosomes by genetic background. (A) The proportion of gene expression reads per cell from Chromosome 19. Points denote individual cells from all samples of each genotype. The maroon line indicates the median cell for a genotype in fixed Pseudotime bins (of 2500 cells). Cells with >50% probability of silencing (see Methods) on Chromosome 19 are highlighted in lilac. Vertical dashed lines indicate the Pseudotime range that corresponds to the interval of X chromosome silencing and likely corresponds to Pachytene. (B) Estimated probability of silencing (averaged over cells in Pachytene) by chromosome and genotype.

**Supplementary Fig. 11.**
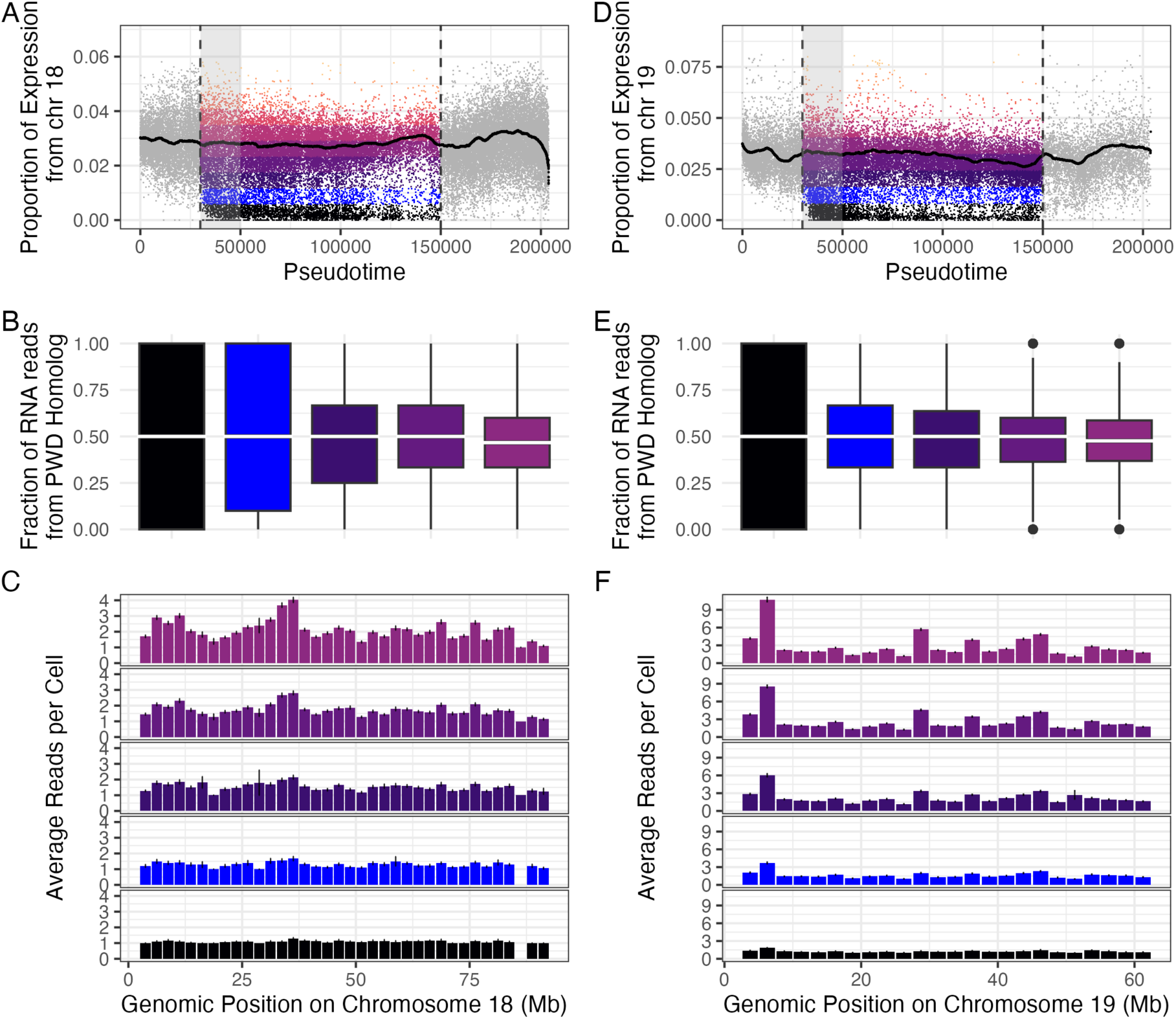
Homolog-specific and region-specific gene expression on B6/PWD chromosomes 18 and 19 in PWD x (B6 x CAST) hybrids. Colors represent bins of gene expression (see panels A and D). The grey shaded region indicates the Pseudotime interval sampled (early pachytene). (A) Proportion of gene expression from chromosome 18 in individual cells through pseudotime. Vertical lines represent the pseudotime interval likely corresponding to pachytene. (B) The proportion of RNA reads from the PWD (vs non-PWD) homologous chromosome for chromosome 18, by the level of gene expression (binned). (C) The number of RNA reads per cell in bins of 2.5 Mb along chromosome 18, averaged over cells binned by the level of gene expression. Standard errors are indicated as vertical bars. (D-F) Same as panels A-C, for chromosome 19.

**Supplementary Fig. 12.**
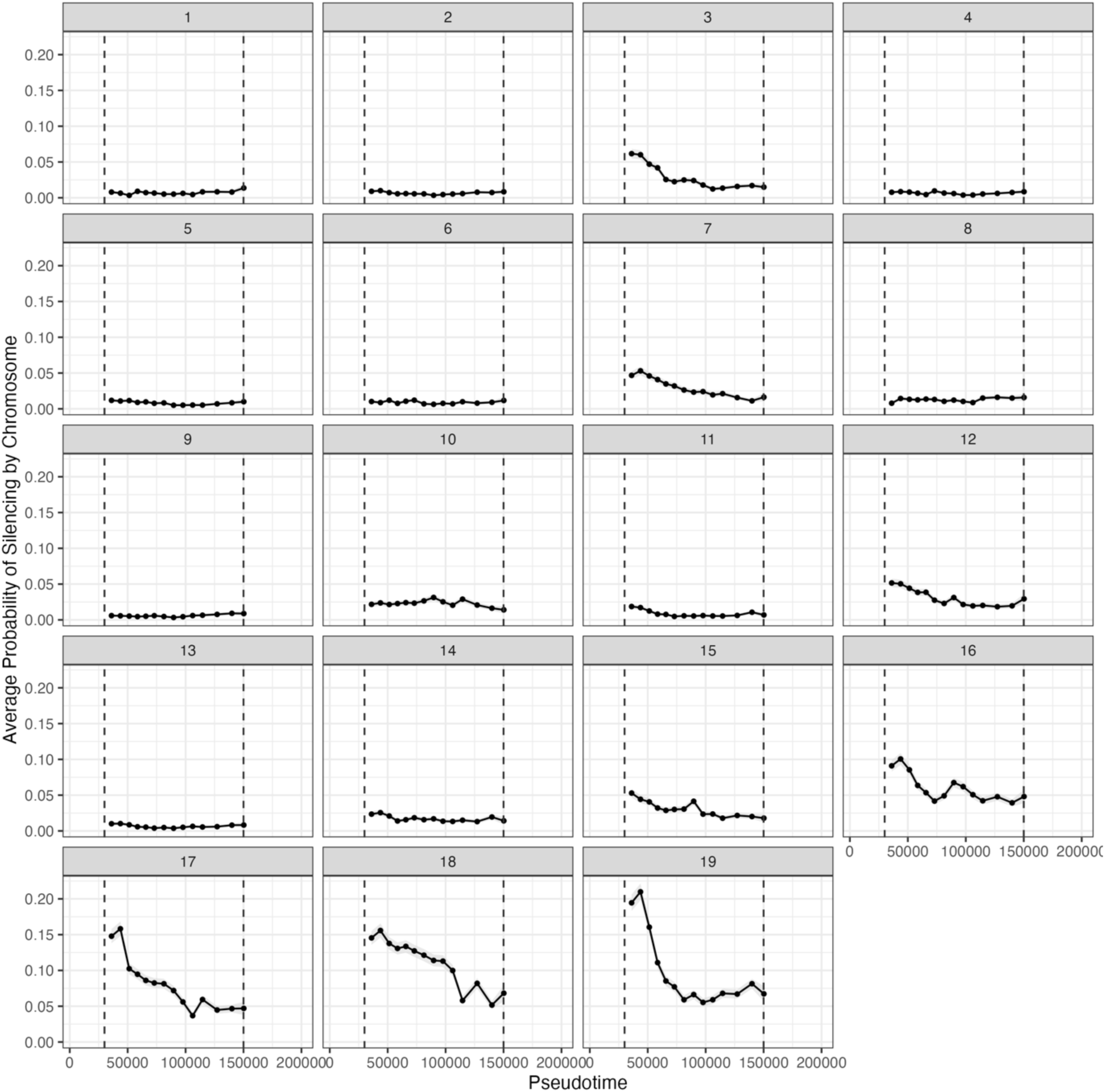
Estimated probability of silencing by chromosome across PWD x (B6 x CAST) hybrids through Pseudotime. Points show the average value in bins of 5000 cells and the 95% confidence interval is indicated by the grey ribbon.

**Supplementary Fig. 13.**
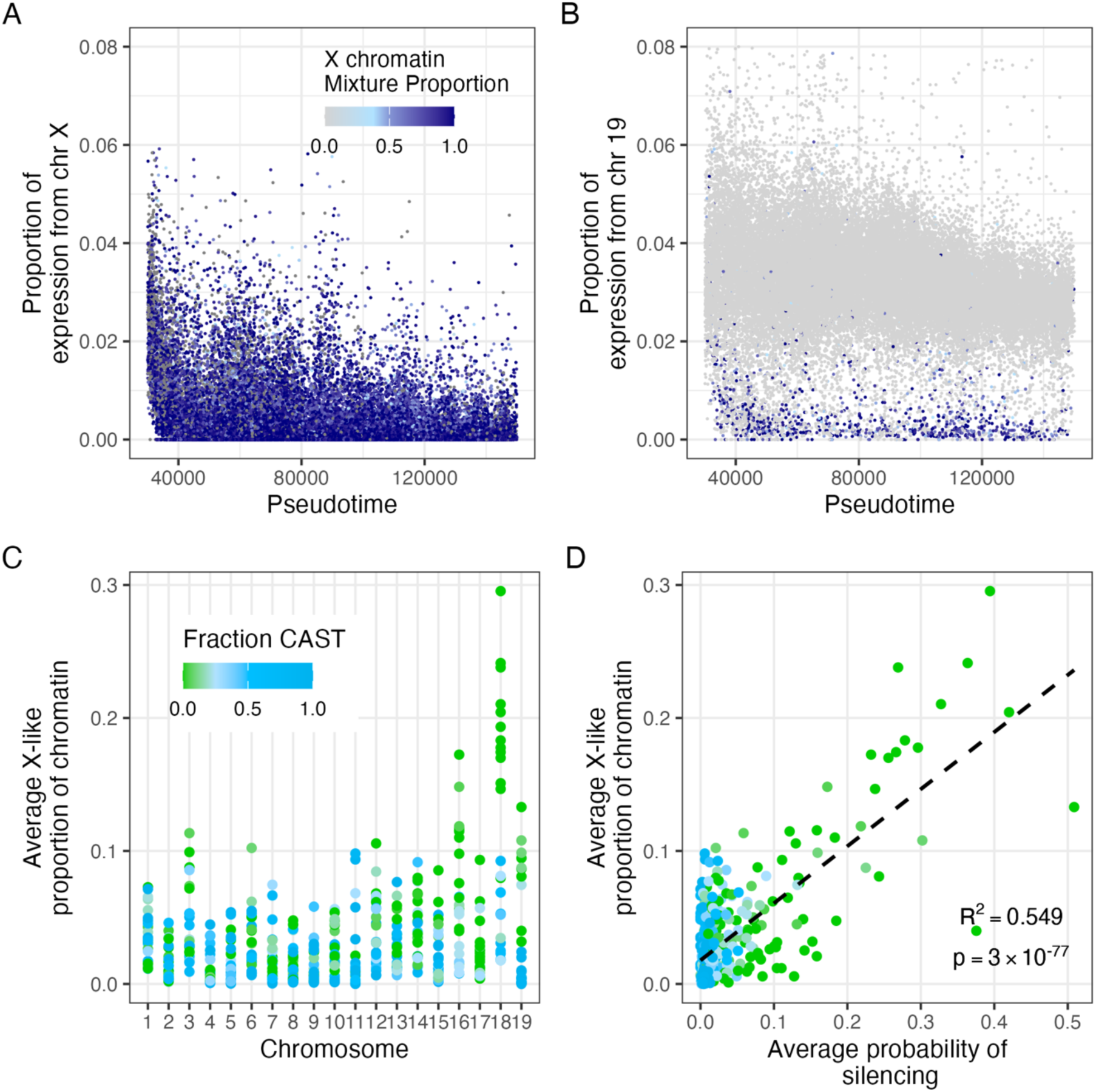
X-chromosome-like chromatin in single cells. (A) Single-cell measurements of X chromosome chromatin signatures during pachytene in PWD x (B6 x CAST) mice. Each point represents a cell, ordered by pseudotime, and colored by the mixture proportion (λ) of X-like chromatin. λ represents the cell-normalized signal (raw chrX signal minus mean autosomal signal) (see Methods). The y-axis shows the proportion of expression from the X chromosome. (B) Single-cell measurements of an X chromosome-like chromatin signature on chromosome 19 during pachytene. Each point represents a cell, ordered by pseudotime, and colored by the mixture proportion (λ) of X-like chromatin. λ is adjusted relative to the mean of the other 18 autosomes to highlight chromosome-specific enrichment of X-like chromatin (vs cell-level signal) (see Methods). The y-axis shows the proportion of total cellular expression from chr 19. (C) Average X-like chromatin proportion across autosomes (chr 1-19) in hybrid animals. Each point represents one animal-chromosome combination, colored by the fraction of CAST ancestry for that chromosome. (D) Relationship between X-like chromatin and transcriptional silencing/asynapsis across autosomes. Each point represents one animal-chromosome combination. Points are colored by the proportion of CAST ancestry. The x-axis shows the average probability of silencing (calculated from RNA-seq), and the y-axis shows the average X-like chromatin proportion (estimated from ATAC-seq fragment lengths) (see Methods).

**Supplementary Fig. 14.**
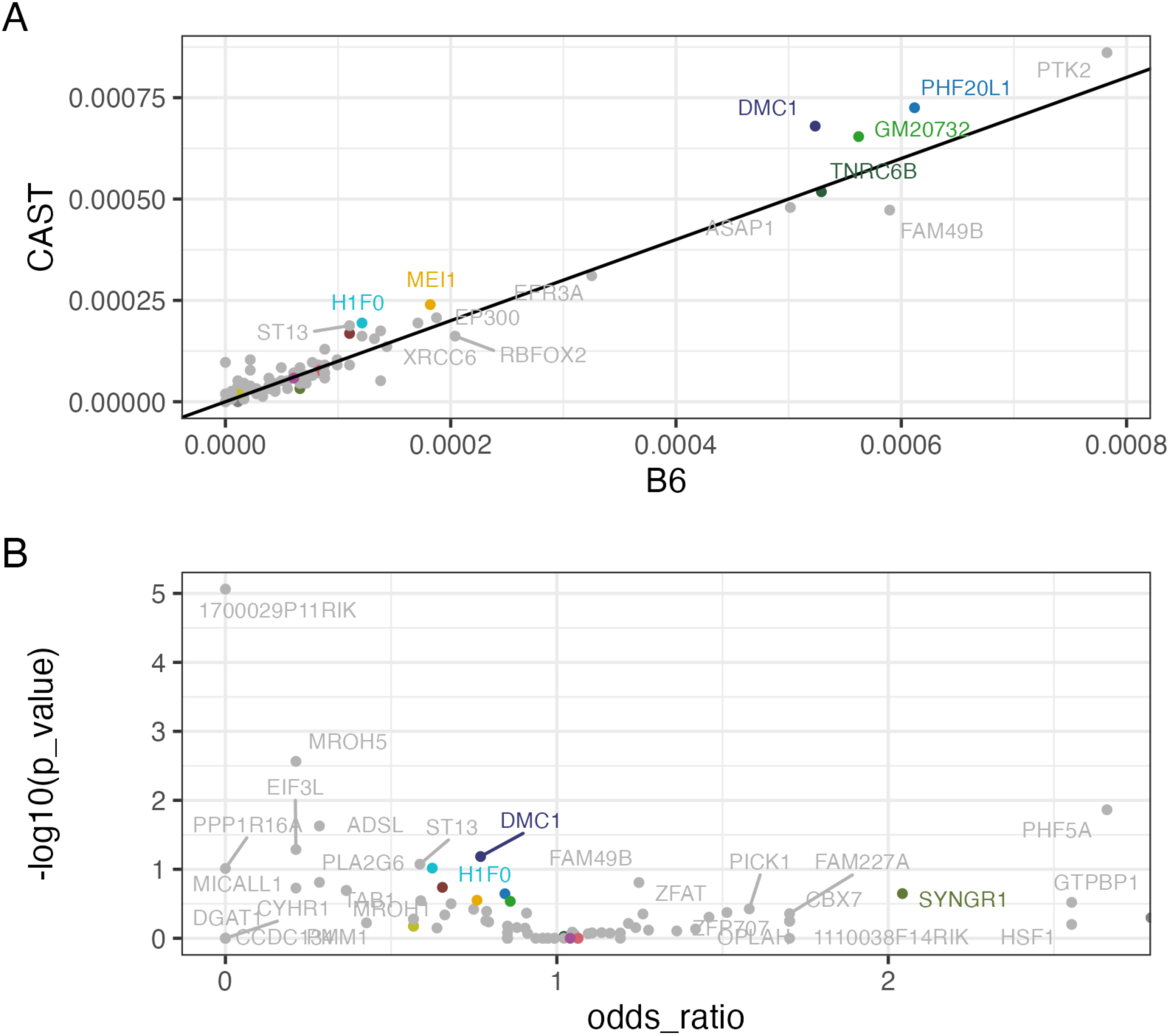
Allele specific gene expression (B6 vs. CAST, in the B6 x CAST animals) for genes in the Chromosome 15 locus. (A) The proportion of all B6 RNA reads genome-wide in a gene vs the proportion of all CAST RNA reads genome-wide in a gene, for genes in the Chromosome 15 locus with at least 10 background informative reads. Genes expressed early in meiosis and representing stronger candidates for affecting synapsis (see also Fig. 4E) are indicated with color, and others are indicated in grey. (B) The odds ratio (CAST vs B6 expression) and p-values from Fisher Exact tests for the genes shown in panel A (and colored as in panel A).

**Supplementary Fig. 15.**
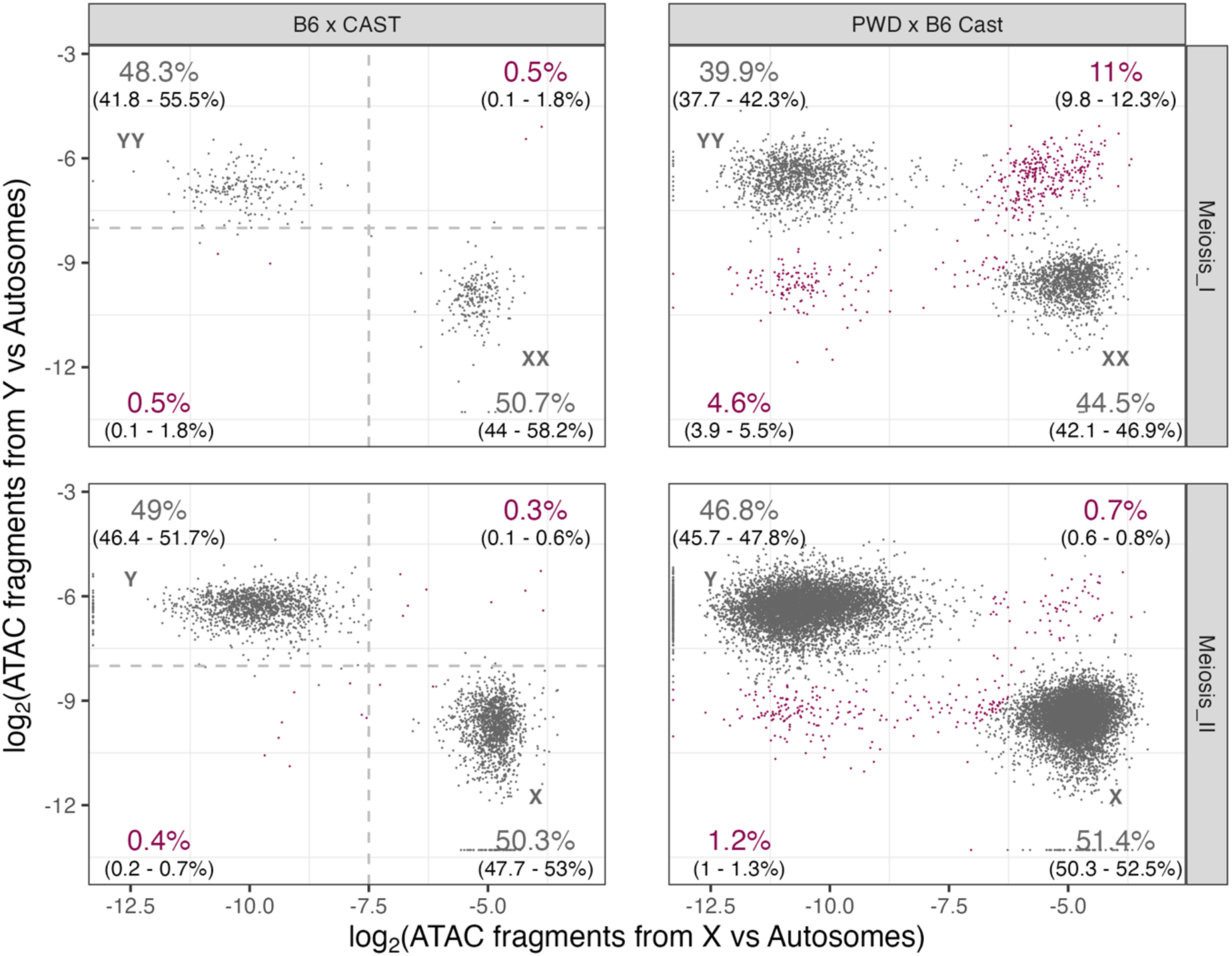
Classification of X and/or Y-bearing MI and MII cells using ATAC-seq. X and Y chromosome content of individual MI and MII cells in PWD x (B6 x CAST) and fertile B6 x CAST hybrids. The x-axis shows the log^2^-transformed ratio of ATAC-seq fragment counts from the X chromosome relative to autosomes; the y-axis shows the corresponding ratio for the Y chromosome. Normal X or Y-bearing cells are shown in grey (XX/YY in MI cells and X/Y in MII cells. Abnormal populations of cells with sex chromosome duplications or deletions are shown in maroon. Percentages indicate the estimated proportions (see Methods) of each type of cell, with 95% confidence intervals calculated assuming Poisson-distributed counts (in parentheses). Grey dashed lines indicate initial classification thresholds based on the distribution of fertile B6 x CAST cells (see Methods).

**Supplementary Fig. 16.**
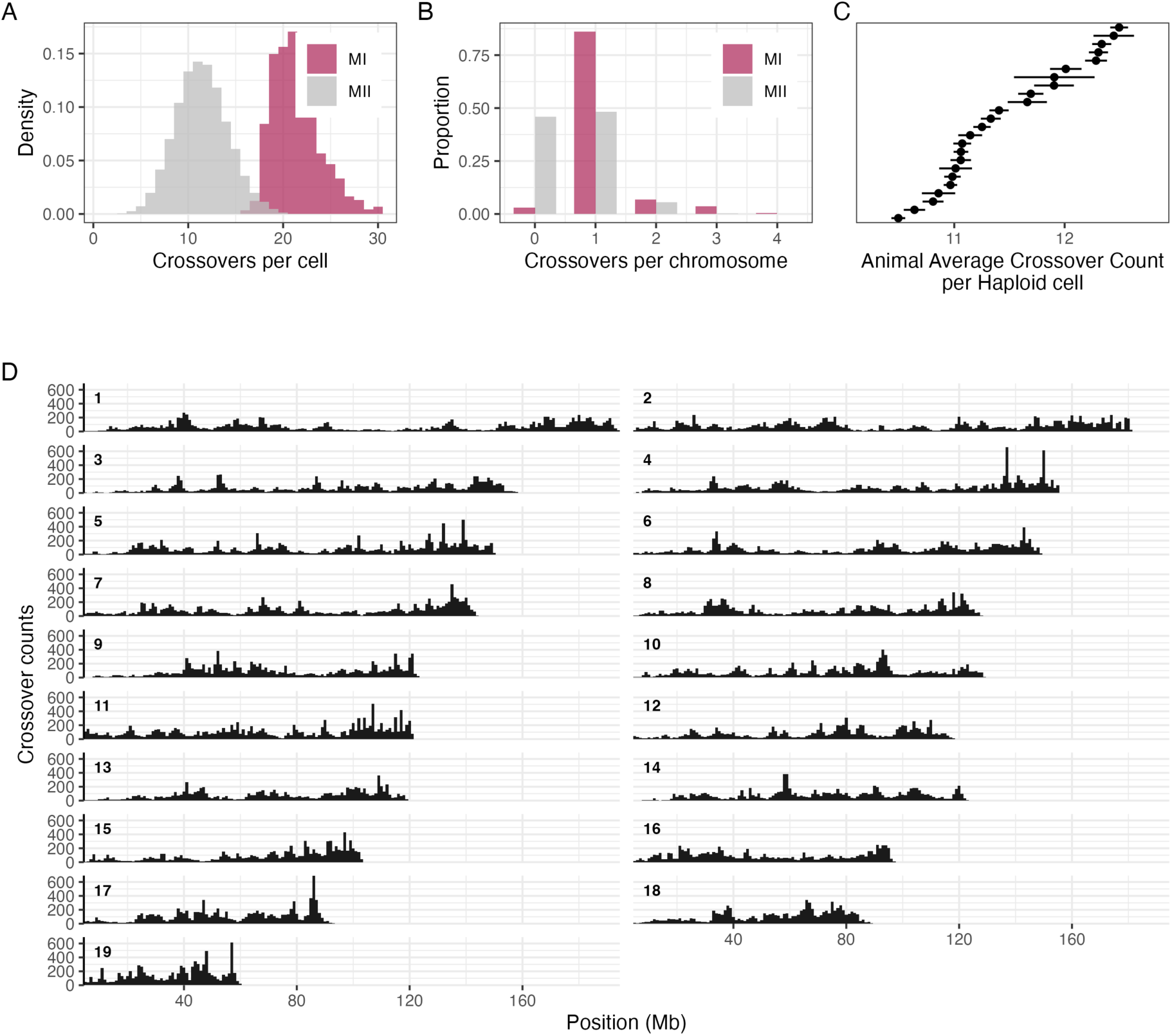
Summary of crossover calls in post-division (MI and MII) cells in PWD x (B6 x CAST) and fertile B6 x CAST mice. (A) The distribution of crossover counts per cell in MI and MII cells for all samples. (B) The distribution of crossover calls per chromosome in MI and MII cells for all samples. (C) Animal-level estimates of crossover rates (estimated from MII cells, and averaged over all cells in the animal). Error bars denote the standard error. (D) Crossover density across the genome in 1 Mb bins, with each chromosome normalized separately.

**Supplementary Fig. 17.**
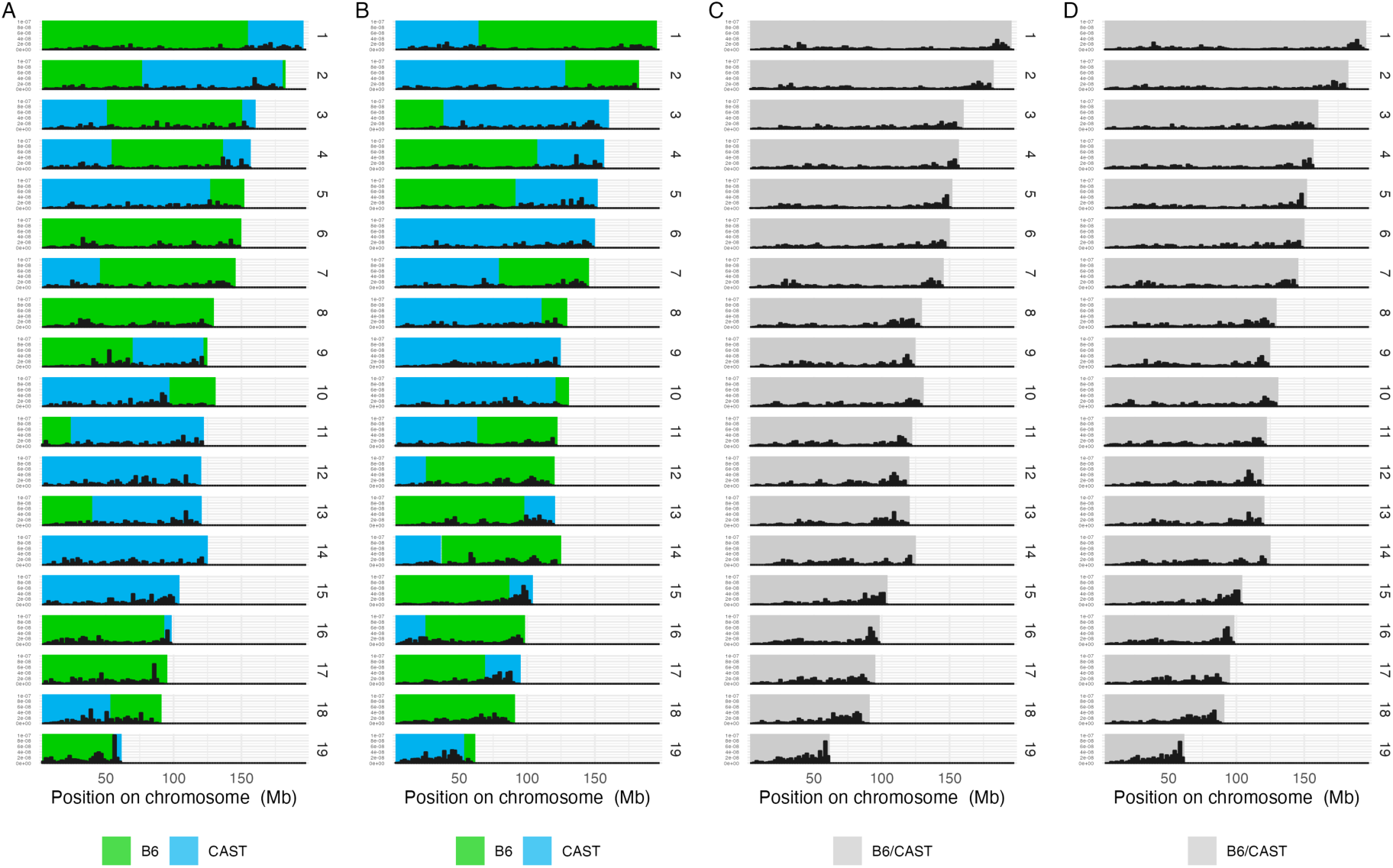
Observed crossover distributions on each chromosome for two PWD x (B6 x CAST F1/F2) and two B6 x CAST animals. Crossover counts are aggregated per Mb, normalized per chromosome and divided by the number of cells in each animal. (A-B) Crossover densities genome-wide for two PWD x (B6 x CAST) animals, with the PWD/B6 and the PWD/CAST backgrounds shown in green and blue, respectively. (C-D) Crossover densities genome-wide in two B6 x CAST animals with humanized *Prdm9* alleles.

**Supplementary Table 1:**
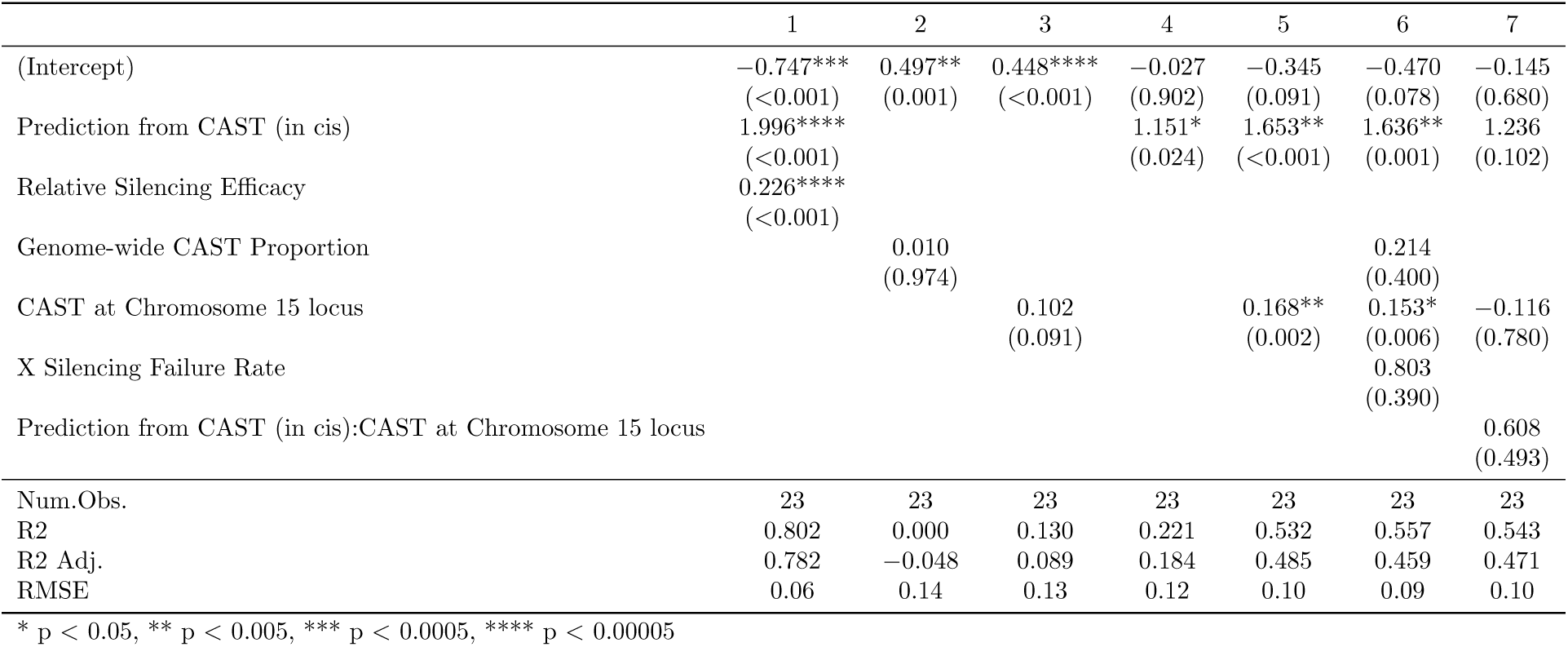
Predicting Overall Autosomal Asynapsis Rates.

**Supplementary Table 2:** Predicting Failure of X chromosome silencing.

|  | 1 | 2 | 3 | 4 | 5 | 6 |
| --- | --- | --- | --- | --- | --- | --- |
| (Intercept) | 0.007<br>(0.884) | −0.015<br>(0.785) | 0.111***<br>($<0.001$) | 0.065<br>(0.301) | 0.103**<br>(0.001) | 0.082*<br>(0.013) |
| Prediction from CAST (in cis) | 0.105<br>(0.330) | 0.141<br>(0.228) |  | 0.084<br>(0.436) |  |  |
| CAST at Chromosome 15 locus |  | 0.012<br>(0.376) |  | 0.015<br>(0.220) |  |  |
| Genome-wide CAST Proportion |  |  | −0.123*<br>(0.028) | −0.125*<br>(0.036) | −0.130*<br>(0.026) | −0.124*<br>(0.024) |
| Relative Silencing Efficacy |  |  |  |  | 0.007<br>(0.517) |  |
| Observed Asynapsis Rate |  |  |  |  |  | 0.059<br>(0.134) |
| Num.Obs. | 23 | 23 | 23 | 23 | 23 | 23 |
| R2 | 0.045 | 0.083 | 0.209 | 0.277 | 0.225 | 0.295 |
| R2 Adj. | 0.000 | −0.009 | 0.171 | 0.163 | 0.148 | 0.224 |
| RMSE | 0.03 | 0.03 | 0.03 | 0.02 | 0.03 | 0.02 |

\* $p < 0.05$ , \*\* $p < 0.005$ , \*\*\* $p < 0.0005$ , \*\*\*\* $p < 0.00005$

**Supplementary Table 3:** Predicting Cell Survival in Pachytene.

|  | 1 | 2 | 3 | 4 | 5 | 6 | 7 |
| --- | --- | --- | --- | --- | --- | --- | --- |
| (Intercept) | 0.429****<br>($<0.001$) | 0.683****<br>($<0.001$) | 0.736****<br>($<0.001$) | 0.494*<br>(0.017) | 0.176<br>(0.225) | 0.839**<br>(0.003) | 0.877****<br>($<0.001$) |
| X Silencing Failure Rate | -3.067**<br>(0.003) |  | -2.026**<br>(0.002) | -2.112**<br>(0.004) |  |  | -4.568*<br>(0.032) |
| Observed Asynapsis Rate | | -0.846****<br>($<0.001$) | -0.727****<br>($<0.001$) | -0.878****<br>($<0.001$) | | | -1.000***<br>($<0.001$) |
| Prediction from CAST (in cis) |  |  |  | 0.661<br>(0.111) |  | -1.094*<br>(0.047) |  |
| CAST at Chromosome 15 locus |  |  |  | 0.023<br>(0.593) |  | -0.150*<br>(0.021) |  |
| Genome-wide CAST Proportion |  |  |  | 0.014<br>(0.935) | 0.185<br>(0.548) |  |  |
| Observed Asynapsis Rate:X Silencing Failure Rate |  |  |  |  |  |  | 4.720<br>(0.197) |
| Num.Obs. | 23 | 23 | 23 | 23 | 23 | 23 | 23 |
| R2 | 0.349 | 0.648 | 0.788 | 0.823 | 0.017 | 0.286 | 0.806 |
| R2 Adj. | 0.318 | 0.632 | 0.767 | 0.771 | -0.029 | 0.214 | 0.776 |
| RMSE | 0.12 | 0.09 | 0.07 | 0.06 | 0.15 | 0.13 | 0.07 |

\* $p < 0.05$ , \*\* $p < 0.005$ , \*\*\* $p < 0.0005$ , \*\*\*\* $p < 0.00005$

**Supplementary Table 4:** Predicting Testes Weight.

|  | 1 | 2 | 3 | 4 | 5 | 6 | 7 |
| --- | --- | --- | --- | --- | --- | --- | --- |
| (Intercept) | 66.135****<br>($<0.001$) | 175.094****<br>($<0.001$) | 113.216*<br>(0.047) | 63.382<br>(0.274) | 72.148****<br>($<0.001$) | 148.831*<br>(0.025) | 71.444<br>(0.134) |
| Cell Survival in Pachytene | 141.727***<br>($<0.001$) | | 84.111<br>(0.233) | 118.122**<br>(0.004) | 65.440<br>(0.111) | | 97.998**<br>(0.005) |
| Observed Asynapsis Rate |  | -93.117*<br>(0.015) | -31.976<br>(0.603) |  |  |  |  |
| X Silencing Failure Rate |  | -460.960*<br>(0.015) | -290.569<br>(0.202) |  |  |  |  |
| Prediction from CAST (in cis) |  |  |  | 19.433<br>(0.837) |  | -100.235<br>(0.352) | 32.047<br>(0.676) |
| CAST at Chromosome 15 locus |  |  |  | -19.590<br>(0.098) |  | -37.827**<br>(0.005) | 7.478<br>(0.551) |
| Genome-wide CAST Proportion |  |  |  | 22.563<br>(0.625) |  | 43.730<br>(0.436) | 29.800<br>(0.427) |
| Sperm Count |  |  |  |  | 0.937*<br>(0.013) |  |  |
| CAST at Chromosome 11 locus |  |  |  |  |  |  | -39.901**<br>(0.005) |
| Num.Obs. | 23 | 23 | 23 | 23 | 23 | 23 | 23 |
| R2 | 0.494 | 0.500 | 0.537 | 0.591 | 0.630 | 0.350 | 0.748 |
| R2 Adj. | 0.470 | 0.450 | 0.464 | 0.500 | 0.593 | 0.248 | 0.674 |
| RMSE | 21.32 | 21.20 | 20.40 | 19.18 | 18.23 | 24.17 | 15.04 |

\* $p < 0.05$ , \*\* $p < 0.005$ , \*\*\* $p < 0.0005$ , \*\*\*\* $p < 0.00005$

**Supplementary Table 5:** Predicting Sperm Count (in Millions)

|  | 1 | 2 | 3 | 4 | 5 | 6 | 7 |
| --- | --- | --- | --- | --- | --- | --- | --- |
| (Intercept) | −6.418<br>(0.228) | 56.981****<br>($<0.001$) | 26.747<br>(0.366) | 13.571<br>(0.683) | −25.375*<br>(0.006) | 71.172<br>(0.074) | 15.660<br>(0.634) |
| Cell Survival in Pachytene | 81.417***<br>($<0.001$) | | 41.098<br>(0.281) | 79.625**<br>(0.001) | 40.790<br>(0.070) | | 74.411**<br>(0.003) |
| X Silencing Failure Rate |  | −275.104*<br>(0.008) | −191.849<br>(0.124) |  |  |  |  |
| Observed Asynapsis Rate |  | −53.958*<br>(0.010) | −24.084<br>(0.472) |  |  |  |  |
| Prediction from CAST (in cis) |  |  |  | −50.233<br>(0.366) |  | −130.901<br>(0.058) | −46.965<br>(0.394) |
| CAST at Chromosome 15 locus |  |  |  | 2.969<br>(0.654) |  | −9.324<br>(0.220) | 9.982<br>(0.269) |
| Genome-wide CAST Proportion |  |  |  | 4.364<br>(0.870) |  | 18.633<br>(0.589) | 6.239<br>(0.814) |
| Testes weight |  |  |  |  | 0.287*<br>(0.013) |  |  |
| CAST at Chromosome 11 locus |  |  |  |  |  |  | −10.338<br>(0.251) |
| Num.Obs. | 23 | 23 | 23 | 23 | 23 | 23 | 23 |
| R2 | 0.513 | 0.544 | 0.572 | 0.565 | 0.644 | 0.222 | 0.598 |
| R2 Adj. | 0.490 | 0.499 | 0.504 | 0.469 | 0.608 | 0.099 | 0.480 |
| RMSE | 11.79 | 11.41 | 11.06 | 11.15 | 10.09 | 14.91 | 10.71 |

\* $p < 0.05$ , \*\* $p < 0.005$ , \*\*\* $p < 0.0005$ , \*\*\*\* $p < 0.00005$

**Supplementary Table 6:** Abnormalities in Late vs Early Meiosis-I cells in PWD x (B6 x CAST) hybrids Cell-level abnormalities Chromosome-type abnormalities.

Cell-level abnormalities
|  | Cell Count (early) | Cell count (late) | Fraction (early) | Fraction (late) | Fold | OR | P |
| --- | --- | --- | --- | --- | --- | --- | --- |
| Any Deletion (sex chromosomes or autosomes) | 1077 | 1867 | 0.0223 | 0.1061 | 4.7591 | 5.2029 | 3.50e-19*** |
| Any Autosomal Homozygous Deletion | 1080 | 1871 | 0.0074 | 0.0428 | 5.7723 | 5.9821 | 3.39e-09*** |
| Any low copy autosome | 1080 | 1871 | 0.1676 | 0.2774 | 1.6552 | 1.9062 | 7.28e-12*** |
| Any high copy autosome | 1080 | 1871 | 0.0194 | 0.0470 | 2.4189 | 2.4882 | 9.93e-05*** |
| Any non-MI-recombinant | 1080 | 1871 | 0.7417 | 0.7980 | 1.0759 | 1.3756 | 4.76e-04*** |
| X and Y deletion | 1077 | 1867 | 0.0149 | 0.0643 | 4.3265 | 4.5532 | 5.64e-11*** |
| Both X and Y present | 1077 | 1867 | 0.0306 | 0.1559 | 5.0869 | 5.8388 | 8.19e-30*** |
| Any X/Y abnormality | 1077 | 1867 | 0.0455 | 0.2201 | 4.8386 | 5.9191 | 7.85e-42*** |

Chromosome-type abnormalities
|  | Mean.Count.per.early.cell | Mean.Count.per.late.cell | Frac.early | Frac.late | Fold | OR | P |
| --- | --- | --- | --- | --- | --- | --- | --- |
| Aberrant/High copy number recombinants | 0.56 | 0.60 | 0.0294 | 0.0318 | 1.0808 | 1.0834 | 0.1222 |
| Heterozygous (Potential Duplication) | 0.33 | 0.40 | 0.0174 | 0.0210 | 1.2046 | 1.2089 | 0.0033** |
| Homozygous (allele 1 or 2) | 0.68 | 0.65 | 0.0356 | 0.0342 | 0.9615 | 0.9602 | 0.4019 |
| Homozygous Deletion | 0.01 | 0.05 | 4.39e-04 | 0.0025 | 5.7723 | 5.7851 | 7.04e-10*** |
| MI Recombinant | 17.43 | 17.30 | 0.9172 | 0.9105 | 0.9927 | 0.9189 | 0.0071** |

\* P < 0.05; \*\* P < 0.01; \*\*\* P < 0.001

**Supplementary Table 7:**
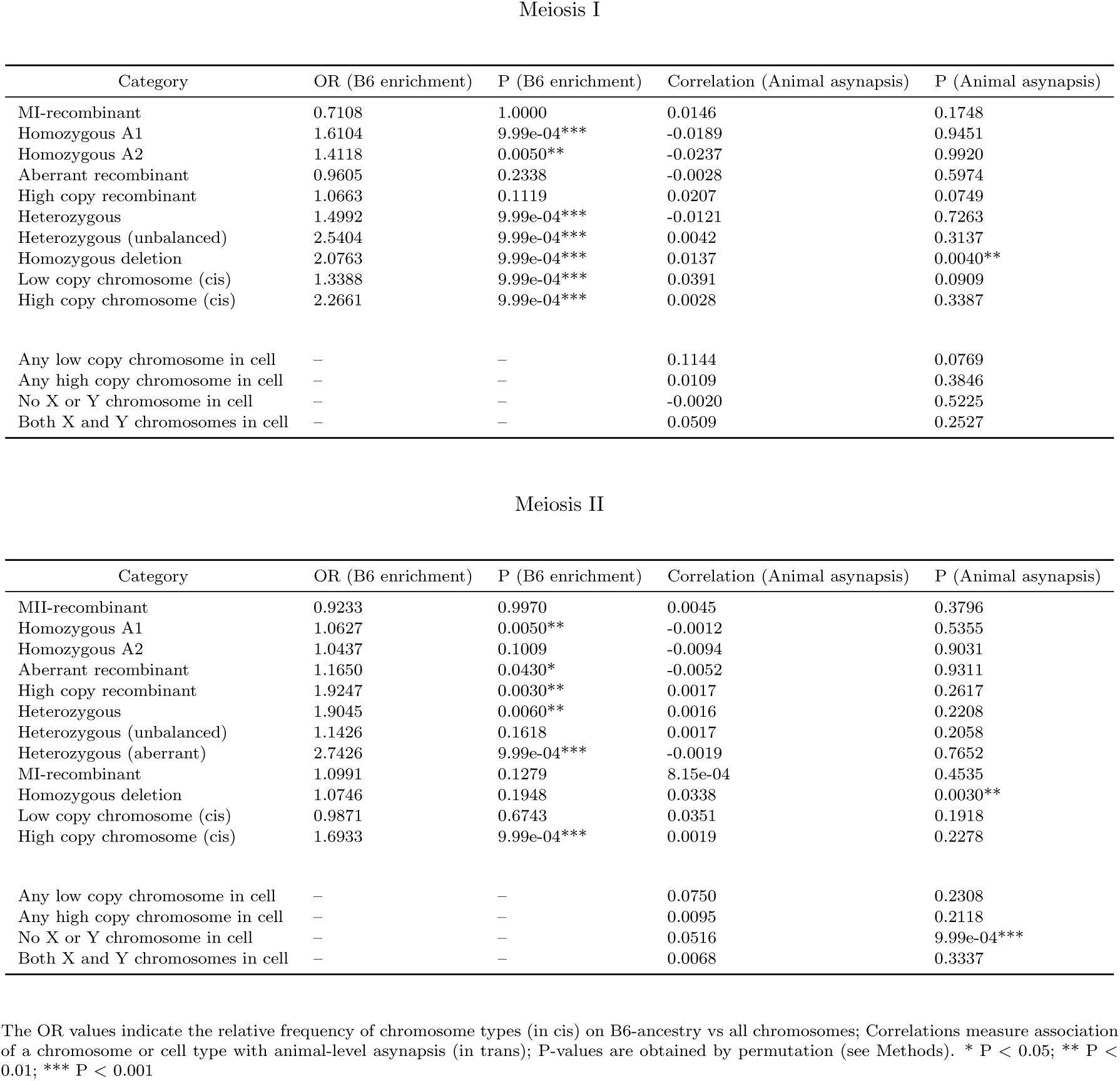
Aneuploidy Enrichment with Asynapsis in PWD x (B6 x CAST) hybrids Meiosis I Meiosis II.

**Supplementary Table 8:** Weighted linear regression: Asynapsis predictors of MII deletion rates All models use weighted linear regression (weights = cell counts per animal). Models MI–MIII: response is mean autosomal deletion rate across all PWD x (B6 x CAST) animals with *>*500 cells. Models MIV–MVI: response is chromosome X deletion rate. Predictor ‘Predicted’ is the fitted value from Model MII.

|  | MI | MII | MIII | MIV | MV | MVI |
| --- | --- | --- | --- | --- | --- | --- |
|  | Autosomal deletions |  |  | Chr X deletions |  |  |
| (Intercept) | -0.004* | 0.000 | 0.000 | -0.010 | 0.006*** | 0.004 |
|  | (0.013) | (0.949) | (0.744) | (0.109) | (<0.001) | (0.430) |
| Animal-wide Asynapsis/Silencing | 0.010** |  | 0.001 | 0.043** |  | 0.005 |
|  | (0.001) |  | (0.716) | (0.002) |  | (0.690) |
| Chromosome 19 Asynapsis/Silencing |  | 0.007*** | 0.007** |  |  |  |
|  |  | (<0.001) | (<0.001) |  |  |  |
| Chromosome 16 Asynapsis/Silencing |  | 0.009* | 0.008* |  |  |  |
|  |  | (0.005) | (0.022) |  |  |  |
| Predicted |  |  |  |  | 4.218**** | 3.916** |
|  |  |  |  |  | (<0.001) | (0.002) |
| Num.Obs. | 13 | 13 | 13 | 13 | 13 | 13 |
| R2 | 0.620 | 0.921 | 0.922 | 0.588 | 0.850 | 0.853 |
| R2 Adj. | 0.585 | 0.905 | 0.896 | 0.550 | 0.837 | 0.823 |
| RMSE | 0.00 | 0.00 | 0.00 | 0.00 | 0.00 | 0.00 |

\* p < 0.05, \*\* p < 0.005, \*\*\* p < 0.0005, \*\*\*\* p < 0.00005

## Materials and Methods

### 1 Mouse samples, tissue collection, and fertility measurements

#### 1.1 Mouse samples

C57BL/6J (B6) and PWD/PhJ (PWD) mice were purchased from Charles River Laboratories. CAST/EiJ (CAST) mice were a gift from Jonathan Godwin at the Sir William Dunn School of Pathology, University of Oxford. The Humanized *Prdm9* mouse was generated in house as described in a previous study [1]. All animal experiments received local ethical review approval from the University of Oxford Animal Welfare and Ethical Review Body (Clinical Medicine board) and were carried out in accordance with the UK Home Office Animals (Scientific Procedures) Act 1986.

PWD *×* B6 mice were generated by crossing female PWD with male B6 mice. B6 *×* CAST F1 mice were generated by crossing female B6 with male CAST mice. B6 *×* CAST F2 mice were generated by first crossing female B6 with male CAST mice and then crossing littermates to generate B6 *×* CAST F2 mice. Mice were genotyped at the *Prdm9* locus using the following primers, and standard cycling conditions: Forward 5’-CAATGTGGGCAATATTTCAG -3’; Reverse 5’-GCAGAACAGATGTTTAGTGA -3’ with an amplicon size of 1.05kb. B6 *×* CAST F1 or F2 males with B6/B6 *Prdm9* alleles were crossed with PWD females to obtain PWD *×* (B6 *×* CAST F1 or F2) mice with B6/PWD *Prdm9* alleles.

All 10x Chromium Next GEM Single Cell Multiome ATAC + Gene Expression experiments were carried out on male mice aged 8 to 24 weeks.

#### 1.2 Tissue collection and fertility measurements

Testes and cauda epididymides were isolated from 40 male mice, including 4 pilot samples that were excluded from our analyses and used only to refine the experimental protocol to target rare cell populations (see below). The cauda epididymis was dissected and cut into small pieces in PBS and incubated allowing the release of spermatozoa into the medium for counting under a microscope.

Fertility was assessed in 36 male mice (four wild-type B6, 17 PWD *×* (B6 *×* CAST F2), 8 PWD *×* (B6 *×* CAST F1), two PWD *×* B6, two B6 *×* CAST F1 mice with humanized *Prdm9*, and one CAST mouse; **Supplementary Data 1**), ranging 8 to 24 weeks of age by measuring testes weight (of the testes pair) and where possible counting sperm (per paired epididymides) in males.

### 2 Single cell experimental protocol

#### 2.1 Isolation of cell nuclei

Nuclei were isolated from mouse testes using the 10x Nuclei Isolation from Complex Tissues for Single Cell Multiome ATAC + Gene Expression Sequencing protocol 1.1 (CG000375 Rev C), with modifications to the lysis time and buffer concentration. To optimize the protocol for nuclei obtained from mouse testes cells, and ensure that the majority of the cell membranes were lysed without excessive damage to the nuclear membrane, we measured the degree of cell lysis under varying concentrations of the NP40 lysis buffer and incubation times. The optimal lysis time was determined to be when the sample reached *>* 95% dead cells while still maintaining nuclei with an intact membrane and well resolved edges. The optimal buffer concentration was found to be 0.5x: with the standard 1x NP40 lysis buffer concentration, significant blebbing was observed in the nuclei under 60x magnification. Fresh mouse testes were detunicated, weighed, and immediately immersed in 300 µL of 0.05x NP40 lysis buffer.

#### 2.2 Flow Cytometry

To assess cell-type proportions, eliminate debris and obtain single nuclei, the samples were stained with 7-aminoactinomycin D (7-AAD), a viability dye commonly used in flow cytometry to distinguish live from dead cells. Nuclei were filtered using a 40µm filter to eliminate aggregates and collected into a tube coated with 1% bovine serum albumin (BSA) and 1U/µl RNAse inhibitor in a final volume of 5ml. Nuclei were separated based on size and granularity, followed by the selection of 7-AAD-positive nuclei into tubes pre-coated with BSA and RNAse inhibitor. FACS sorting was performed using either BD FACS Aria Fusion or BD FACS Aria III using a 100 µm nozzle.

To ascertain that we capture all meiotic stages, we conducted preliminary experiments to examine the content of flow sorted populations. Nuclei were stained with (7-AAD) and flow-sorted. The observed nuclei populations, forming distinct column-like patterns, were individually gated and isolated and staged by a cytogeneticist, in particular to identify populations of prophase I nuclei. Chromosome spreads were prepared using the surface spreading technique and immunostained as previously described [1]. The following primary antibodies were used: mouse anti-SYCP3 (Santa Cruz Biotechnology, sc-74569) and rabbit anti-HORMAD2 (Santa Cruz Biotechnology, sc-82192). Alexa Fluor 594- or 488-conjugated secondary antibodies targeting rabbit or mouse IgG, respectively, were obtained from ThermoFisher Scientific. Meiotic staging was performed as previously described [2].

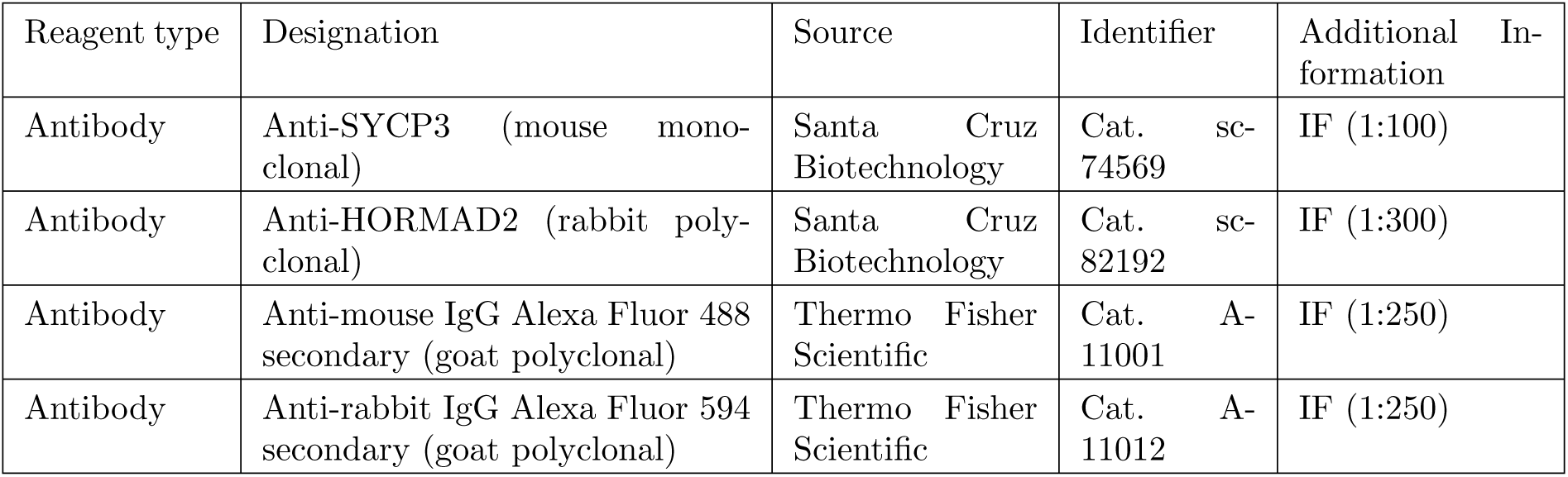

| Reagent type | Designation | Source | Identifier | Additional Information |
| --- | --- | --- | --- | --- |
| Antibody | Anti-SYCP3 (mouse monoclonal) | Santa Cruz Biotechnology | Cat. sc-74569 | IF (1:100) |
| Antibody | Anti-HORMAD2 (rabbit polyclonal) | Santa Cruz Biotechnology | Cat. sc-82192 | IF (1:300) |
| Antibody | Anti-mouse IgG Alexa Fluor 488 secondary (goat polyclonal) | Thermo Fisher Scientific | Cat. A-11001 | IF (1:250) |
| Antibody | Anti-rabbit IgG Alexa Fluor 594 secondary (goat polyclonal) | Thermo Fisher Scientific | Cat. A-11012 | IF (1:250) |

Following the successful delineation of nuclear stages (4x, 2x, and 1x nuclei) and identification of prophase-I nuclei in preliminary analyses, flow sorted nuclei in subsequent experiments were selected and gated using the same criteria to reduce debris and enrich for prophase-I nuclei. Subsequent experiments, including sequencing, were performed on these flow-sorted and gated populations. After sorting using the flow cytometer, a subset of nuclei were sampled and inspected under the microscope for damage to the nuclear membrane due to the stress undergone while flow sorting.

#### 2.3 Permeabilization of nuclei

Flow sorted nuclei were next permeabilized using the 10x Nuclei Isolation from Complex Tissues for Single Cell Multiome ATAC + Gene Expression protocol (Step 1.2, CG000375 Rev C), with a modified lysis buffer concentration (0.05x). The optimal buffer concentration was chosen to ensure that flow-sorted nuclei were not over-permeabilized or damaged. Post FACS sorting and nuclei permeabilization, nuclei were counted using the Countess II Automated Cell Counter and their concentration adjusted to meet the recommended target of 10,000 nuclei for the Chromium Next GEM Single Cell Multiome ATAC + Gene Expression workflow. The final nuclei suspension was prepared in a chilled diluted nuclei buffer, with the sorted 4x, 2x, and 1x nuclei pooled in a ratio of 50:25:25, respectively. Although we targeted a concentration of *∼*10,000 nuclei in total per sample, our post-sequencing analysis showed variability in nuclei sampled per mouse; this may reflect cell counting errors, clumping during counting, or variation in the proportions of cell types within samples affecting the number of nuclei finally captured. Despite the variability in counts we obtained good quality cells from 32 mice (Supplementary Data 2; see below for exclusion criteria for 4/36 mice).

#### 2.4 Single cell Multiome library generation

Single-cell multiome ATAC and gene expression libraries were generated following the Chromium Next GEM Single Cell Multiome ATAC + Gene Expression User Guide (CG000338) Rev F, in accordance with the manufacturer’s instructions. The resultant libraries underwent size distribution quality checks using the High Sensitivity D5000 Screen Tape Assay (Tapestation 2200, Agilent).

#### 2.5 Sequencing

Libraries were sequenced in five batches (with 4, 4, 4, 12, and 16 animals) using either a NovaSeq 6000 S4 lane, NovaSeq SP Flowcell or NovaSeq X Plus 10 billion lane (Illumina) depending on the number of libraries sequenced at a given time, all employing 150-bp paired-end sequencing. The ATAC-seq and RNA-seq libraries were kept separate but mouse samples were pooled together. The sequencing aimed to target 25,000 read pairs per cell for the ATAC component and 20,000 read pairs per cell for the RNA component. Read lengths recommended by 10x were specified: Gene expression libraries (Read 1: 28 cycles, i7 Index: 10 cycles, i5 index: 10 cycles, and Read 2: 90 cycles), and ATAC libraries (Read 1: 50 cycles, i7 Index: 8 cycles, i5 index: 24 cycles, and Read 2: 49 cycles).

In the final batch, 12 samples (41-1 to 41-12), yielded an insufficient number of reads on initial sequencing on two NovaSeq X Plus 10BN lanes (one each for RNA-seq and ATAC-seq libraries) and were resequenced (ATAC-seq libraries on a NovaSeq X Plus 25BN lane and RNA-seq libraries on a NovaSeq X Plus 10BN lane).

### 3 Processing and Alignment of raw reads

We used the CellRanger-arc (2.0.2) pipeline from 10x Genomics with default parameters and the mm10 reference assembly to de-multiplex and align raw RNA-seq and ATAC-seq reads. This involves a series of steps including trimming adapter sequences from read ends, extracting Barcodes and UMI sequences, and separately mapping ATAC-seq reads with BWA-MEM and RNA-seq reads with the splicing-aware STAR aligner. The output from this pipeline includes sample-level bam files for RNA and ATAC reads, with a unique barcode for all reads from one cell/droplet [3]. The pipeline also generates a number of secondary outputs and data summaries, including cell calls, UMI count matrices and a set of ATAC peaks with the corresponding read counts by Barcode, which we used for quality control purposes.

### 4 Quality control: Choosing high quality cells and filtering reads

We undertook a number of quality control steps to obtain a set of high-quality cells and reads, as described below:

#### 4.1 Delineating cells from empty droplets

Only a subset of sequenced barcodes tag true cell nuclei; most are associated with droplets containing background levels of RNA or DNA. We relied in the first instance on the default cell calling algorithm in cellranger-arc [3]. Briefly, this approach first excludes droplets in which the fraction of ATAC reads overlapping called peak regions (FRiP) is lower than the fraction of genome in peaks (since these cells may have cut sites randomly distributed over the genome), and then uses thresholds on the numbers of ATAC and RNA reads to define cells. A sample-specific threshold that is 10-fold less than the maximum value after removing outliers is defined independently for gene expression and ATAC levels and barcodes above both thresholds are labeled as cells. A K-means algorithm (*k* = 2) is initialized using centroids of cell and non-cell groups to define final boundaries of the cell cluster.

To avoid missing unusual cell populations with low RNA content or distinct chromatin configurations, we also used EmptyDropsMultiome, which statistically models background RNA and ATAC profiles to exclude non-cell droplets and has been shown to work well for germ cell populations [4]. We obtained cell calls using the suggested FDR of 0.1%.

We provisionally classified as cells all droplets that were called as cells by either CellRanger-arc or EmptyDropsMultiome (in our data, *>* 90% of cell calls were shared between the two methods). These provisional cells were then subject to further quality control.

#### 4.2 Using gene expression to assign reference cell types and initial cell ranks

To obtain a biologically informed representation of our gene expression data in a smaller number of dimensions, we mapped the sample-specific gene expression count matrices from cellranger-arc to a published set of 50 reference gene expression components that capture key cellular processes in mouse spermatogenesis [5].

We first normalized our counts data as described in the previous study [5]. Briefly, we removed cells with a total transcript count below 100. We filtered our gene expression data (for a given sample) to genes shared with the reference dataset (*∼*16,800 for all samples). We normalized library size by adjusting counts per cell by the median square root of total transcript counts. Expression values for each gene were then scaled to unit variance (across all the cells in the sample). To mitigate the effects of extreme values, all expression values exceeding a predefined threshold (default: 10) were capped. For each cell in our sample, we then modeled the vector of normalized gene expression as a linear combination of the gene loadings of the reference expression components, and estimated the set of 50 component weights using ordinary least squares.

These component weights were then used to assign a reference cell type and an initial “pseudo-time” to each cell in our sample. Using the 12K cells with assigned cell types and pseudotime ranks that passed QC in the previous study [5] as the reference set, we identified the reference cell closest to each cell in our study (maximizing pairwise cosine similarity between component weights). Each cell in our sample was assigned the cell type and (initially) the pseudotime rank of the nearest reference cell.

#### 4.3 Filtering cells and samples

Starting with all cell-associated barcodes in each sample (excluding non-cell barcodes obtained in 4.1), we used *Seurat 5.1.0* [6] in R for visualization and initial examination of data quality. We excluded cells that met the following criteria: (a) cells with a large fraction of gene expression from mitochondrial genes (*>* 2.5%) or with a non-negligible proportion of (nuclear) ATAC fragments that are mitochondrial (*>* 0.25%); (b) cells with RNA from less than 100 genes or RNA UMI counts in the top 2.5th percentile; (c) cells with an informative (based on pre-called peaks) ATAC read count (nATAC Count) *<* 750 or in the top 2.5th percentile of informative ATAC counts; (d) cells that are extreme outliers in the proportion of gene expression (above 99.8th percentile) from one or more of chromosomes 1 to 19, or the X chromosome.

To examine cell clusters in individual samples and the effects of our quality filters, cells were processed (before and after quality filtering) with the *Seurat* package using default parameters as follows: normalization with *SCTransform*, PCA inference with *RunPCA*, *FindNeighbors* (using the first 50 PCs), *FindClusters*, and finally *RunUMAP* using 50 PCs. We excluded 4 samples from our analyses: two where post-QC cells did not resolve into clear cell populations using either RNA-seq or ATAC-seq, and two where our RNA-seq library preparation failed and yielded very low RNA counts for the majority of cells. We used *DoubletFinder* [7] with default parameters to check for clear doublets based on our gene expression data. Because this flagged a negligible proportion of cells, we also checked for and excluded potential doublets using a different approach based on our ATAC-seq data (see below).

We flagged somatic cell populations using reference cell type designations [5] as well as using cell weights in our data for the somatic components defined in the above study. Cells were flagged as somatic if such components together constituted *≥* 40% of expression. We verified visually that these cell populations clustered separately from the germ cell trajectory in UMAP projections and were well captured by the described markers. Somatic cells were removed prior to estimating Pseudotime ranks. Germ cell stages (defined as groups specified below) were retained and labeled (Fig. 1D) by aggregating reference cell type designations [5] as follows: Spermatogonia (“Differentiating A1-4 Spermatogonia”, “Spermatogonial Stem Cells”, “B Spermatogonia”, “Spermatogonia (Broad)”), (pre)Leptotene, Zygotene, Pachytene (“Pre-Pachytene (& Hormad1)”, “Early Pachytene 1”,“Early Pachytene 2”, “Pachytene”, “Late Pachytene”, “Unknown Pachytene Subset”), Meiotic Divisions, Spermatids (“Round Spermatid”,“Early vs Late Acrosomal”,“Acrosomal”), and Spermiogenesis (“Late Spermiogenesis 1”, “Late Spermiogenesis 2”, “Spermiogenesis”, “Spermiogenesis V2”).

The above steps were all done at the level of individual samples. Across all 32 remaining samples, we collated 203,869 post-QC cells, used in all subsequent analyses.

#### 4.4 Filtering RNA-seq and ATAC-seq reads

We excluded reads corresponding to poor quality cell barcodes (as defined in 4.3). ATAC-seq and RNA-seq reads were processed separately. Although both read types are paired-end, ATAC-seq reads contain genomic sequence (*∼*50bp) on both reads in a pair, and RNA-seq reads only on the second read of the pair (*∼*90bp). For raw aligned ATAC-seq reads in BAM format, we used samtools (samtools view -hf 3 -F 1024 -q 20), to remove PCR duplicates and only retained properly paired reads (both mates in a read pair mapped), with a mapping quality of at least 20. For RNA-seq reads we used a custom 10x tag (xf:i:25) to identify reads that are confidently mapped to the genome/transcriptome, are not duplicates, and are confidently assigned to a genomic feature (gene or other transcribed unit). In both cases, cell barcode information, batch, and sequencing run details, linking reads pooled within samples to individual cells (and the associated gene or other genomic feature where applicable) were extracted from read tags. The filtered sample-level BAM files were then converted to BED format with one read per row, keeping read, feature, and cell-level annotations.

### 5 Phasing reads to their parental chromosome

For both ATAC-seq and RNA-seq reads we next assigned the strain of origin using overlap with ancestry-informative SNPs. To do so, we first obtained lists of good quality bi-allelic SNPs (with B6 as the reference) in PWD and CAST strains (19,976,241 and 19,972,425 SNPs, respectively) from [1]. We combined information at these positions to get a list of 30,615,018 (unique) background informative SNPs genome-wide, annotated with the observed alleles in the B6, CAST, and PWD backgrounds.

To assign reads to one of these three backgrounds, we processed our post-QC filtered BAM files using *samtools mpileup*. Read alignments were queried at informative SNP positions. For each SNP position, this allowed us to generate a list of reads, with the allele at the queried position and its base quality, and the cell barcode associated with each read. To minimize sequencing errors, bases with a quality score below 20 were not used in assigning reads to strains.

For a given read, we compared the variant allele against the list of background informative SNPs to assign it to its strain of origin. At each SNP, an allele could correspond to one or two backgrounds (for example, at a SNP an A allele might indicate the B6 background, and G might be present both in the PWD and CAST backgrounds). Reads where the matching allele was exclusive to one strain were labeled as “confident” (e.g., “conf b6 only”, “conf pwd only”, “conf cast only”). Reads at a SNP shared by two different strains (e.g., CAST and PWD) were classified as “shared” (e.g., “shared cast pwd”, “shared b6 pwd”, “shared b6 cast”). Reads with neither the reference nor alternate allele were labeled as ambiguous.

Where individual reads overlapped multiple SNP positions, we aggregated allele calls across all variant positions covered by a read. We combined information for the two mates in a read pair, since these come from the same DNA molecule. If a read pair overlapped multiple SNPs but all were confidently assigned to the same strain, it was confidently classified. If a read overlapped a set of SNPs with consistent assignments for one or more strains, it was given the classification corresponding to the intersection of these assignments (i.e. the most confident call possible). If different SNPs within a single read supported conflicting strain assignments, the read was marked as ambiguous. Reads that did not overlap with any informative SNPs, or where the only SNP had low base quality (BQ *<* 20) were retained but marked as having an “unknown” strain of origin.

After assigning strain of origin information to ATAC read pairs, we converted read pairs to fragments (i.e., such that the two reads in the pair, oriented appropriately by strand, represent the ends of a variable-length DNA fragment). As is standard, the BED interval of the fragment was obtained using the BAM alignment interval of the sequenced read-pair (using *bedtools bamtobed*), and was adjusted by +4bp from the left-most alignment position and -5bp from the right-most alignment position to account for the transposase cutting DNA with a 9bp overhang on the two strands. BED files containing RNA reads and ATAC fragments and their assigned background for each mouse were then used in downstream analyses. Overall we were able to assign some information about ancestry to on average 50% of our ATAC-seq reads and 33% of RNA-seq reads.

### 6 Calling genomic background in hybrids using ATAC-seq reads

Using ATAC-fragments annotated with the strain of origin, we summarized read classifications for each mouse. We checked that these recapitulated the expected genotypes: in B6 and CAST animals, we observed 0.15% and 1.2% reads with discordant alleles, respectively; in PWD x B6 animals we observed ”conf cast only” reads at a rate of 0.4%; in B6 x CAST animals this number was 0.26% for ”conf pwd only” reads. Of the total number of reads that supported a single strain (e.g., ”conf b6 only”, ”conf pwd only”, ”conf cast only”), PWD x (B6 x CAST) mice had on average 48% ”conf pwd only”, 27% ”conf b6 only”, and 25% ”conf cast only”, with variable proportions of B6 and CAST in individual mice.

We then inferred local ancestry along the genome for each hybrid mouse using a statistical approach. To this end, we binned ATAC reads according to their strain classification (i.e. whether the read supports the PWD, B6, or CAST allele) in 50 kb bins along the genome by animal. PWD x (B6 x CAST) hybrids have a maternal chromosome that is PWD (referred to as the ”pwd homolog”), and a variable amount of B6 and CAST on the paternal homolog (the ”non-pwd homolog”). The procedure for calling local ancestry on the non-pwd homolog is detailed below.

#### 6.1 Hidden Markov Model for smoothing local ancestry calls

We first focused on the reads that confidently support the B6 or CAST background in each 50 kb bin along the paternal genome by animal. We defined confidently callable bins as those with more than 10 reads (i.e., *n*_conf b6 only_ + *n*_conf cast only_ *>* 10) where at least 90% of reads were confidently assigned to a single background (i.e., *n*_conf b6 only_*/*(*n*_conf b6 only_ + *n*_conf cast only_) *>* 0.9 for B6, or *<* 0.1 for CAST). Using only these criteria, we were able to make ancestry calls (”B6” or ”CAST”) for all but 4396 Mb across 23 PWD x (B6 x CAST) animals, i.e., *∼*7% of the genome (*<*200 Mb on average for an animal).

To make ancestry inferences in regions that are difficult to call directly and obtain smoothed ancestry segments genome-wide, we further used a two-state Hidden Markov Model (HMM) to infer the underlying state (“PWD/B6” or “PWD/CAST”) from the observed read counts in each 50 kb bin. Because the fathers are B6 x CAST F1s and F2s, the B6 and CAST ancestry segments are expected to break up only once or twice along a chromosome. We parametrized this with a fixed probability of 0.05% of switching between the two states (i.e., an expected switch rate of once per 100 Mb). We obtained the likelihood of observing a read *R* given either the “PWD/B6” or “PWD/CAST” states, assuming the read is equally likely to sample the PWD or the non-PWD homolog, as follows (with an error rate *ε*). Although the genome wide error rate is much smaller, we set *ε* = 5% to smooth inference over difficult to call segments:

| | $L(R \mid \text{PWD/CAST})$ | $L(R \mid \text{PWD/B6})$ |
| --- | --- | --- |
| conf_b6_only | $\varepsilon$ | 0.5 |
| conf_cast_only | 0.5 | $\varepsilon$ |
| conf_pwd_only | 0.5 | 0.5 |
| shared_b6_pwd | 0.5 | $1 - \varepsilon$ |
| shared_cast_b6 | 0.5 | 0.5 |
| shared_cast_pwd | $1 - \varepsilon$ | 0.5 |

Given a sequence of *K* genomic bins and observed read data for the *k*th bin, the likelihood of the data in bin *k* for state *i* is calculated by multiplying the likelihoods specified over all *n* individual reads:

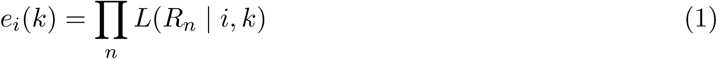

We implemented the forward-backward algorithm with these parameters to obtain the posterior probability of each state in each genomic bin. The algorithm was initialized once per chromosome per animal (with both states initially equally likely). The forward (*α_k_*) and backward (*β_k_*) proba-bilities for the *k*th bin were calculated as:

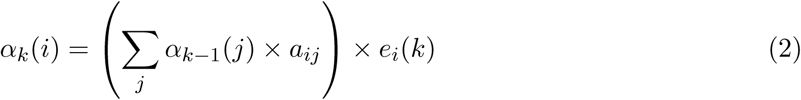

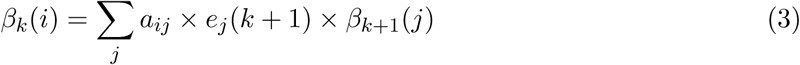

where *a_ij_* is the fixed transition probability (0.9995 if *i* = *j*, 0.0005 if *i ̸*= *j*). We recorded the state with the greater posterior probability for each bin. We masked a subset of regions (20 Mb on average per animal; 0.7% of the genome) with potential mismapping where we suspect the HMM calls may be erroneous. Specifically, we masked discordant segments less than 2 Mb in length with *>* 50% of bins containing no reads, or segments less than 1.5 Mb in length: given that we expect 1-2 switches per chromosome in these animals, segments of this size are highly likely to be artifacts. Masked segments were imputed as possessing the background inferred at either end of the masked region if both ends were identical, and remained uncalled otherwise. Post-smoothing (see Supplementary Fig. 4 for an example), we were able to reclassify almost all previously ambiguous regions with less than 2 Mb remaining uncalled on average per animal; smoothed ancestry calls were consistent with our initial calls in confidently callable regions 99.7% of the time. For PWD x B6 mice, our approach correctly inferred the PWD/B6 background in 100% of bins. On average, on chromosomes other than chromosome 17, we get 53% B6 and 47% CAST ancestry, consistent with B6 x CAST F1 or F2 fathers; because we chose animals which specifically have PWD and B6 *Prdm9* alleles, the *Prdm*9-containing chromosome 17 has 94% B6 ancestry on average across animals. Of 437 total autosomes in 23 animals, 48% showed one background switch, 12% showed two, and 3% showed 3; an average of 15 per animal. Among only animals with B6 x CAST F1 fathers, this number was 11 for all chromosomes (representing crossovers between the B6 and CAST homologs in the father’s germline, and consistent with reported rates [8]); 57% of chromosomes had 1 and only 0.6% had two background switches. Overall, these checks provide substantial confidence that our inferred ancestry segments have high accuracy.

#### 6.2 Calling LD blocks across animals

We used the local ancestry calls to determine the boundaries of shared ancestry segments across PWD x (B6 x CAST) animals, which yielded 324 blocks of distinct (though not independent) ancestry patterns genome-wide, with no ancestry change within each block in any animal. We excluded 38 blocks that remained unclassified after the above procedure in *>* 5 animals or were shorter than 200 Kb, and which span 2.3 Mb of the genome in each animal in total, leaving 286 blocks.

#### 6.3 Re-assigning allelic information to reads using local background information

As described above, we obtained a strain of origin for each read using overlap with ancestry informative SNPs, and used this to call local background. Using this inferred local background, we next re-assigned allelic origin to reads: this allowed us to assign a larger number of reads with alleles shared between strains, and to avoid spurious assignments based on individual erroneous SNPs.

Specifically, in PWD x (B6 x CAST) mice, given animal-level background along the genome (PWD/B6 or PWD/CAST), we classified reads by homolog: where the local ancestry was PWD/B6, for instance, reads could be classified into (1) those that came from the B6 homolog (”conf b6 only” or ”shared b6 cast”), (2) those that came from the PWD homolog (e.g., given B6 on the non-PWD homolog, reads that were classified as being either ”conf pwd only” or ”shared pwd cast” were counted towards the PWD homolog), (3) those that carry an allele shared in the two backgrounds and could come from either, and (4) those that had no allelic information and/or were inconsistent with the inferred background. The same approach was used for PWD x B6 mice, where the background is always PWD/B6. In B6 x CAST mice, where the background is always B6/CAST, reads can similarly be partitioned by homolog except when carrying alleles shared by CAST and B6. Using this approach, we obtained the number of fragments stratified by background and homolog in 50 kb windows along the genome in each cell, used as input to the cell caller (see below). The same data were used to obtain allele-specific fragment counts in hotspots (to examine allele-specific binding of PRDM9) and the same approach was used to obtain allele-specific RNA counts to identify allele-specific expression (see below).

### 7 Calling homolog-aware copy number state along the genome in individual cells

The number of ATAC reads in a cell in the germline reflects both chromatin accessibility and also the DNA copy number of the cell stage. Meiosis involves a whole genome duplication followed by two successive divisions. Each locus thus starts out as diploid (2n, i.e., 2 copies of each chromosome), transitions to 4n in early meiosis, then to diploid (2n) after the first meiotic division, with the segregation of homologous chromosomes, and finally to a haploid (n) state after the second meiotic division.

Before beginning, we note that (i) it is difficult or impossible even in principle to distinguish the early 2n and 4n states (5 and 11 in the below table), which only differ in terms of a doubling of DNA content genome-wide, because inter-cell variability in total ATAC-seq counts is high and so we condition on the total number of ATAC-seq reads in each cell, and (ii) our aim in this analysis is therefore *not* to generate a fully accurate inference of copy number at each position along the genome, which is difficult in any individual cell to verify and might be impacted e.g. by broad patterns of accessibility that might vary from cell-to-cell given we are using ATAC-seq data. Rather, we seek to generate measures that are (demonstrably) informative for downstream analyses, e.g. to identify cells that have undergone the MI and MII divisions, or have abnormal properties; see section 8.

Given data (see section 6) for each cell and 50kb bins giving the total number of reads assigned to each parental background (“PWD” vs “non-PWD” backgrounds, with the latter always “B6” or “CAST”), we again used a Hidden Markov Model to infer the maximum *a posteriori* copy number state in 50kb windows along the genome, now in individual cells. For B6 *×* CAST F1 animals, we instead use B6 and CAST as the two homolog backgrounds but otherwise proceed identically.

We defined 13 copy number states as follows, by considering all possible states that are either expected as meiosis progresses or which differ by *±*1 chromosome from an expected state:

| State | 1 | 2 | 3 | 4 | 5 | 6 | 7 | 8 | 9 | 10 | 11 | 12 | 13 |
| --- | --- | --- | --- | --- | --- | --- | --- | --- | --- | --- | --- | --- | --- |
| PWD homolog | 0 | 1 | 0 | 0 | 1 | 2 | 3 | 2 | 1 | 0 | 2 | 3 | 2 |
| Non-PWD homolog | 0 | 0 | 1 | 2 | 1 | 0 | 0 | 1 | 2 | 3 | 2 | 2 | 3 |
| Group | hom | hom | hom | hom | het | hom | hom | het | het | hom | het | het | het |

For each cell, our approach aims to estimate a vector *p* whose entries *p_i_* are the prior probabilities of each state in that cell (which is within in some particular animal), and then from these a posterior probability matrix *P* , which depends on *p*, such that *P_ij_*is the probability that bin *j* has state *i* in this cell. We do this under a model where (i) transitions occur once every 100Mb on average at boundaries between 50kb bins, so with probability 0.0005 between successive bins, and each time a transition occurs a new state is sampled according to *p*, (ii) within each 50kb bin the expected number of informative reads per chromosome copy on the PWD/non-PWD backgrounds is known up to a constant of proportionality. In practice, we estimate this from the sum of all such reads across cells within the animal, because we expect the overall number of PWD/non-PWD copies is equal after averaging over thousands of cells; the main source of variability is across bins rather than backgrounds, because different bins contain differing numbers of informative markers. (iii) reads have a 95% probability of correct assignment to the PWD/non-PWD background and a 5% probability of misassignment.

Given these assumptions, we model the number of reads from each bin on each background as approximately Poisson-distributed (see below), because there are many bins genome-wide so the probability of a read coming from any particular bin is small. This results in a hidden-Markov-model structure with Poisson emission probabilities, conditional on the underlying states detailing the number of PWD/non-PWD copies.

The transition parameter is chosen to approximately mimic the recombination rate genome-wide for mice, to allow analysis of MI/MII cells (in other cells it functions as a strong prior against copy number changes), while the misassignment parameter (5%) is likely conservative but aims to guard against false inference of copy number changes along the genome. We believe our model is plausible, but we validate its usefulness in practice using other measures e.g. gene expression in the same cells, comparison with crossover detection using an alternative approach, and comparisons across chromosomes. The posterior probabilities are then used in downstream chromosome-level and cell-level classifications (e.g. identifying MI and MII cells as well as abnormal cells).

For parameter inference we iteratively update *p* and *P* to convergence. The approach is motivated by an (approximate) EM algorithm to estimate *p* which is formally valid if our emission probabilities were *fixed*. In this EM approach, we consider as missing data the underlying copy-number state vectors along the genome. By standard theory, given *p*, we can then obtain via the forward-backward algorithm the expected posterior number of genome-wide number of transitions *t_i_* into state *i* for each *i*. In turn, this allows, by standard theory, given an initial inference of *p_i_*for each *i* an EM update whereby if, at iteration *t* we have current probabilities *p^t^* of each state, in iteration *t* + 1 we obtain

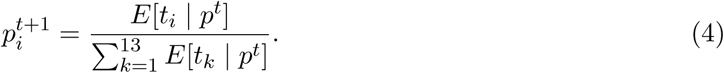

In practice, and given we are using other approximations we update using the (very similar) observed fraction of the genome in each state rather than the number of switches, because this is efficiently calculated elsewhere in our algorithm.

In our setting, the above EM algorithm is not precisely correct, because our emission probabilities depend on both the underlying parameters, but also on the hidden states, so that the expected log-likelihood is not easily precisely calculable. However in practice, we can approximately solve this by using the fact that the genome is large and so the observed hidden state samples are expected to be very close to those expected from their average values. We then expect the resulting emissions to similarly be very close to those were we to know the underlying hidden states. Therefore, at each iteration approximate emission probabilities are calculated from the previous iterations’ expected values for the hidden states at each locus.

To implement this note that for a particular animal, the number of informative PWD and non-PWD reads varies among bins due to local SNP density (and presence of B6/CAST segments varying by animal mean this impact is animal-specific). Given a very large number of cells in each animal generating (by the WLLN) a very close overall 50% balance of PWD/non-PWD chromosomes, we capture this impact by the total number of reads in each bin *j* for this animal: *t_j_*_1_ and *t_j_*_2_ for PWD and non-PWD reads respectively. This acts as a prior expectation of the number of informative reads of bin *j* for each background. Then suppose in a particular individual cell, state *i* has *n_i_*_1_ copies of the PWD genome and *n_i_*_2_ copies of the non-PWD genome. The probability a given read comes from this bin and the PWD background is then proportional to *n_i_*_1_ *× t_j_*_1_ and similarly for the non-PWD background.

At a given iteration, suppose our cell has a (genome-wide) probability matrix *P* such that *P_ij_* is the (current iteration) posterior probability that bin *j* has state *i* in this cell. Then from the above, conditional on bin *j* possessing state *i*, the probability that a random read originates from that bin and the PWD background is modelled as:

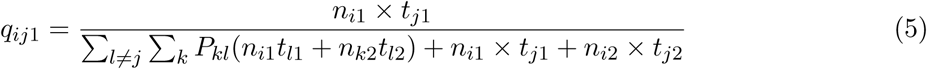

and similarly we may define *q_ij_*_2_ for the non-PWD background. In practice, we calculate these *q_ij_* by summing the denominator over all *l*, because the impact of the single bin *j* is negligible.

Now if for a given cell, *N* is the total reads in that cell, the number of reads *r_j_*_1_ matching the PWD background has a Poisson distribution with mean:

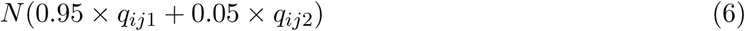

while the number of reads *r_j_*_2_ matching the non-PWD background has a Poisson distribution with mean:

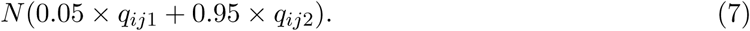

Using these distributions and modeling reads in each bin as independent allows us to calculate emission probabilities *e_ij_*_1_ and *e_ij_*_2_ and overall emission probability *e_ij_*= *e_ij_*_1_*e_ij_*_2_ conditional on the hidden states for each bin, and we can obtain posterior inferred probabilities for each state for each bin along the genome via the standard forward-backward algorithm as described above.

Finally, our algorithm then proceeds as follows:

- Initialise 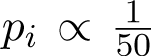 for each *i* except 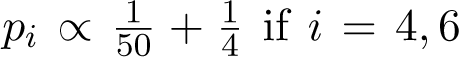 and 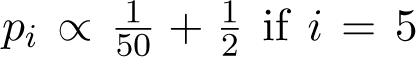. This initially allows all states but with an increased likelihood of the “null” diploid cell states as well as homozygous states expected after the MI and MII divisions. (In practice, we observed convergence of *p* vectors away from these initial values.)
- At a given iteration *t* with current parameters *p^t^*, run the forward-backward algorithm to obtain *P^t^*, the current state probability matrix with entry *P^t^* is the posterior probability of state *i* in bin *j*. Then update the parameters:

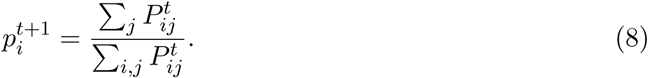

- Iterate step 2 until convergence, defined as a maximum change in parameters *p* between successive iterations of *<* 0.0001 (in our setting, 100 iterations were always sufficient).

### 8 Calling cell classes

We next summarized the genome wide information from the HMM for each cell in our hybrid mouse samples, with the aim of classifying cells into meiotic cell types, including those that are likely pre-division, and those that have undergone the first or second meiotic divisions). For this, we note that chromosomes in cells that have not undergone a meiotic division (i.e., spermatogonia and primary spermatocytes) are expected to be heterozygous throughout the genome because the homologous chromosomes are from different parental subspecies; similarly chromosomes in cells that have undergone both divisions are haploid and homozygous throughout the genome (see Fig. 1E). In cells that have only undergone the first meiotic division, which separates homologs, chromosomes are expected to mainly have homozygous centromeres and heterozygous telomeres due to crossing over between homologs followed by segregation of homologous centromeres into separate daughter cells, except rarely when e.g. the crossover has occurred very close to the centromeric or telomeric end and is therefore undetected. Because the prior on our single-cell state-HMM is uniform, the extent to which cells divide into groups with these properties also provides a powerful test of the accuracy of our inference procedure in practice (see Fig. 1F).

We also identified cells carrying deletions or duplications of one or more chromosomes, and other cells with a potentially abnormal or ambiguous genomic state. Our overall approach was to first make a consensus call for each chromosome based on the inferred states in 50 kb bins (as described above), and then use these chromosome-level calls jointly to classify cells.

#### 8.1 Classifying chromosome types

The chromosomal calls were made as follows:

| Classification | Allowed states (H1:H2) | Criteria |
| --- | --- | --- |
| **Normal chromosome states** |  |  |
| Heterozygous | Both homologs present: 1 : 1, 2 : 1, 1 : 2, 2 : 2, 3 : 2, 2 : 3 | $\leq 10$ bins in any homozygous state |
| Meiosis-I recombinant | Homozygous: 1 : 0, 0 : 1, 2 : 0, 0 : 2, 3 : 0, 0 : 3, 0 : 0. Heterozygous: 1 : 1, 2 : 1, 1 : 2, 2 : 2, 3 : 2, 2 : 3 | Homozygous and heterozygous blocks with transition at crossover site. $> 10$ bins homozygous and $> 10$ bins heterozygous; first bin homozygous |
| Homozygous (homolog 1) | 1 : 0, 2 : 0, 3 : 0 (partial 0 : 0 tolerated) | $\leq 10$ heterozygous bins, $> 10$ homolog-1-only bins, $\leq 10$ homolog-2-only bins |
| Homozygous (homolog 2) | 0 : 1, 0 : 2, 0 : 3 (partial 0 : 0 tolerated) | $\leq 10$ heterozygous bins, $\leq 10$ homolog-1-only bins, $> 10$ homolog-2-only bins |
| Meiosis-II recombinant | Homozygous throughout but switches parental origin at crossover point(s): 1 : 0, 0 : 1, 2 : 0, 0 : 2, 3 : 0, 0 : 3 (partial 0 : 0 tolerated) | $\leq 10$ heterozygous bins, $> 10$ homolog-1-only bins, $> 10$ homolog-2-only bins |
| **Aberrant chromosome states** |  |  |
| Aberrant recombinant | Homozygous: 1 : 0, 0 : 1, 2 : 0, 0 : 2, 3 : 0, 0 : 3, 0 : 0. Heterozygous: 1 : 1, 2 : 1, 1 : 2, 2 : 2, 3 : 2, 2 : 3 | Meiosis-I recombinant but first bin heterozygous |
| High copy number recombinant | Homozygous: 3 : 0, 0 : 3. Heterozygous: 2 : 1, 1 : 2 | $> 10$ bins homozygous and $> 10$ bins heterozygous; $> 90\%$ of chromosome |
| Unbalanced heterozygous | 2 : 1, 1 : 2 | $> 90\%$ of chromosome |
| Aberrant (high copy number) heterozygous | 3 : 2, 2 : 3 | $> 90\%$ of chromosome |
| Chromosomal deletion | Both homologs absent: 0 : 0 | $> 90\%$ of chromosome |

#### 8.2 Classifying cell types

The main meiotic cell types are determined as follows:

- Cells with all 19 autosomes in a (normal) heterozygous state are classified as undivided, or

*Normal*.

- Cells with all 19 autosomes homozygous (Meiosis-II recombinant or Homozygous a1 or a2) are confidently classified as having gone through two meiotic divisions (and labeled *Meiosis-II or MII*).
- Cells with *>* 17 of 19 autosomes possessing both a homozygous centromere and showing a non-centromeric heterozygous segment indicating a crossover are confidently classified as *Meiosis-I/MI* (up to two chromosomes are allowed to be fully homozygous, suggesting a telomeric crossover).

We further classified some cells less stringently as *Mainly Normal*, *Mainly MI*, and *Mainly MII* if they carry at least 14/19 chromosomes that are heterozygous, Meiosis-I recombinants, and normal Meiosis-II chromosome types (Meiosis-II recombinant or Homozygous a1 or a2), respectively, but have other chromosomes that do not meet these criteria, potentially as a result of erroneous calls or varying chromatin accessibility, aberrant recombination, failure to recombine, or potential mis-segregation during cell division: we only allow for this relatively small number of aberrant chromosomes in these categories, to avoid being misled by potential doublets.

Normal (undivided) cells that are actively replicating their DNA can contain large numbers of heterozygous chromosomes with unbalanced homolog copies. Among cells where at least 95% of genome-wide read counts were assigned to heterozygous states (states 1:1, 1:2, 2:1, and 2:2), we calculated the proportion of 50 kb bins with unbalanced homolog states (1:2 or 2:1) genome-wide, and flagged cells as likely replicating if more than 33% of the genome showed this unbalanced state (see Fig. 1G). While heterozygous chromosomes with the two homologs unbalanced are not aberrant *per se*, we re-classified (see below) undivided cells if the majority (*>* 14) of chromosomes are in an extremely unbalanced state (*>* 90% unbalanced), as these are more likely to represent potential doublets than replicating cells.

Of all cells in our hybrids, 65% are classified as pre-division, 15.2% have completed both meiotic divisions, and 2.4% have completed only the first division (see Supplementary Fig. 5 for a summary of the inferred genome-wide states). We note that the relative proportions of these cell types may not reflect the actual distribution of germline cells, as our experiment enriches for spermatocytes over sperm, and may affect other sub-populations of cells. Our criteria capture the majority of our cells, but 17.4% of cells fall outside of all the above (stringent) criteria by possessing a mixture of inferred states (see Supplementary Fig. 5), despite passing our basic cell quality filters. A subset might be potential heterotypic doublets, e.g., between haploid cells and other cell types, if they have much of their genome in 3 or 5 copies, or a large number of aberrant chromosomes. We excluded such unclassified (“unknown”) cells in our analyses of post-division cells (including calling crossovers and aneuploidy), but included them elsewhere (they do not impact our conclusions).

### 9 Pseudotime inference

As described above in 4.2, we first obtained an initial pseudotime assignment for each cell using a previous study of single cell gene expression in mouse testes [5]. To resolve ties between cells, we initially randomly ranked cells assigned the same initial value. Because these ranks obviously do not fully represent properties of expression in our system, and in particular are noisy and based on a smaller prior dataset with many ties, we refined these ranks in the following steps: (a) we obtained a set of gene regulatory profiles through time using an iterative Gibbs sampling algorithm using the initial ordering of cells, and (b) we smoothed loadings of regulatory components through time and used linear interpolation, to give a final unique rank to each cell.

#### 9.1 Calling gene expression time profiles

To more precisely capture the key time trends in gene expression in our data, we modeled gene expression as arising from a mixture of *K* (=75) regulatory components each with a particular profile through time. We started with per gene read counts in individual cells ordered by initial values of pseudotime, and binned into *T* = 20 pseudotime bins. We used an iterative Gibbs sampling algorithm with Dirichlet priors to factorize gene expression through time (*g* genes *× T* time bins) into gene loadings on components (*g* genes *× K* components) and component time profiles (*K* components *× T* time bins), encouraging gene loadings to be sparse (i.e., we expect that a given gene would typically be subject to a small number of regulatory programs or transcription factors). Our model is that each component has a loading for each gene that dictates the overall fraction of reads from that component for that gene. Each component also has a loading for each time, which dictates the “shape” of that component (so we fit 75 shapes in total), i.e. the probability that a read from this component comes from each of the 20 (pseudo)time bins. Then each read samples a component, gene and pseudotime independently of other reads, yielding a Multinomial distribution on read counts in each such category conditional on the total number of reads. Having observed data, we can conceive of complete data where as well as the (directly) observed gene and time information for each read, we also observe the component from which a read derives.

This allows us to use a standard Gibbs sampling approach to iteratively update gene and time loadings for each component, with each having a Dirichlet (conjugate) prior. We initialized component time profiles by averaging expression across genes at time *t* so all components start with the same time profile through time (i.e., a vector *x*_1*j*_*, . . . , x_tj_* for the *j*th component). Gene loadings (*l_ij_* for the *i*th gene and *j*th component) were initialized uniformly (= 1*/K* for *K* components). We then iteratively (over 200 iterations) assigned reads to components given current parameters, and re-estimated parameters based on these assignments. The log-likelihood of the observed data with the current parameters was calculated at each step to monitor convergence, which occurred in *<* 100 iterations.

Specifically, in the E-step, we probabilistically assigned each gene’s reads at each time (*r_it_*) to the *K* components. For gene *i* at time *t*, the probability that a read came from component *j* (*p_ij_*) is proportional to the gene’s loading on component *j* times component *j*’s activity at time *t*.

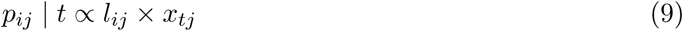

We sampled this distribution, yielding *n_ijt_* reads from gene *i* assigned to the *j*th component summed across all genes at a given time *t*, or across all time bins at a given gene, and we then obtain overall counts:

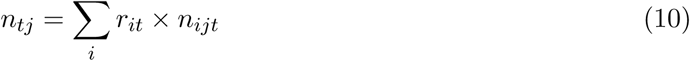

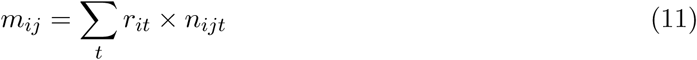

Given the read assignments, we updated gene loadings and component activities through time by sampling from Dirichlet posterior distributions, conjugate to the prior distributions. Posterior component time profiles and gene loadings were then sampled respectively, as:

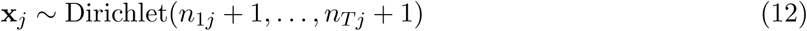

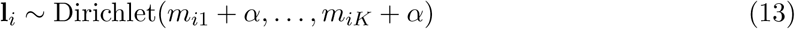

*α* is a sparsity term which was initialized as 0.1 for all *K* components (favoring sparse gene loadings) but updated for each component in each iteration as the optimal degree of sparsity for each component was learned from the data. We did not place a prior on *α*; instead the updates were done using penalized maximum likelihood with Newton-Raphson optimization to tune *α* to the appropriate level of sparsity given the (average) gene loadings by component observed in each iteration.

#### 9.2 Smoothing and interpolation procedure

To use the inferred components to order cells, we multiplied gene loadings per component (genes *×* components) by normalized gene expression in each cell (genes *×* cells) to calculate component activities by cell. The resulting matrix of 75 component weights per cell, with cells ordered by their initial pseudotime ranks, was used as the input to the smoothing algorithm. The aim is to start with the initial pseudotime ranking (which assigns many cells the same value based on expression) and smooth this using only the components, so that gene expression varies maximally “smoothly” through time, once we order cells by pseudotime, with some noise (which we fit as a multivariate Gaussian; see below).

First, we sampled a random subset of 2000 cells. For all other cells, we identified the nearest (in euclidean distance) of these 2000 cells based on the component weights. For the 2000 cells, we now obtained a pseudotime for each cell based on the median initial pseudotime rank of those cells in the larger dataset for which it is the nearest neighbour. We also replaced the actual component weights for each cell with the mean of weights among the component cells for which it is nearest neighbour, resulting in 2000 ordered “grid point” values with times and component values, each representing an average over many (around 1000) similar cells. To now reorder these to ensure smooth expression changes through time, we first took each of the 75 expression components, predicted their expression through time at the 2000 grid points using the grid component weights, and then smoothed by fitting an optimal piecewise linear function allowing up to six breakpoints to the initial fitted expression through time. Next, we obtained a modified pseudotime using these piecewise smooth functions by defining 100 evenly-spaced checkpoints along the smoothed cell trajectory and for all successive points, calculated the interpolation parameter *λ ∈* [0, 1] that minimized squared Euclidean distance between the cell’s observed component values and the predicted value given *λ* were the cell placed at a position a fraction *λ* between the points:

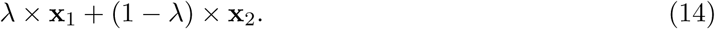

We assigned each cell (in the full set) to the checkpoint interval and lambda value with minimum error, which then reorders cells and redefines pseudotime - reclassifying cells that were originally in a sub-optimal location along the trajectory.

To adjust for the fact that some of our component weights are strongly correlated (because some components are very similar), we re-sampled a large number (10,000, for computational efficiency) of newly re-ordered cells, and estimated the (*K × K*) covariance matrix of the residuals (i.e., the observed component values - their predicted values given the fitted *λ*). Using the same 100 check-points as above, we repeated the previous step of finding optimal *λ* values between checkpoint pairs, now minimizing covariance-weighted rather than Euclidean distance, for a final set of pseudotime values. The covariance weighting accounts for the correlation structure among components, penalizing deviations that are unlikely. Finally, we ordered cells by their predicted positions and assigned ranks 1:N for N ordered cells: these were used as Pseudotime ranks in subsequent analyses. We verified these using expression patterns of marker genes, and separately using the ATAC-seq data (as described elsewhere), noting that the ATAC-seq data is not used for our pseudotime inference so is useful for validation.

### 10 Marker gene expression

We downloaded a list of standard mouse genes from Ensembl (using *EnsDb.Mmusculus.v79* in *R*). We limited our UMI count matrix to genes in the above list. We summed across genes within a cell to get total UMI counts per cell.

To examine *Prdm9* expression through spermatogenesis, we ordered cells by pseudotime and grouped them into bins of equal size (5,000 cells). We summed *Prdm9* UMI counts by bin, and divided by the total expression for all cells in the bin to obtain the proportion of RNA reads in a pseudotime bin that come from *Prdm9*. For other selected genes, we similarly calculated expression (relative to total expression) in cells binned by Pseudotime (and genotype).

### 11 Chromatin accessibility in PRDM9 Hotspots

We obtained a previously published list of PRDM9-binding hotspot locations in PWD *×* B6 males, along with information on DMC1-binding heat, H3K4Me3 signal, PRDM9 motif presence and position, and other hotspot-specific information [1]. We included hotspots that had *>*95% posterior probability of containing a true PRDM9-binding motif (either the B6 or PWD motif), obtaining a final list of 12,175 PRDM9 hotspots. We defined the center of the motif as the center of the hotspot, and tiled the hotspots (center *±* 1kb) in fixed windows of 25bp with the motif at the center of a 25bp window.

To calculate average chromatin accessibility in all PRDM9-binding motifs through meiosis, we ordered cells by pseudotime and grouped them into bins of 5,000 cells. Within each pseudotime bin, we counted the total number of sub-nucleosomal ATAC fragments (length *≥* 30bp and *<* 120bp) in the central 25 bp windows corresponding to 12,175 hotspot centers, and divided by the total number of such fragments genome-wide (in the same pseudotime bin). We used fragment midpoints to ascertain fragment overlap with the genomic windows, as these can indicate both the location of nucleosome-free regions and nucleosome positioning (but obtained the same results using the distribution of fragment ends to identify the nucleosome-free region).

To examine local chromatin patterns around hotspot centers, we identified the peak time of PRDM9-binding and for that pseudotime, calculated the genome-wide proportion of sub-nucleosomal ATAC fragments that likely arise from transposase cutting in nucleosome-free regions in 25 bp windows in *±* 1 Kb around the hotspot centers (averaging within these bins across all hotspots). To examine the distribution of nucleosomes, we calculated the genome-wide proportion of fragments likely corresponding to single nucleosomes (length *≥* 150 bp and *<* 250 bp) in the same genomic windows (again averaged in each window over all hotspots).

As described above (see 6.3), we assigned background information to each ATAC fragment that maps to hotspot regions (i.e., whether the fragment came from the PWD vs non-PWD homolog, and if non-PWD whether it was B6 or CAST in origin). For each hotspot we then counted the number of fragments, stratified by background and homolog, combining data from all PWD *×* (B6 *×* CAST) animals. Homolog-specific binding in the PWD/B6 (or PWD/CAST) background was calculated as the proportion of sub-nucleosomal ATAC fragments from each homolog at hotspot centers (in 25 bp windows centered on motif centers) for individual hotspots controlled by the PWD or B6 *Prdm9* alleles. Average homolog-specific binding for each type of hotspot in the two backgrounds was calculated by weighting the homolog-specific proportions by hotspot heat (calculated for a given hotspot as the fraction of all short fragments, including those without background information, over 12,175 hotspots that are in that hotspot; alternative measures of hotspot heat produced similar results).

Each hotspot was with a previously published measure (using H3k4me3 ChIP-seq) of PRDM9 binding asymmetry in PWD *×* B6 mice [1]. We compared this with our chromatin accessibility based measure for individual hotspots in PWD/B6 regions in PWD *×* (B6 *×* CAST) animals. We also examined homolog-specific chromatin accessibility around hotspot centers in PWD/B6 regions in PWD *×* (B6 *×* CAST) animals, averaged over hotspots, for strongly asymmetric hotspots with either *<*10% or *>*90% H3K4me3 signal from the B6 (vs. PWD) homolog. We used 25 bp windows, and counted ATAC-seq reads from each homolog in each genomic window in the hotspot region, normalizing these counts by the total number of informative (i.e., about either homolog) reads across the entire hotspot region (*±*1 Kb around the hotspot center). Note that we only used reads from the Pseudotime bins (of 5000 cells each) corresponding to cells with substantial PRDM9 binding at hotspots (in this case the peak of binding *±*1 Pseudotime bin)

### 12 Estimating Asynapsis/Silencing levels

We developed a pipeline to infer animal and chromosome-specific silencing probabilities as parameters in a statistical model, and then used the values of these probabilities to study properties of silencing, asynapsis and fertility in our mice.

#### 12.1 Per chromosome expression counts in single cells

We first recorded the number of RNA reads (UMIs) by chromosome and cell (by animal). We excluded cells with fewer than 400 total reads from the estimation procedure. Read counts per chromosome per cell, as well as animal ID and genotype, and the pseudotime rank for each cell were used as input to our pipeline below.

#### 12.2 Modeling silencing parameters

This model estimates the underlying fraction of RNA reads in a given cell drawn from a particular chromosome. It takes into account several issues: first, there is sampling noise around this fraction, which we model using a multinomial distribution. Secondly, there can be systematic variation among cells, and through Pseudotime as well as across mice due to e.g. higher expression of particular genes on a chromosome, and this leads to overdispersion of this proportion vs. expectations under the multinomial model. Finally, we note that even “silenced” chromosomes still produce some expression, and so we threshold “silencing” to be the setting where the underlying value for a chromosome is *<*75% of the distribution expected under a null model.

We fit the model to individual chromosomes. Thus the input data relating to cell *i* and chromosome *c* is simply the total autosomal RNA reads for that cell, *n_ai_*, and the number *n_aic_* originating from chromosome *c*. Assuming reads are independent conditional on some overall fraction of sequenceable transcripts *p_ic_* coming from this chromosome, *n_aic_*has a binomial distribution with parameters *n_ai_* and *p_ic_*. For the reasons above it is important to allow *p_ic_* to depend on *i*, i.e. vary among cells. A natural (conjugate) prior distribution is that at time *t*, *p_ic_* has a beta distribution with parameters *α_ct_* and *β_ct_* which can depend on *t* and *c*. Under this prior, we may integrate out the prior to give the likelihood for a single cell:

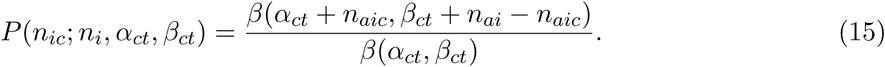

This likelihood multiplies across cells *i* to produce an overall likelihood.

We extend this model to allow for variability among animals *a* as follows. We introduce a vector of 51 multipliers *f* = *c*(0, 0.025, 0.05*, . . . ,* 1, 1.05*, . . .* 1.25), whose *j*th value *f_j_* is associated with a probability *p_acj_*, representing the probability the underlying multiplier takes this value (note this parameter depends in general on animal and chromosome), with Σ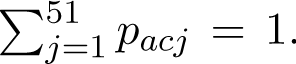 *p_acj_* = 1. We model the actual proportion *p_ic_* as being now drawn from a mixture of 51 beta distributions, with the *j*th distribution chosen with probability *p_acj_*and having parameters *f_j_α_ct_* and *β_ct_*, i.e. multiplying the first parameter so as to reduce/increase the fraction of reads from chromosome *c* accordingly. We can write down the likelihood of the data for a cell under this model:

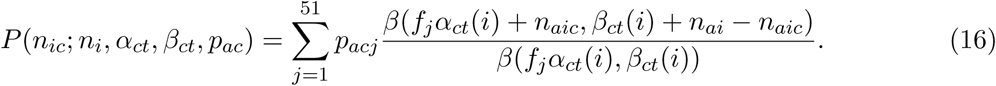

The overall likelihood then simply multiplies across cells and animals.

By selecting the parameters *p_acj_*appropriately we can clearly fit a near-arbitrary set of prior proportions, and so we constrain the fit and model space as follows. Firstly, we fix parameters *α_ct_* and *β_ct_* across animals and fit their distributions using the canonical B6 mouse, as a common reference point. Secondly, we allow arbitrary mouse- and chromosome-specific values for values of *j* from 31 to 51, i.e. multipliers (0.75*, . . . ,* 1, 1.05*, . . .* 1.25) within 25% of the B6 mouse – we regard these as all coming from the appropriate null distribution for a particular mouse and chromosome, and note we typically have a large number of cells to fit these distributions. Finally, we regard - indeed, define - *silencing* as comprising those reads drawn from multipliers (0.0*, . . . ,* 0.725) with a larger reduction in underlying expression. Although this definition is slightly arbitrary, we found it to work well in practice in separating animals with substantial (visible) silencing from e.g. homozygous B6 and CAST animals where in almost all cases we observed expression within the “normal” range for all chromosomes. Thus, one could regard the 25% range as encompassing the spread observed in such animals. Note that by using *α_ct_* and *β_ct_*, we account for a base level of natural “noise” in the observations, so that even cells with fewer reads from a given chromosome may not be inferred as having silencing of that chromosome, unless other cells support that this occurs and the observed fraction is low enough to suggest it lies outside the range expected from noise. The parameters *p_acj_* act to adjust this noise for *j* = 31, 32*, . . . ,* 51 to be specific to a certain animal.

Because we see fewer cells with evidence of silenced expression, we constrain the fit by setting *p_acj_* = *S_ac_p_cj_* for *j* = 1, 2*, . . . ,* 30, where 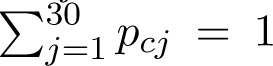 *p_cj_* = 1, so that animals share a common silencing profile for a given chromosome, but have an animal- and chromosome-specific parameter *S_ac_* representing the probability of silencing for this animal and chromosome. Ultimately, for many downstream applications we are interested in the values of *S_ac_*.

At the heart of our model-fitting we use an expectation maximization procedure to find maximum likelihood estimates of parameters *S_ac_* (for 19 chromosomes *c* and all animals *a*), *p_cj_* for 30 values *j* of the extent of silencing and 19 chromosomes, and *p_acj_* for 21 values *j* of the extent of normal variation without silencing, 19 chromosomes and all animals *a*, as well as the “base” parameters *α_ct_*and *β_ct_*.

##### Initial model fitting (shared null distribution across animals)

We first fit *α_ct_* and *β_ct_* using 41 time bins (based on equal spaced pseudotime blocks corresponding to 10,000 cells per block, sliding 5,000 cells each time. For the block *t* we took all reads from chromosome *c* in B6 mice, and used the “dirmult” R package with input data the number of expression reads in each such cell from each autosome to select values maximizing the likelihood (15). We fixed the resulting *α_ct_* and *β_ct_* values for the remainder of our modeling. A cell is assigned to whichever of the 41 time bins has centre is closest to the pseudotime for that cell, and uses the corresponding *α_ct_* and *β_ct_* values. In downstream analyses, we define pachytene cells as being those in time bins corresponding to times later than those with strong *Prdm9* expression and binding, and earlier than those of cell divisions, but also encompassing the time range where we observe strong departures from the null models defined above in some mice and some cells (time bins 8-30, of 41). Note that the ratio term on the RHS of equation (16) is then independent of the parameters we still must fit, so can be precalculated and viewed as constant. For a given chromosome and cell, we calculate these terms and then fit an initial model of the form *p_acj_* = *S_ac_p_cj_* for *j* = 1, 2*, . . . ,* 30, and *p_acj_* = (1 *− S_ac_*)*p_cj_* for *j* = 31, 32*, . . . ,* 51 and constrained so that 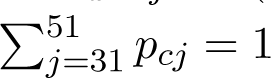 and also 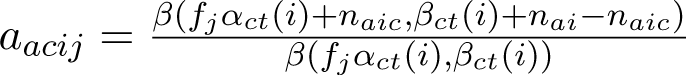 *p_cj_* = 1. This differs from our final target model by having a shared and so more constrained null distribution (*j* = 31, 32*, . . . ,* 51) among animals.

To fit this model we develop an Expectation-maximisation (EM) algorithm by introducing an indicator variable *I_acij_* which equals 1 if mixture component *j* is used for cell *i* from animal *a* on chromosome *c* and 0 otherwise, so that the combination of observed data and these additional variables constitutes “full” data. Note that the full data log-likelihood is then

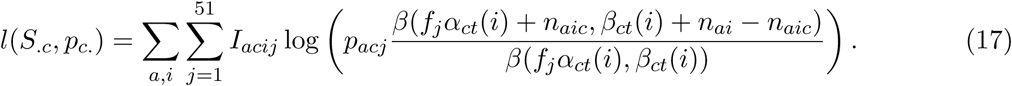

The EM algorithm proceeds by maximizing the expected-log-likelihood over the posterior distribution given the current parameters. Noting part of the right hand log term is a fixed constant so does not impact the maximization, up to a constant this expected log-likelihood is given by

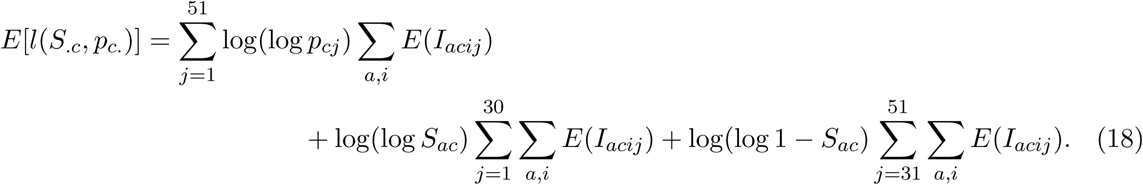

This is then maximized subject to the above constraints by setting

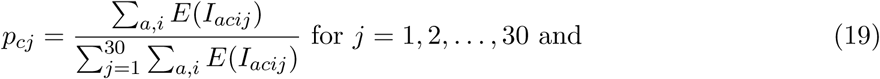

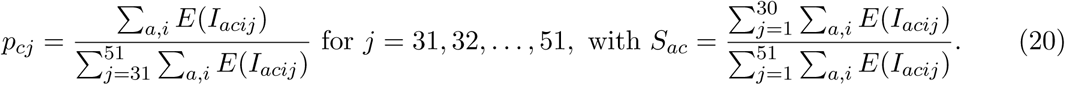

It remains to calculate the expectation terms (of the missing data), over the conditional distribution given current parameter values. To do this note that we may write down

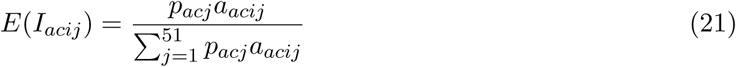

setting the constant *a_acij_* = 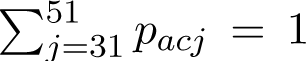

Thus we iteratively calculate the expectations in (4) given current parameters and the resulting updated parameter estimates (3), in our EM algorithm. To initialize this algorithm we set *p_cj_* to be proportional to 0.1, 0.2*, . . . ,* 1.1, 1*, . . .* 0.1 for *j* = 31, 32*, . . . ,* 51 and constant for *j* = 1, 2*, . . . ,* 30. We initialized *S_ac_* to be the fraction of cells in this animal for this chromosome rejecting a chi-squared test (with 1 d.f.) at *p <* 0.05 by using the maximum likelihood ratio varying over *f_j_*, vs. *f* = 1. We terminated the algorithm when the improvement in the log-likelihood was below 0.01.

##### Adjusted fitting with an animal-specific null distribution

Using the results of this fit as a starting point, we now fixed the values of *p_cj_* for *j* = 1, 2*, . . . ,* 30 (i.e. fixed the “shape” of silenced chromosomal copies), but now allow *p_acj_* to be unconstrained for *j* = 31, 32*, . . . ,* 51 save that 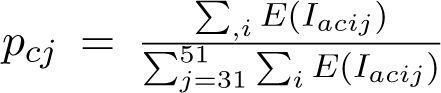 *p_acj_* = 1, i.e. we allow an unconstrained animal-specific null distribution. We also re-estimated *S_ac_* for each animal *a*. Otherwise we proceeded similarly to above, with the resulting EM algorithm having identical updates except *p_cj_* remains fixed for *j* = 1, 2*, . . . ,* 30 while the updated *p_cj_* = 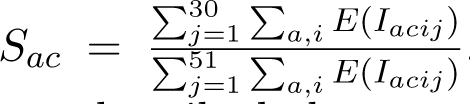 i.e. we now separate by animal, and as before *S_ac_* = 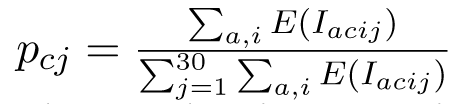. Again, we iterated until convergence of the likelihood with parameters as described above apart from the fixed parameters. We structure the run this way to allow this step to be parallelizable among animals.

Finally we conversely fixed the values of *p_acj_* for *j* = 31, 32*, . . . ,* 51, and fit the values of *p_cj_* for *j* = 1, 2*, . . . ,* 30 using an EM algorithm run as above, apart from the fixed parameters. The resulting final parameter estimates were used for all downstream analyses. Updates are identical to those in (3) for these parameters: *p_cj_* = 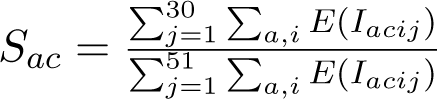 for *j* = 1, 2*, . . . ,* 30, with *S_ac_* = 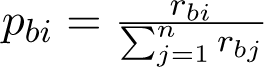 This step runs all animals together but speed is improved by the fixed non-silencing terms. Given these parameters, we also obtained final chromosome- and cell-specific posterior probabilities of silencing under the final fitted model above. Note that the values of *S_ac_* represent the probabilities of silencing by animal and chromosome, and are used downstream as measures of asynapsis by animal and chromosome.

#### 12.3 Estimating probability of autosomal silencing (on any autosome) by animal

We estimated the probability of silencing or asynapsis by animal *S_a_* (on all autosomes jointly, with silencing on autosomes assumed to be occurring independently), as follows:

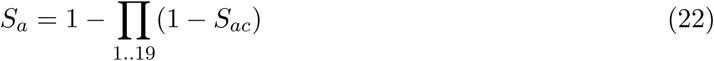

#### 12.4 Predicting silencing from chromosomal CAST content

We modeled the cis impact of CAST sequence on silencing as exponential decay from the level of silencing in the absence of CAST sequence on the chromosome:

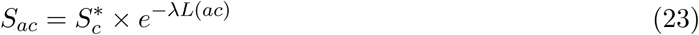

where *S*^∗^ is the average probability of silencing on chromosome *c* in the PWD *×* B6 animals, *λ* is a decay parameter that reflects the genome-wide average effect of CAST sequence on silencing per Mb (or alternatively the impact of restoring symmetric PRDM9 binding per Mb) and *L*(*ac*) is the total length of the CAST background (in Mb) on chromosome *c* in animal *a*.

We fit the single parameter *λ* using estimated asynapsis/silencing probabilities across all autosomes and all animals jointly. For each animal *a*, we first calculated the predicted probability of silencing on each chromosome *c* using the exponential decay model above (12.4) for a chosen value of *λ*. We then calculated *p_a_*(*λ*), the probability that a single cell from animal *a* has at least one silenced chromosome under this model, assuming independence across chromosomes (12.3). We modeled the number of cells with at least one chromosome silenced in each animal as a binomial random variable, with animals treated as independent, so that for animal *a* with *n_a_* total pachytene cells and an observed proportion *S_a_* of cells with silencing, the joint log-likelihood across all animals is:

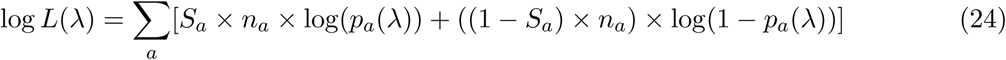

We estimated *λ* by maximizing this log-likelihood using a grid search over the interval [0.01, 1]. The resulting maximum likelihood estimate of *λ* was then used to predict per-chromosome asynapsis/silencing probabilities from *cis* CAST content (i.e., the “genetic prediction” for a chromosome). Animal-level predicted silencing was obtained as before from per-chromosome predictions, treating chromosomes as independent (12.3).

#### 12.5 Calculating Relative Silencing Efficacy

We calculated the total asynapsis proportions (i.e., the sum of observed silencing levels) for only chromosomes whose genotype was PWD *×* B6 throughout (with *<*2Mb of inferred CAST DNA) and which had an estimated probability of silencing of at least 5%, within each animal, and divided by the average of the corresponding sum for the two PWD *×* B6 sterile mice to obtain this quantity.

#### 12.6 Estimating X chromosome silencing

As for autosomes, where we observe some RNA reads even under transcriptional silencing, we defined silencing on the X chromosome as reduction to 75% (or less) of the median proportion of expression from the X in cells from early meiosis (pre-silencing). We then estimated X silencing by animal as the fraction of cells in pachytene with the X chromosome silenced per the above criterion.

### 13 ATAC-seq Fragment size Distributions on the X and Autosomes

We used the distribution of fragment sizes in the genome to examine chromosome-level chromatin properties in individual cells.

#### 13.1 Comparing nucleosome configurations across genomic subsets

To compare the contributions of different chromatin signatures on the X and autosomes, we binned cells ordered by Pseudotime, and summed fragments belonging to a given size category on the X, dividing by the total number of fragments across all size categories in that bin (and calculated also the corresponding value on autosomes). The same approach was used to compare other subsets of the genome (e.g., hotspots): we flagged fragments (using their midpoints) located within *±*300 bp of a PRDM9 hotspot center, as previously defined. The size categories were defined as follows: “sub-nucleosomal fragments” (30-120 bp); “normal nucleosomes” (150-180 bp); and altered nucleosomes (210-290 bp). These size categories were chosen on the basis of observing shifts in the distribution of the X and autosome fragment size distributions through Pseudotime, and checked using a principal component analysis (below).

#### 13.2 Principal component analysis of fragment size distributions

Fragments were grouped by size into bins with 10 bp increments, representing a range of fragment sizes from 30 bp to 500 bp, and were counted by cell and chromosome (including the X chromosome; Y chromosome excluded). Cell-chromosome combinations with fewer than 20 total fragments were filtered out. Fragment counts were normalized by dividing by the total count for each cell-chromosome combination. Principal component analysis was performed on this normalized matrix (cell-chromosome combinations *×* fragment size bins) with centering and scaling, yielding PC loadings on fragment sizes and weights on the X chromosome and autosomes for all cells. The first three PCs explained 15.6%, 4.3%, and 4% of the variance in genome-wide fragment size distributions on chromosomes through meiosis, with fragment size loadings corresponding to nucleosome-free regions, regions with standard nucleosomes, and an altered nucleosome configuration as defined above (see also Supplementary Fig. 11).

#### 13.3 Estimating the mixture proportion of X-like chromatin by chromosome in cells

The X chromosome has a distinct chromatin structure (vs autosomes) in normal Pachytene cells. We modeled the ATAC-seq fragment size distribution on individual chromosomes for each cell in Pachytene as a mixture of autosomal and X-like chromatin. We first calculated for each cell and chromosome the expected (null) relative fragment size distribution on the X vs autosomes using cells from B6 and/or CAST animals, limiting to a window of *±*200 neighboring cells in pseudotime to account for variation in this quantity through time. We calculated the relative proportions of fragment counts in 10 bp fragment size bins (for fragments of length 40-500 bp) on the X vs Autosomes, denoted as *g*, where *g_k_* indicates the X/A ratio for the *k*th size bin.

Under our model, the likelihood of observing a fragment in size bin *k* is simply proportional to (1*−λ*)*×*1+*λ×g_k_*, where *λ* is the mixture proportion of X-like chromatin. We fit *λ* by summing log-likelihoods across all observed fragments (for each cell and for individual chromosomes within each cell) and obtaining the value using a grid search over the interval [0, 1] that maximized the overall likelihood, yielding a per chromosome measure of the presence of X-like chromatin on autosomes in individual cells.

To identify chromosomes with significant X-like chromatin signatures, we tested whether each chromosome showed evidence of X-like chromatin by comparing the best-fitting *λ* value to the null hypothesis (*λ*=0) using a likelihood ratio test. Second, we tested whether the chromosome-specific signal differed from the cell-wide baseline by comparing the chromosome-specific maximum likelihood to the likelihood at the cell’s overall best *λ* value. We considered chromosomes where both tests exceeded a threshold of 5 (p-values *<* 0.0015 under a chi-squared distribution) to have an X-like chromatin proportion significantly different from zero. We excluded cells in the bottom 10% of total ATAC-seq fragment counts. To account for cell-to-cell variation in baseline chromatin, we normalized each chromosome’s *λ* value by subtracting the mean *λ* across all 19 autosomes within that cell. For comparisons across autosomes, we normalized each autosome by subtracting the mean *λ* of the remaining 18 autosomes. Animal-level and chromosome-level estimates were calculated as the mean normalized *λ* values across all cells for each animal-chromosome combination.

### 14 Genome-wide genotype-phenotype associations

Together with our measurements of testes weight and sperm count, we collated the following phenotypes for each of 23 PWD *×* B6 Cast mice: the fraction of pre-division cells in late pachytene (with late pachytene defined as Pseudotime values in the range 100K - 150K, of 203K ranked cells, and pachytene defined to correspond to the interval of X chromosome silencing, cell ranks 30K - 150K in our data), the probability of autosomal silencing by animal (calculated in 12.3), relative silencing efficacy (calculated as described in 12.5) and crossover rates (see below). At 286 loci in the genome with distinct (though often correlated) configurations of local ancestry, we tested for associations between genotype (“B6” or “CAST”) and these phenotypes, using a simple linear model with genotype as the predictor.

We assessed genome-wide significance using a conservative Bonferroni-corrected significance threshold of 0.05*/*286 = 0.00017. We further used a genome-wide permutation approach to calibrate p-values. Specifically, the chosen phenotype was randomly permuted across animals 1000 times and the permuted phenotypes tested for association at each of the 286 loci: the minimum p-value from each GWAS was used to generate an empirical null distribution of minimum p-values given our data. We obtained the rank of the observed p-values in this empirical distribution, as a measure of the probability of recovering the observed p-value by chance given the sample and the phenotype.

### 15 Variant consequences

We downloaded predicted variant effects for all SNPs in the CAST background (with B6 as reference) from the Mouse Genomes Project [9] (VCF file *CAST EiJ.mgp.v5.snps.dbSNP142.vcf.gz* ; Assembly: Grcm38/mm10; SNP RELEASE v5; Annotations from the Ensembl Variant Effect Predictor software; Mouse gene models from Ensembl release 78; further details can be viewed at the Mouse Genomes Project FTP site). We excluded variants with low quality heterozygous calls or those that failed other filters (FORMAT/FI = 0). We extracted variant effects from the ‘CSQ’ field and Ensembl Gene IDs where available. We counted the total number of variants by type and by gene in genic regions: where there were multiple consequences associated with the same variant, the stronger effect was considered (e.g., variants that were missense and impacted splicing were counted as missense). Non synonymous mutations were defined as missense (including missense + other consequences), and nonsense (including stop gained, splice donor variants, and splice acceptor variants, and combinations of these with other consequences). The length of the coding region for each gene was calculated by summing exonic lengths (obtained from *EnsDb.Mmusculus.v79* in *R*) in the longest transcript for each gene.

### 16 Timing and allele-specificity of expression in early Pseudotime windows

We used the inferred gene expression time profiles (in 9.1 above, Supplementary Fig. 1) and selected time profiles corresponding to largely pre-Pachytene expression (i.e., those with greater than a third of their weight in time bins corresponding to leptotene and zygotene). We then identified genes with *>*10% loading on these profiles as our group of early expressed genes likely to be most relevant for (a)synapsis.

We examined the subset of early genes in our locus of interest for evidence of allele-specific expression. Because we specifically wanted to examine the effect of the B6 vs CAST background on gene expression, we did so using our B6 *×* CAST hybrid mice. Having identified reads clearly assigned to each homologous chromosome in these mice (see Section 6.3 above), we counted the number of reads from each background for each gene and the total number of reads from each background genome-wide, and used Fisher’s exact tests to obtain p-values by gene.

### 17 Chromosomal abnormalities in post-division cells

To examine potential chromosomal abnormalities in late meiotic cells, we focused on PWD *×* (B6 *×* CAST) mice (since PWD *×* B6 mice have a negligible number of such cells), using fertile B6 *×* CAST F1 hybrids as controls. We analyzed ATAC-seq fragment counts by chromosome in populations of cells that have undergone one or both meiotic divisions, including cells that predominantly had the features of MI and MII cells, but potentially had defects on a small number of autosomes (which we previously identified as “MI”, “Mainly MI”, “MII”, or “Mainly MII”; see section 8 above), as well as cells that have potentially failed to undergo the meiotic divisions but have overall gene expression consistent with post-division cells (see below). Consistent with a relatively brief period (several hours) between the meiotic divisions, we observed small populations of MI cells (n=440 cells for fertile B6 *×* CAST mice and n=3,306 cells for PWD *×* (B6 *×* CAST) animals; we observed *>*6-fold more cells that have completed Meiosis II, for both backgrounds.

#### 17.1 Classifying post-division cells by sex chromosome content

The first meiotic division normally separates homologous chromosomes and the second sister chromatids, so that secondary spermatocytes are expected to have either XX or YY, and spermatids X or Y. To separate X and Y bearing MI and MII cells, we calculated the fraction of ATAC fragments that mapped to the X and Y chromosomes relative to autosomes (for the Y chromosome we counted all fragments, including those with low mapping quality).

We first defined reference populations of MI and MII cells with at least 1000 autosomal ATAC fragments in the two fertile B6 *×* CAST F1 hybrids, assigning them to one of four groups (X-only, Y-only, both X and Y, and neither sex chromosome) based on whether their log2-transformed X and Y chromosome fractions exceeded manually determined thresholds of -7.5 and -8, respectively (as these thresholds, corresponding to 0.55% and 0.39% of autosomal fragments required to call X and Y presence, cleanly separated the four populations in log2 space; Supplementary Fig. 15). We fit two-dimensional kernel density estimates using the ks R package. Having learned the parameters of the distribution of X and Y proportions in each category of cells from the reference data, for each cell in the full dataset we evaluated its likelihood under each of the four distributions. We estimated group proportions using an expectation-maximization procedure. Starting from equal proportions (= 0.25 each), we iteratively (over 50 iterations) computed posterior probabilities for each cell, then updated group proportions as the average posterior across all cells. These classifications were used to calculate the proportion of cells with abnormal sex chromosome configurations (Supplementary Figure 16, Figs. 6F and 6G).

#### 17.2 Identifying abnormal autosomal configurations

For autosomes, we previously obtained an ATAC-seq based classification of individual chromosomes (for example “Meiosis-I-recombinant”, or “Homozygous deleted”) in the same MI and MII cells as above (see section 8 above). In parallel, we also examined these cells for putative autosomal deletions and duplications using ATAC-fragment proportions by chromosome to supplement the HMM-based chromosomal designations (as there is a huge amount of variability in ATAC-seq read counts by cell, these alone are not reliable indicators of autosomal duplications in particular). For each group of cells (MI and MII), we calculated the mean proportion of ATAC-seq reads by chromosome, and for chromosomes in individual cells, we calculated the fold enrichment over this value.

We then defined chromosomes as (a) having a homozygous deletion if they were called as “Homozygous deleted” based on the HMM approach above (which requires 90% of the chromosome inferred to be in the 0 : 0 state) and they additionally have *<* 0.1-fold the mean proportion of ATAC fragments for the chromosome in the same cell type, or (b) being in a “low-copy” state if they have *<* 0.6-fold the mean proportion of ATAC fragments for the chromosome in the same cell type. We defined chromosomes as being in a “high-copy” state and/or being duplicated if they have *>* 1.4-fold the mean value for that chromosome and cell type, and meet the following criteria: for MI cells, the chromosome must be called as a Heterozygote (i.e., a chromosomal call that is “Heterozygous”, “Heterozygous (unbalanced)”, or “Heterozygous (aberrant)”); for MII cells, both Heterozygote and MI-recombinant states (i.e., “Heterozygous”, “Heterozygous (unbalanced)”, “Heterozygous (aberrant)”, “MI recombinant”, “Aberrant recombinant”, “High copy recombinant”) were allowed. These categories are defined in section 8, and reflect the possible configurations of high copy products of improper segregation at the first and second meiotic divisions in our system (see also Fig. 1E).

#### 17.3 Examining the distribution of MI and MII cells in pseudotime

We started with pseudotime ranks obtained for each cell based on gene expression patterns (see Section 9) and the ATAC-based cell type classifications described above. Examining pseudotime values for MI and MII cells, 95% of MI cells were ranked *>* 150, 000 (of 203, 869 cells) in our animals, and 95% of MII cells are ranked *>* 157, 000, validating our inference of these as late meiotic cells using gene expression data (see also Fig. 6E).

Of normal (or mainly normal) cells that passed the mid-pachytene checkpoint (defined as pseudotime interval containing 90% of all cells in sterile PWD *×* B6 mice, corresponding to cell rank 61, 090), 1.6% ranked *>* 150, 000 in B6 *×* CAST and 5.8% in PWD *×* (B6 *×* CAST) animals (OR = 3.8, *p <* 10*^−^*^16^), indicating an enrichment of undivided cells in PWD *×* (B6 *×* CAST) animals.

In turn, B6 *×* CAST animals had MI cells occurring in a narrow time range: only 5% of MI cells exceed pseudotime values of 157,000, with MII cells dominating beyond this time. However, in PWD *×* (B6 *×* CAST) animals, while the lower bound of such cells was near-identical, we observed *>* 63% of cells at times above this (late to early OR= 33.3, using Fisher’s exact test: *p <* 10*^−^*^16^), consistent with these cells, with expression patterns similar to healthy cells that have completed the second division, but DNA patterns from the ATAC-seq data showing only Meiosis I, not completing the second division and accumulating as a population of aberrant/arrested cells.

#### 17.4 Investigating errors associated with likely abnormal MI cells

Stratifying MI cells into “early” and “late” populations using pseudotime ranks (150,000 - 157,000 and *>*157,000, respectively, see also Fig 6E), for each identified type of abnormal sex chromosome and autosomal configuration identified above, we tested for enrichment in early vs. late MI populations by a Fisher’s exact test using a binary indicator of that chromosome or cell type (Supplementary Table 6). We also calculated the proportion of MI cells with sex chromosome deletions and duplications (obtained as above) that showed evidence of arrest/failure to undergo the second division (based on belonging to the “late” MI population).

#### 17.5 Investigating sources of deletions and duplications in MI and MII cell divisions

Given the earlier defects of synapsis and transcriptional silencing downstream of PRDM9-binding asymmetry in many PWD *×* (B6 *×* CAST) animals, we investigated whether these processes further impact chromosomal segregation at the two meiotic divisions. To this end, we tested whether chromosome copy number changes (deletions or duplications) in MI or MII cells were significantly associated with either asynapsis levels of the respective chromosome in that PWD *×* (B6 *×* CAST) animal (i.e., associated with cis impacts of asynapsis, which varies by chromosome and animal in our experiment) or the estimated animal-level asynapsis/silencing (associated with trans impacts of asynapsis, i.e., not necessarily arising from the same chromosome).

As a test statistic, for the *cis* testing we used the fraction of chromosomes of the relevant type (e.g, high or low copy number) possessing very little (*<* 5 Mb) CAST DNA, so that issues induced by PRDM9-induced asynapsis should have higher-than-expected values for this statistic. For the *trans* testing we used the correlation of the 0-1 label of a chromosome being a particular type with the corresponding animal-wide asynapsis number (treating both as vectors over all cells and chromosomes examined), expected to be significantly positive if this type is associated with overall asynapsis. To perform the test of significance of these statistics, we permutation-tested a null of no association by keeping the chromosome-cell data constant, including which animal these data correspond to, but then performed 1000 bootstraps by permuting the corresponding animal labels (separately for each chromosome for testing the *cis* chromosome-specific measures, or only overall for testing the *trans* measure of overall animal-wide synapsis level). This retains variable data amounts (or even quality) among animals.

For both MI and MII cells, we used this approach to test for enrichment in B6-only autosomes and correlation with animal wide asynapsis rates, of likely low copy or likely high copy number of autosomes, as defined above, and correlation with animal wide asynapsis rates for either no X or Y chromosome or both X and Y chromosomes being present (Supplementary Table 7).

#### 17.6 Investigating sources of MII chromosome deletions (sex chromosomes and autosomes)

For each animal (filtering animals with less than 500 cells), we counted MII cells with deletions of the X and Y (i.e., instances where both sex chromosomes were absent), and/or deletion of at least one autosome, with the deletion calls made as described above.

We first tested whether deletions of X and Y, and of at least 1 autosome, occur in the same animals: rates of deletion in chromosomes 1-10 vs 11-19 were highly correlated (*r* = 0.96*, p* = 1.6 *×* 10*^−^*^7^), and rates of deletion of autosomes and the X were also highly correlated (*r* = 0.83*, p* = 3.6 *×* 10*^−^*^4^). Given that animals differ in unsynapsed autosomes, these imply a shared *trans* driver of MII deletions.

We fit an initial model using the average autosomal probability of an inferred deletion as the target (response) variable and the animal-wide asynapsis probability (Supplementary Table 8) as predictor, finding that although the association is significant, the resulting fitted values showed notable lower correlation with the observed values than the correlations across chromosomes and suggesting the model might be improved upon. We therefore considered additional models allowing different weights for asynapsis on specific chromosomes (estimated using their inferred silencing probability, see above). Starting from the empty model we step-wise successively added that chromosome into the model whose p-value is minimal, while this p-value was below the Bonferroni-corrected significance threshold (*p <* 0.05*/*19) for testing all autosomes. This resulted in a minimal significant model including chromosomes 16 and 19 only (Supplementary Table 8, model II). We checked additional testing of these chromosomes plus the animal-wide asynapsis probability (Supplementary Table 8, model III), which did not improve the fit and revealed the animal-wide asynapsis probability to be no longer predictive once chromosomes 16 and 19 were included, while these chromosomes remained highly significant.

Finally, we *fixed* the coefficients for chromosomes 16 and 19 to those inferred in model II, to generate a single fixed predictor, and changed the response variable to the independently estimated probability of no X or Y chromosome being present in MII. Then, we repeated the testing of models I-III using this new response, producing 3 model fits (IV, V, VI; Supplementary Table 8), which showed analogous properties to the autosomes: significant association with the model based on chromosomes 16 and 19, which explains the correlation with overall fertility; overall *r*^2^ values were slightly lower for the X/Y response but this might be explained by reduced data compared to averaging all 19 autosomes. Although given our modest sample size other explanatory models are possible (for example other chromosomes than 16 and 19 might be involved), nonetheless these analyses provide strong evidence that genetic factors varying by animal, and well predicted by asynapsis (much earlier in meiosis), drive MII deletions in *trans*.

### 18 Crossovers

#### 18.1 Calling crossovers in post-division cells

To call crossovers in single cells, we used post-division cells that are confidently classified as “MI” or “MII” (1,219 and 19,717 cells, respectively, in PWD × (B6 × CAST) and B6 × CAST mice based on their genome-wide state proportions (see section 8 above). As input we used the homolog-specific counts in 50kb bins along the genome for individual cells that were used above for cell classification. We aggregated counts in 10 50kb bins to get homolog specific counts per 500kb.

To identify sites of crossovers in haploid (MII) cells, we used a two state HMM to classify genomic bins by the homolog of origin of the reads. As in Section 7 above, given a sequence of *K* genomic bins with *j* ∈ {1 = “PWD”, 2 = “non-PWD”} possible states, observed counts (*n*_PWD_(*k*)*, n*_non-PWD_(*k*)) for the *k*th bin, and an emission parameter *p_i_*for state *i*, which is the probability of drawing a PWD read when in the *i*th state (*p*_1_ = 0.9*, p*_2_ = 0.1), we calculated the likelihood of the observed counts as an emission probability with binomial sampling for state *i* as:

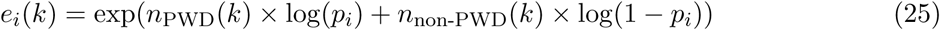

Chromosomes in MI cells are expected to be homozygous for one of the two homologs at the centromeric end (with at most 2 heterozygous starts allowed per MI cell by definition), and to switch to a heterozygous state at the point of a crossover. We modified the above HMM to classify bins as homozygous or heterozygous. With an expected equal contribution (50/50) from both homologs in heterozygous regions, the emission likelihood for the heterozygous state was calculated as:

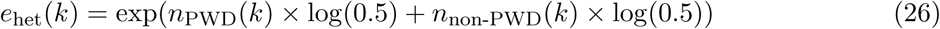

For the homozygous state, we used data from the first (centromeric) 5 Mb of each chromosome to choose from the two possible homologs at the centromeric end of each chromosome (PWD or non-PWD) in every cell and then calculated binomial likelihoods as:

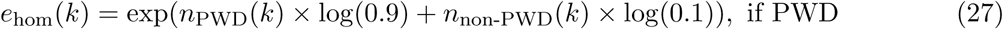

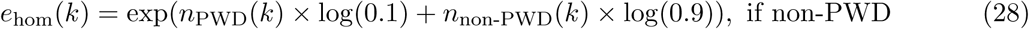

As described in more detail above, we implemented the Forward Backward algorithm (initialized with both states equally likely once per chromosome per cell) with fixed transition probabilities of 0.005 between states for MI cells and 0.0025 for MII cells (i.e., as described above, one switch expected per 100 Mb, and half that rate expected in MII cells). We assigned the maximum a posteriori state at each position and recorded the positions of switches between the two states on each chromosome, by cell.

We stringently filtered cells (removing 97 cells in total) using the following criteria: more than 30 crossover calls or a heterozygous call at the centromere on more than one chromosome if MI and more than 20 crossover calls if MII; at least one chromosome had *>*4 crossovers called, as we expect at least some of these to be spurious calls. We excluded a small number of (*<*500) apparent recombination events less than one Mb apart, for the same reason. We note that we obtained very similar results (including for the relationship with asynapsis, see below) using the genome-wide cell states in 50 kb bins from our cell classification algorithm (described above), where heterozygous and homozygous segments by homolog (and therefore crossovers) can be identified directly and extremely quickly; however it required filtering a much larger number of cells to remove spurious calls because of more limited information in 50 kb bins.

We summarized crossover events (245, 368 in total across 24 animals, including two B6 *×* CAST hybrids) by chromosome and cell for MI and MII cells, and checked that MII crossover events were on average approximately half as frequent as those in MI cells (Supplementary Fig. 16). 97% of chromosomes in MI cells had at least one detectable crossover in our sample, and 86% possessed exactly 1 crossover (Supplementary Fig. 16). In MII cells across all mice, 46% of chromosomes were non-recombinants, and 48% showed exactly 1 crossover (Supplementary Fig. 16). We assigned each crossover an ancestry background (”PWD/B6” or PWD/CAST”) using the local ancestry segments obtained above. We aggregated crossovers (in MII cells) by animal and chromosome (binned by position) in B6 x CAST animals with the human *Prdm9* allele, and in PWD *×* (B6 *×* CAST) mice; the two B6 x CAST animals had highly correlated (95%, *p <* 10*^−^*^10^) crossover rates in 5 Mb bins (see also Supplementary Fig. 17).

Because we have few MI cells per animal (and as we show, many of these cells likely undergo arrest), we estimated crossover rates by chromosome (and over all chromosomes) by averaging all MII cells by animal and checked that these were consistent with reported ranges [8]; we tested these rates for association with varying local ancestry along the genome in PWD *×* (B6 *×* CAST) mice (as described in Section 14); we found no genome-wide significant loci.

#### 18.2 Investigating crossover positioning across mice

We filtered the list of 207, 475 crossovers in 22 PWD *×* (B6 *×* CAST) animals to those cases (the majority) where that chromosome contained exactly one crossover event, and studied the relative odds an event occurs in B6 or CAST DNA segments. We split each chromosome into 2Mb bins, allowing the recombination rate to vary across bins and to differ between the CAST and B6 backgrounds (events can also be initiated on the PWD background, but this is shared between all animals, so this does not impact our modelling).

For each animal, we summarised their data for a given chromosome as a vector of total recombination counts in each 2Mb bin, and we then modelled the probability the event occurs in each segment as follows.

First, if there are *n* 2Mb segments then we parameterise rates *r_b_*_1_*, r_b_*_2_*, · · · , r_bn_* of recombination in the PWDxB6 regions and *r_c_*_1_*, r_c_*_2_*, · · · , r_cn_* in the PWDxCAST regions respectively. Then the probability an event occurs in segment *i* in a 100% PWD *×* B6 chromosome is simply *p_bi_* = 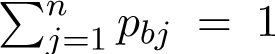 and similarly for analogously defined probabilities *p_ci_* in a 100% PWDxCAST chromosome.

For an arbitrary chromosome, define *I_i_* = 1 if segment *i* is CAST in a particular animal and 0 if segment *i* is B6. In practice, we identified values of these indicator variables using informative SNPs to define local ancestry (see above), taking that background covering *>*50% of each interval near breakpoints. Now, we introduce a parameter *λ* which is the relative odds of a break occurring in CAST DNA, so that the probability of a break occurring in segment *i* is proportional to (1 *− I_i_*)*p_bi_* + *λI_i_p_ci_* and so precisely equal to:

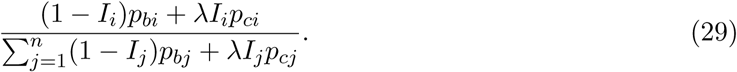

Note this results now in different rates for the same background, depending on the makeup of other parts of the chromosome. Our focus of interest is in the value of *λ*, which measures the relative “heat” of CAST segments compared to B6 segments, in cases where part of the chromosome is from each background. If *λ >* 1 then CAST segments are more crossover-prone, while if *λ <* 1 the opposite occurs. We must also infer the nuisance parameters *p_bi_, p_ci_*. We assume these are shared across animals (i.e. there is an underlying shared background-specific heat), while *λ* is either also shared across animals (but varies by chromosome), or, for later analysis, is allowed to vary across animals also.

We fit the above model via maximum-likelihood using Newton-Raphson and used bootstrapping of events to estimate empirical CI’s for lambda. To do this, we note that if we observe *k_i_* recombination events in interval *i*, the log-likelihood we must maximise is given by

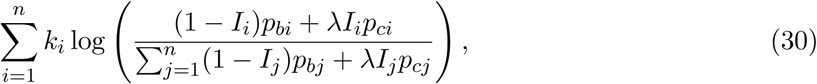

which is a differentiable function of the parameters. Given data for multiple animals we simply add up the likelihood contribution of each animal (and it remains differentiable).

We used Newton-Raphson iteration to identify the maximum likelihood values of the above parameters, initialising *p_bi_* simply to be the average number of events occurring in segment *i*, across animals where *I_i_* = 0 and then normalising these values to sum to 1, *p_ci_*simply to be the average number of events occurring in segment *i* across animals where *I_i_* = 1 and then normalising these values to sum to 1, and then setting *λ* = 1. Noting that our parameters have the constraint 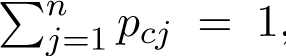 and similarly 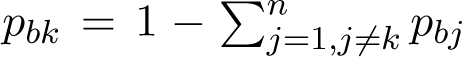, this constraint is accounted for by choosing any k and then setting, before differentiating, *pbk* = 1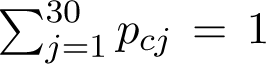 *pbj* , similarly for any l and *pcl*, and applying Newton-Raphson on the remaining 2*n −* 1 parameters. Rather than fixing *k, l* we instead at each step apply Newton-Raphson for successive possible pairs of values, trying all possible pairs if necessary, and starting with those values of *k, l* corresponding to the largest current probabilities *p_bk_* and *p_cl_*, then in decreasing order. We accept any update increasing the log-likelihood while remaining feasible (all *p_bj_, p_cj_ ≥* 0; if no feasible update only *λ* updated), and iterate 100 times (in all cases this was sufficient for convergence). Our testing demonstrated this procedure to converge and increase the log-likelihood with each iteration as expected. For bootstrapping, we randomly sampled recombination events with replacement and repeated the entire procedure; we used 1000 bootstrap samples.

After fitting an initial model with fixed chromosome-wide *λ* values (i.e., one *λ* value per chromosome, shared across animals), and using these to investigate chromosomal differences in crossover enrichment in the CAST background, we further fit the model as above, but now using animal-specific *λ* values for each chromosome. Rather than update all animals simultaneously, we fixed the values of *p_bj_, p_cj_* at their estimates from the animal-wide analysis and simply used the events in each particular animal (on a given chromosome) to estimate *λ*, using Newton-Raphson to identify the maximum likelihood value for each animal-chromosome combination. For animal-chromosome combinations where the fraction of CAST ancestry fell between 5% and 95%, we performed likelihood ratio tests comparing the maximized log-likelihood under the estimated values of *λ* to the log-likelihood when *λ* was constrained to equal 1 (i.e., no enrichment in the CAST background), calculating p-values using the chi-squared distribution.

